# Immune-parenchymal multicellular niches are shared across distinct thyroid autoimmune diseases

**DOI:** 10.1101/2025.03.30.646176

**Authors:** Michelle Rengarajan, Rachelly Normand, Hoang Tran, Linda T. Nieman, Benjamin Arnold, Michael Calcaterra, Katherine H. Xu, Peter Richieri, Enrique E. Rodriguez, Kamil Slowikowski, Yuhui Song, Alice Tirard, Antonia E. Stephen, Peter M. Sadow, Sareh Parangi, Gilbert H. Daniels, Andrew D. Luster, Alexandra-Chloé Villani

**Affiliations:** Department of Medicine, Thyroid Unit, Massachusetts General Hospital, Boston, Massachusetts (MA), USA; Center for Immunology and Inflammatory Diseases, Department of Medicine, Massachusetts General Hospital, Boston, MA, USA; Krantz Family Center for Cancer Research, Massachusetts General Hospital, Boston, MA, USA; Broad Institute of Massachusetts Institute of Technology and Harvard, Cambridge, MA, USA; Harvard Medical School, Boston, MA, USA; Department of Surgery, Massachusetts General Hospital, Boston, MA, USA; Departments of Pathology, Massachusetts General Hospital and Harvard Medical School, Boston, MA, USA; Division of Rheumatology Allergy and Immunology, Massachusetts General Hospital, Boston, MA, USA

**Keywords:** Autoimmunity, Thyroid, Single-cell multiomics, Translational Immunology, Immune hubs Short title: Thyroid Immune-Parenchymal Niches

## Abstract

Thyroid hormone, produced in the thyroid gland, regulates metabolism, development, and cardiac function. The thyroid is susceptible to autoimmune attack by both cellular and humoral immunity exemplified by Hashimoto’s thyroiditis (HT) and Graves’ Disease (GD), respectively. In HT, immune-mediated destruction impairs thyroid hormone production, while in GD, stimulating autoantibodies promote over-production. Here, we generated a multi-modal atlas of 604,076 human thyroid and blood cells from HT, GD, and control patients. We found that, despite markedly different clinical presentations and distinct antigenic triggers, HT and GD exhibit convergent cellular dynamics resulting in a shared continuum of immune infiltration. Along this continuum, a key feature is a thyrocyte niche containing *CD8*^+^ T cells that may segregate pathogenic T cells from regions with preserved thyroid hormone production. These findings of a shared disease continuum characterized by spatially defined immune niches provide a new framework for understanding tissue homeostasis in human autoimmune disease.

## INTRODUCTION

Autoimmune disease, in which the body’s immune system inappropriately attacks its own tissues, affects up to 10% of the global population and is a significant driver of healthcare utilization and cost^1,2^. A substantial fraction of organ-specific autoimmunity, in which immune responses are directed against antigens confined to a specific tissue, target components of the hormone-producing apparatus of endocrine glands, such as pancreatic *β*-cells in type 1 diabetes or thyroid-hormone producing epithelial cells (thyrocytes) in the thyroid. These disorders are characterized by the presence of self-reactive T cells that infiltrate target tissues, and may lead to destruction of hormone-producing cells and, ultimately, loss of tissue function.

The thyroid gland serves as an unparalleled model to investigate the progression of tissue-specific autoimmune disease in humans. Thyroid autoimmunity is not treated with immune-modulatory therapy; thus, tissue dynamics reflect the natural course of disease in humans. Chronic lymphocytic (Hashimoto’s) thyroiditis (HT), one of the most prevalent autoimmune disorders, demonstrates that immune infiltration of a target tissue does not invariably culminate in loss of thyroid function. Strikingly, over 15% of women in the United States have circulating antibodies targeting thyroid peroxidase^3^, which are diagnostic of inappropriate thyroid immune infiltration^4^; less than one-third of antibody-positive individuals have loss of tissue function requiring hormone replacement therapy^5,6^. Based on these observations, we postulated that robust regulatory processes within the thyroid could contain immune infiltration and preserve thyroid function. Therefore, we performed a comprehensive examination of the dynamic landscape of autoimmune thyroid disease directly in humans, which we believe could reveal common features of human tissue resilience across organ-specific autoimmunity.

In this study, we constructed a high-resolution multimodal single-cell atlas of the human thyroid across a spectrum of autoimmune progression, including patients with different degrees of tissue immune infiltration and with and without loss of function. Importantly, thyroid function, specifically the production of thyroid hormone, is readily quantifiable with a single blood test measuring levels of thyroid stimulating hormone (TSH); this enables direct analysis of the relationship between immune infiltration and organ function in patients who are not on hormone replacement.

We focus on two common and clinically distinct forms of autoimmune thyroid disease: Hashimoto’s thyroiditis (HT) and Graves disease (GD). HT is defined by the presence of thyroid-infiltrating lymphocytes that target hormone-producing thyrocytes leading to gradual tissue destruction and, in some cases, loss of function^6^. In contrast, GD is defined by the presence of stimulatory antibodies against the TSH receptor, leading to thyroid hyperactivity and excessive hormone production^7^. By integrating data from HT and GD, as well as from individuals without thyroid autoimmunity, we were able to characterize the dynamic cellular landscape of organ-specific autoimmunity in human tissue from minimal immune infiltration to advanced thyroid dysfunction. Our findings not only shed light on the pathogenesis of thyroid autoimmunity but also provide insights into broader principles of tissue tolerance that may be applicable to other organ-specific autoimmune diseases.

## RESULTS

### A comprehensive multimodal cellular atlas of the human thyroid in health and disease

To determine how the cellular landscape of human tissue evolves with progressive autoimmune infiltration, we constructed a multi-modal single cell atlas incorporating data from thyroid tissue from 23 adult patients, including 8 patients with Hashimoto’s thyroiditis (HT), 9 patients with Graves disease (GD), and 6 gender-matched controls; collectively this cohort spans the full spectrum of tissue immune infiltration and remodeling (Figure 1A-D, Supplementary Figure 1A-C, Supplementary Table 1, 2). Tissue was obtained from patients undergoing thyroid surgery for a nodule (HT, control specimens) or as a therapeutic intervention for GD; control specimens were obtained from patients without autoimmune thyroid disease (see Methods). In addition, we collected paired blood from each patient at the time of surgery.

**Figure 1:**
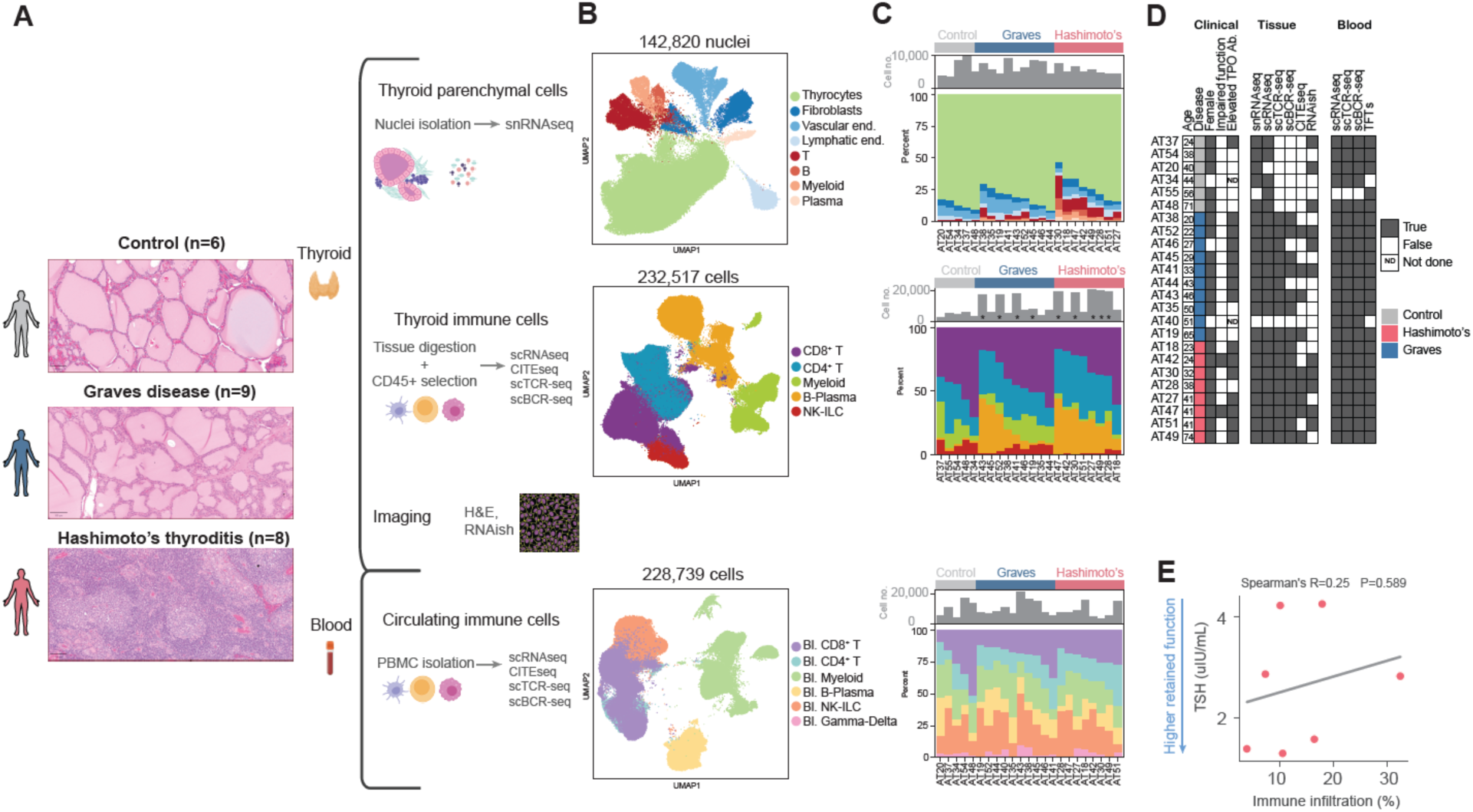
A multi-omic single cell atlas of the human thyroid. **A.** Experimental design. Thyroid tissue and paired blood samples were collected from patients without thyroid autoimmunity (control, n=6), with Hashimoto’s thyroiditis (HT, n=9) and with Graves disease (GD, n=8). Representative H&E staining was performed on paraffin-embedded thyroid tissue (left). Thyroid samples were profiled by snRNAseq (top). CD45^+^ (immune) cells from the thyroid were enriched by FACS and profiled by scRNAseq with paired CITE-seq and TCR/BCR profiling (middle). Peripheral blood mononuclear cells (PBMCs) were profiled by scRNAseq with paired CITE-seq and TCR/BCR profiling (bottom). **B**. UMAP embedding of thyroid nuclei (top), thyroid-associated immune cells (middle) and PBMCs (bottom). Colors indicate major cell lineages. **C.** Composition of thyroid cell lineages within each sample profiled by snRNAseq (top), thyroid immune cells profiled by scRNAseq (middle) and PBMCs (bottom). Each bar represents a single sample. Gray bar plots indicate the number of nuclei or cells profiled per sample. Colors indicate major cell lineages. * indicates samples for which antibody-based cell hashing was employed. **D.** Summary of patient cohort, including measurements collected for each patient. **E.** Relationship between immune infiltration (frequency of thyroid-associated immune cell lineages as determined by snRNAseq, x-axis) and thyroid function (serum TSH level, y-axis) in the 7 HT patients who were not on hormone replacement. The line represents linear regression of the data.

We comprehensively profiled all parenchymal, stromal, and immune cells in tissue with paired blood using single-nuclei RNA sequencing snRNAseq) and single-cell RNA sequencing (scRNAseq) with parallel measurement of surface protein using cellular indexing of transcriptomes and epitopes by sequencing (CITE-seq)^8^ with paired single cell T cell receptor (TCR) and B cell receptor (BCR) sequencing (scTCR-seq, scBCR-seq) (Figure 1A-C). Figure 1D describes the data types generated for each tissue and blood specimen. To maximize recovery of thyroid epithelial cells (thyrocytes) and other parenchymal cells, we compared snRNAseq profiling with scRNAseq of dissociated tissues generated from the same thyroid specimens. snRNAseq was superior in capturing the full-breadth of expected cell types and was thus performed for all samples to determine the native proportion of major thyroid cell lineages and define the parenchymal and stromal cell populations in the human thyroid (Supplementary Figure 1D-G).

To enable deeper multi-modal characterization of thyroid-infiltrating immune cells, we isolated CD45^+^ cells from thyroid tissue from the same patient samples by FACS (Supplementary Figure 1H) for paired CITE-seq, scTCR-seq and scBCR-seq. The thyroid is a highly vascular tissue. Therefore, immune cells obtained from direct dissociation of thyroid tissue, which we term thyroid-associated immune cells, could include: (1) CD45^+^ cells that infiltrated the tissue parenchyma (referred to as thyroid-infiltrating immune cells) or (2) circulating CD45^+^ cells in the lumen of thyroid blood vessels. CD45^+^ cells isolated from samples without significant immune infiltration are likely primarily circulating cells. In contrast, CD45^+^ cells isolated from thyroid samples with high immune infiltration, more likely represent true tissue-infiltrating cells.

From thyroid tissue, we collectively obtained 142,820 high-quality single nuclei and 232,517 single immune cells (CD45^+^) from 23 samples, along with scTCR-seq data from 84,573 single T cells and scBCR-seq data from 21,879 single B cells. CITE-seq thyroid data was generated for 148,792 immune cells across 9 samples. From peripheral blood mononuclear cells (PBMCs), scRNAseq and CITE-seq data was generated from 228,739 cells, along with scTCR-seq data from 18,888 single T cells, and scBCR-seq data from 10,644 single B cells. We additionally performed imaging with RNA *in situ* hybridization (RNAish) on 11 patients for which samples were available (Figure 1B-C; Supplementary Table 2).

We first performed iterative clustering and *post hoc* annotation at the cell lineage level to define large-scale changes in the cellular landscape of the human thyroid in health and disease. SnRNAseq differential abundance analysis revealed a significant increase in all immune cell lineages in tissue for HT compared to control (Figure 1C, top panel, Supplementary Table 3): myeloid (P=0.004), T cells (P=0.004), B cells (P=0.008) and plasma cells (P=0.004)) while fewer thyrocytes in HT relative to control (P=0.017) were observed. There were no significant differences in abundance of blood lineages across disease (Supplementary Table 3), emphasizing the need to profile tissue directly when studying tissue-specific autoimmunity.

Although HT samples showed significantly higher abundance of immune cell lineages in thyroid than control, we noted considerable variability in the degree of immune infiltration across individual samples (compare, e.g., AT30 to AT27 in Figure 1C, top panel). We therefore hypothesized that immune infiltration alone could drive loss of hormone production (i.e. loss of thyroid function) in patients with HT. Thyroid stimulating hormone (TSH) is made by the pituitary gland to stimulate production of thyroid hormone. It is inversely proportional to thyroid hormone production and is the most sensitive clinical test of thyroid function^9,10^. To test our hypothesis, we measured TSH levels in paired plasma samples obtained concurrently with surgical tissue in patients with HT who were not on hormone replacement. A HT patient on hormone replacement and GD patients were excluded from this analysis as their TSH levels do not reflect native thyroid function. We found no relationship between the degree of immune infiltration and TSH level (Spearman’s R=0.25, P=0.589, Figure 1E, Supplementary Tables 1, 4) demonstrating that immune infiltration was not sufficient to determine loss of thyroid function.

### An interferon-responsive thyrocyte population is a hallmark of autoimmune thyroid disease

We then assessed whether immune infiltration could instead induce changes in parenchymal cell states that could, in turn, modulate tissue function. First, we focused on defining the thyrocyte cellular landscape as these constitute the predominant cell lineage in the human thyroid across all subjects (Figure 1C, top panel). We sub-clustered 103,681 thyrocytes and identified 7 transcriptionally distinct cell subsets (Figure 2A). Among these, subset 1.E most closely captured key transcriptional features of classical thyrocytes – including high expression of genes associated with hormone production (*TPO*, *TG*, *TSHR*) and thyrocyte identity (*PAX8*) – but minimal expression of markers of activation, inflammation, and stress response that characterized the other thyrocyte populations (Figure 2A, Supplemental Figure 2A). We therefore refer to this population as 1.E_classical.

**Figure 2:**
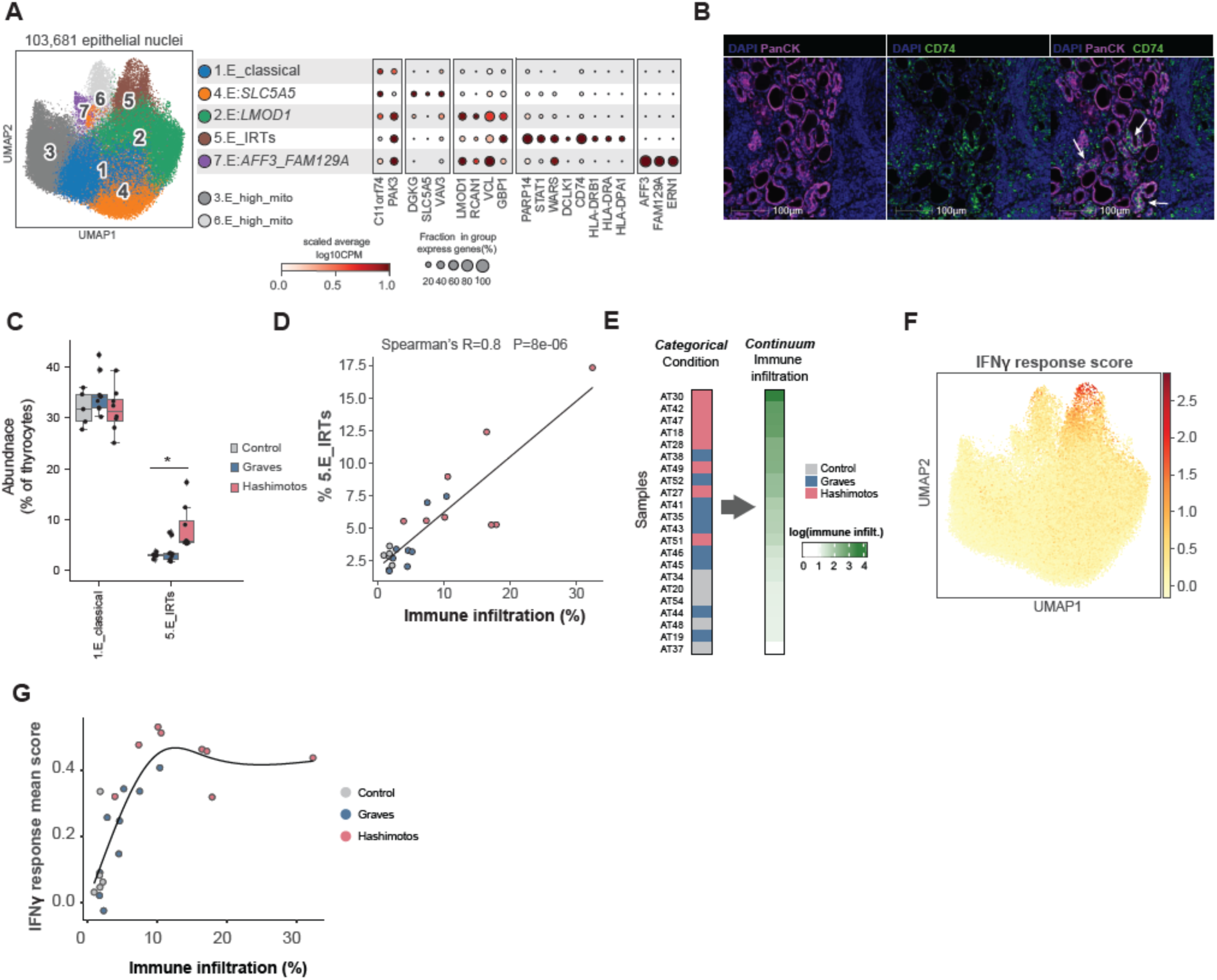
An interferon-γ-responsive thyrocyte population (IRTs) is strongly correlated with immune infiltration. **A.** UMAP embedding of thyrocytes from snRNAseq on thyroid tissue (left). Dot plot displays marker genes for each population (right). Dot size represents the percent of cells in the population with non-zero expression of a given gene and color indicates scaled expression of each gene. **B.** Staining of thyroid tissue from a patient with HT for DAPI (blue), panCK (magenta), and *CD74* (green). White arrows indicate IRTs, which stain for both *CD74* and panCK. **C.** Boxplot illustrates differential abundance analysis comparing the frequencies of 1.E_classical and 5.E_IRTs among thyroid epithelial cells across control, HT and GD. **D.** Relationship between immune infiltration (x-axis) and frequency of 5.E_IRTs among all epithelial cells (y-axis), both determined by snRNAseq. The line represents linear regression of the data. **E.** Categorial modeling of the patient cohort based on each patient’s disease condition (left) versus modeling the cohort as a continuum, based on thyroid immune infiltration (right). **F.** UMAP embedding of thyrocytes colored by mean Z-score of IFN-**γ** response score. **G.** Mean IFN-**γ** response Z-score in 5.E_IRTs per patient colored by condition. The line represents a generalized additive model of the data. P-value: * (<0.05)

A striking population, subset 5.E_IRT, showed up-regulation of interferon-responsive genes (*PARP14*, *STAT1*, *WARS*) and ectopic expression of multiple components of the major histocompatibility complex class II (MHCII) antigen presentation machinery (*CD74*, *HLA-DRB1*, *HLA-DRA*, *HLA-DPA1*) (Figure 2A). MHCII is constitutively expressed on professional antigen-presenting cells, such as B cells and myeloid cells, which present antigen to CD4^+^ T cells^11^. During inflammation, MHCII is expressed in epithelial cells, fibroblasts, and endothelial cells^12^. Notably, 5.E_IRT did not show expression of other markers of professional antigen-presenting cells (Supplementary Figure 2B), arguing that 5.E_IRT cells are indeed thyrocytes that have up-regulated MHCII expression. To orthogonally validate this result, we examined thyroid specimens from an HT patient by RNAish and identified thyrocytes (panCK^+^) in thyroid follicles that clearly expressed *CD74*, a component of MHCII (Figure 2B). 5.E_IRT was specifically enriched in specimens from HT patients (HT vs. Control *P*=0.011, HT vs. GD *P*=0.07; Figure 2C, Supplementary Figure 2C, Supplementary table 4). Strikingly, even in HT, on average 31.3% of thyrocytes still maintain the 1.E_classical cell state even in the context of a high degree of immune infiltration. Altogether, our data show the emergence of a disease-specific thyrocyte population in HT.

We next leveraged the heterogeneity in immune infiltration across our patient population (Figure 1C), and found that the frequency of 5.E_IRT thyrocytes was strongly correlated with immune infiltration across all disease states (Figure 2D, Spearman’s R = 0.8, *P* = 8 x 10^6^, Supplementary Table 4); this was not seen for any other epithelial population (Supplementary Figure 2D). These results further indicate that thyroid immune infiltration exists as a continuum from minimal to high infiltration, without a clear delineation between GD and HT. Indeed, when we modeled patient samples along a continuum of immune infiltration, we observed mixing of patient conditions across this spectrum (Figure 2E). We therefore hypothesized that analyzing our data along the immune infiltration continuum would more accurately represent the underlying biological processes than categorial disease classifications.

To understand if 5.E_IRT thyrocytes have a similar expression profile across patients we constructed an IFN-**γ** response score comprising 53 genes, including genes that were members of the Hallmark gene set for IFN-**γ** response and were identified in a study examining thyrocyte specific response to IFN-**γ** *in vitro*^13^ (Supplementary Table 5). The IFN-**γ** response signature was strongly and selectively expressed in 5.E_IRT cells (Figure 2F, Supplementary Figure 2E). While 1.E_classical thyrocytes showed a subtle response to IFN-**γ** in HT, this was minimal, and 1.E_classical thyrocytes largely maintain their classical identity, even in HT. It is apparent that the expression of this gene set that is central to 5.E_IRTs identity forms a gradient from control to HT, with GD patients having intermediate values (Figure 2G) showing once more that the patients are better modeled along a continuum. Given this population-specific response to IFN-**γ**, we term 5.E_IRT cells IFN-**γ**-responsive thyrocytes, or IRTs.

### GZMK^+^ and exhausted tissue resident-like CD8^+^ T cells may drive IFN-γ-induced tissue remodeling in thyroid during progressive immune infiltration

To identify which thyroid-associated immune cells could induce IRTs, we examined expression of *IFNG* among all immune cell lineages in our scRNAseq dataset. *IFNG* expression was highest in CD8^+^ T cells and was largely absent from other cell types except in NK cells (Figure 3A). Notably, *IFNG* expression in NK cells was restricted to a CD16^hi^CD56^lo^ NK population that was depleted in HT and GD samples (Supplementary Figure 3A-E) and resembles the predominant circulating NK population^14^. We therefore turned our attention to thyroid-associated CD8^+^ T cells.

**Figure 3:**
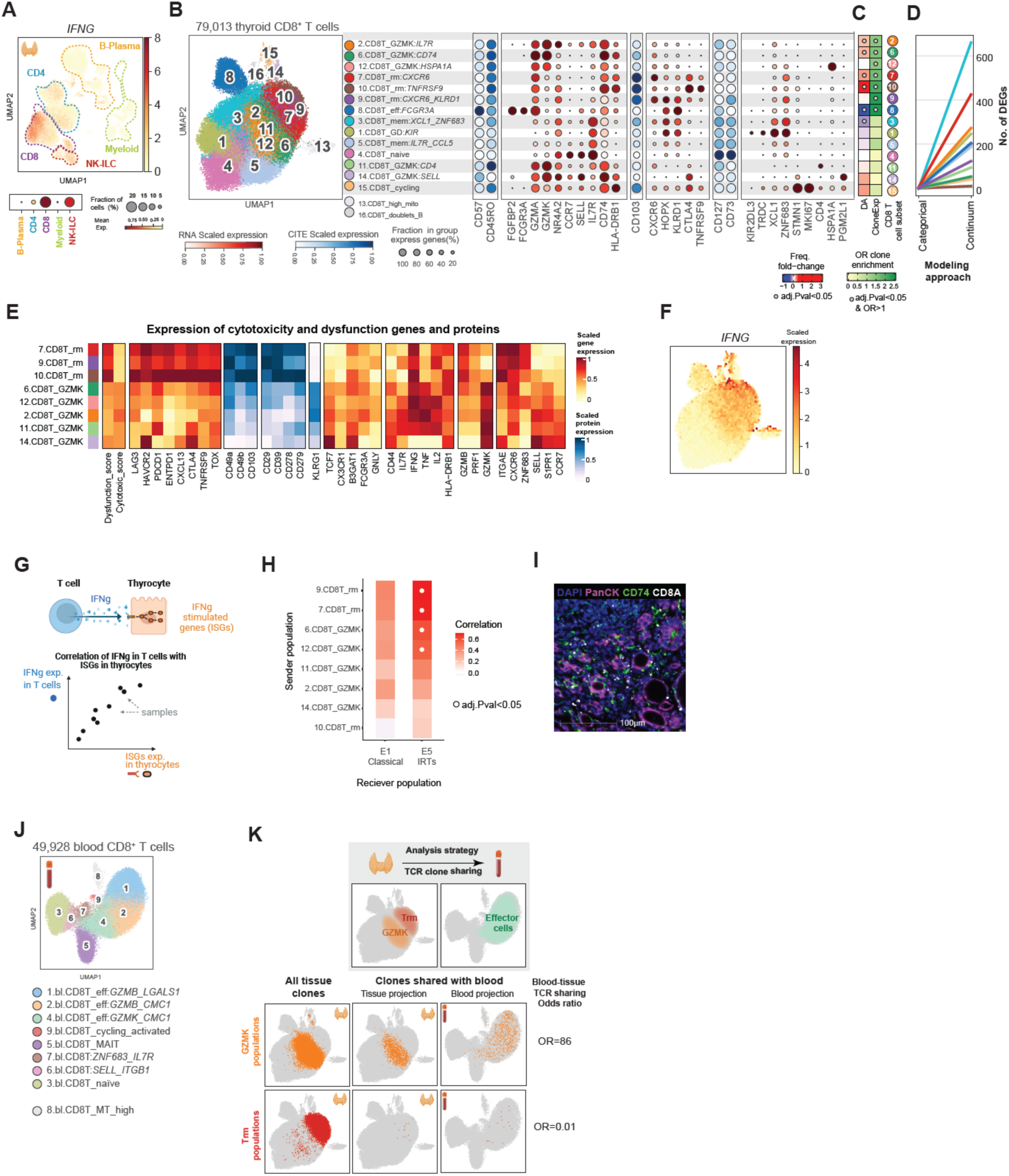
*GZMK*^+^ and T_rm_-like CD8^+^ T cells are key components of the tissue microenvironment during thyroid autoimmunity. **A.** UMAP embedding of thyroid-associated immune cells from scRNAseq of CD45^+^ cells from thyroid tissue, colored by *IFNG* expression (top). Dot plot of *IFNG* expression per lineage (bottom); dot size denotes percent of cells with non-zero expression and color indicates expression level. **B.** UMAP embedding of thyroid-associated CD8^+^ T cells (left) and dot plot illustrating main gene and protein markers (right). **C.** Differential abundance (DA, left) and clonal expansion enrichment (CloneExp, right) of CD8^+^ T cell populations. For DA, color denotes frequency log(fold-change) between samples with high and low immune infiltration. For CloneExp, color denotes odds ratio (OR) for enrichment of expanded clones. Gray dots denote significance. **D.** Number of differentially expressed genes (DEGs) in CD8^+^ T cell populations found by categorically comparing HT to GD patients (left) or by regression based on immune infiltration (right). FDR cut-od in both approaches is 0.05. Color denotes CD8^+^ T cell population. **E.** Heatmap of gene expression in the CD8^+^ GZMK^+^ cell populations and T_RM_-like cell populations. **F.** A hex-bin plot illustrating *IFNG* expression on CD8^+^ T cell UMAP embedding. **G.** Schematic illustrating the approach to identify CD8^+^ T populations correlated with IFN-**γ** response in 5.E_IRTs. **H.** Heatmap illustrating correlation between *IFNG* expression in CD8^+^ T cell populations (y-axis) and IFN-**γ** response expression in epithelial cell populations (x-axis). White dot denotes significance. **I.** Staining of thyroid tissue from a patient with HT for DAPI (blue), panCK (magenta), *CD74* (green) and *CD8A* (white). **J.** UMAP embedding of blood CD8^+^ T cells. **K.** Analysis strategy for TCR sharing (top): for a given tissue cell population, we quantify the clones shared between tissue and blood. Tissue-blood TCR clone sharing results for *GZMK*^+^ CD8^+^ T cells (middle row) and CD8^+^ T_rm_-like cells (bottom row). UMAP embedding of tissue-associated CD8^+^ T cells. Colored dots indicate cells from the population of interest with TCR clone data (left column). In the middle and right columns, colored dots indicate cells with TCR clones shared in both thyroid and blood, projected on the thyroid UMAP (middle) and blood UMAP (right).

We sub-clustered 79,013 CD8^+^ T cells based on RNA expression, and used surface protein data to annotate each cell population (Figure 3B, Supplementary figure 3F,G). We binned our samples into high or low immune infiltration, based on the frequency of immune cells determined by snRNAseq (Figure 1C, Figure 2D; high infiltration defined as >6%). Among the 14 cell subsets identified, 4 were more abundant in thyroid samples with high immune infiltration: 2.CD8T_GZMK^IL7R^, 6.CD8T_GZMK^CD74^, 7.CD8T_rm^CXCR^^6^, 10.CD8T_rm^TNFRSF^^9^ (Figure 3C, Supplementary figure 3H,I). Of note, we did not identify any populations that were specifically enriched in HT compared to GD (Supplementary Table 4). Thus, at the tissue level, GD and HT may represent a continuum of infiltration rather than distinct tissue inflammatory states. Similarly, when we examined differential gene expression with CD8^+^ T cell populations, a regression model that examined gene expression changes along the continuum of immune infiltration identified substantially more differentially expression genes compared to categorical analysis comparing gene expression in HT versus GD patient samples (Figure 3D, Supplementary Figure 12, Supplementary Table 19).

Phenotypically, 7.CD8T_rm*^CXCR^*^6^, 10.CD8T_rm*^TNFRSF^*^9^, and 9.CD8T_rm*^CXCR^*^6^*^,KLRD^*^1^ all expressed high levels of several genes and proteins associated with tissue residence (*ITGAE*, *CXCR6*, *ZNF683*, CD103, CD49a, CD49b) and we therefore collectively refer to them as T resident memory (T_rm_)-like cells. These cells had high expression of a gene module associated with T-cell dysfunction and low expression of a gene module associated with cytotoxicity^15^ (Figure 3E, Supplementary Table 6). 2.CD8T_GZMK*^IL7R^* was notable for strong expression of GZMK, along with 6.CD8T_GZMK*^CD74^*, 12.CD8T_GZMK*^HSPA1A^*, and 14.CD8T_GZMK*^SELL^* and 11.CD8T_GZMK*^CD^*^4^; we therefore collectively refer to these populations as *GZMK*^+^ *CD8*^+^ T cells. In contrast with T_rm_-like cells, GZMK^+^ CD8^+^ had intermediate expression scores for both gene modules associated with T-cell dysfunction and cytotoxicity, and showed higher expression of some markers of T-cell activation (*TNF*, *IL2*) (Figure 3E). Notably, although these several GZMK^+^ CD8^+^ T cell populations expressed high levels of *CXCR6*, suggestive of tissue-residence, most of these populations still expressed genes suggestive of circulating cells (*SELL*, *S1PR1*) (Figure 3E).

We examined *IFNG* expression within the CD8^+^ compartment and identified GZMK^+^ and T_rm_-like populations as the likely source of *IFNG* in tissue (Figure 3F). To examine probable signaling between GZMK^+^ and T_rm_-like CD8^+^ and 5.E_IRTs, we determined the correlation between *IFNG* expression in CD8^+^ T cells and our IFN-**γ** response signature in 5.E_IRTs as well as in 1.E_classical thyrocytes across all patients (Figure 3G). 9.CD8T_rm*^CXCR^*^6^*^,KLRD^*^1^, 7.CD8T_rm*^CXCR^*^6^, 6.CD8T_GZMK*^CD74^*, and 12.CD8T_GZMK*^HSPA1A^* showed significant correlation with IFN-**γ** response signature, specifically in 5.E_IRTs, suggesting that these populations may provide the IFN-**γ** that drives 5.E_IRT identity (Figure 3 G-H, Supplemental Figure 3J, Supplementary Table 7). Further supporting these results, RNAish imaging revealed that 5.E_IRTs are in close proximity to *CD8*^+^ T cells (Figure 3I). Collectively, these data suggest a critical role for *GZMK*^+^ and T_rm_-like *CD8*^+^ T cells in reshaping the thyroid cellular landscape during autoimmune infiltration.

### Single cell TCR-based lineage tracing of thyroid-associated *CD8*^+^ T cells suggests *in situ* expansion and phenotypic shaping of *GZMK*^+^ and T_rm_-like *CD8*^+^ T cells

We next examined how *CD8*^+^ T cells might populate the thyroid during autoimmune infiltration by leveraging the TCR-seq data as a lineage tracer. Somatic recombination of the genes encoding the *α* and *β* chains of the TCR locus in early T-cell development generates enormous diversity in the TCR repertoire, such that each individual T-cell has a unique TCR (“clone”) that functions as an exclusive identifier of that cell^16^. Therefore, in tissue, expanded CD8^+^ T cell clones – defined as TCR clones that are found in at least 1% of CD8^+^ T cells in each patient specimen (see Methods) – reflect activation and proliferation of a clonal population of CD8^+^ T cells, potentially targeting a single antigen of interest. CD8^+^ T cell populations with expanded clones are likely to be key players in tissue immune infiltration. Concordant with our differential abundance analysis, multiple *GZMK*^+^ and T_rm_-like CD8^+^ populations showed significant enrichment for expanded clones (2.CD8T_GZMK*^IL7R^*, 6.CD8T_GZMK*^CD74^*, 12.CD8T_GZMK*^HSP1A^*^1^, 7.CD8T_rm*^CXCR^*^6^, 10.CD8T_rm*^TNFRSF^*^9^, 9.CD8T_rm*^CXCR^*^6^*^,KLRD^*^1^); we also saw significant expansion in 8.CDT_ed*^FCGR3A^* (Figure 3C, Supplementary Figures 4A-C, Supplementary Tables 8, 9).

In addition, because scTCR-seq provides paired sequencing for TCR *α* and *β* chains in each single T cell, this allows lineage tracing of T cells by defining and tracking shared TCR clones^17^; if two T cells from the same sample share a TCR clone, we can infer that they must share a common precursor cell. Therefore, we leveraged TCR as a lineage tracer to precisely identify which expanded thyroid-associated *CD8*^+^ T populations shared a common cellular origin with 8 blood *CD8*^+^ T cell populations (Figure 3J, Supplementary Figure 4D-E). *GZMK*^+^ cells shared clones with effector T cells in blood (Figure 3K, top row; Supplementary Figure 4F, Supplementary Tables 10). Strikingly, T_rm_-like cells shared almost no TCR clones with blood *CD8^+^* T cells (Figure 3K, bottom row), suggesting that T_rm_-like cells proliferate and adopt their identity in the thyroid microenvironment.

### *CXCL9*-expressing dendritic cells are a hallmark of high immune infiltration and express genes associated with T cell activation

To determine how proliferation of CD8^+^ GZMK^+^ and T_rm_-like cells could be induced in tissue, we sub-clustered tissue-associated and blood mononuclear phagocyte (MNP) cells (Figure 4A, Supplementary Figure 5A, 18,155 cells and 49,997 cells respectively). As with other lineages examined, we did not find significant differences between blood MNP populations in patients with high and low infiltration in the thyroid (Supplementary Figure 5B). In the thyroid data, we found that 4 populations of dendritic cells (DC) – 4.MNP_DC2, 13.MNP_DC1, 11.MNP_DC5 and 9.MNP_DC3:CXCL9 – were all significantly more abundant in highly infiltrated samples (Figure 4B, Supplementary Figure 5C, Supplementary Table 4) and strongly correlated with immune infiltration (Supplementary Figure 5D, Supplementary Table 4). When we examined differential abundance based on disease status, none of these populations were significantly enriched in HT when compared to GD (Supplementary Table 4). Likewise, and similar to our findings with CD8^+^ T cells, a regression model that examined gene expression changes along the continuum of immune infiltration identified substantially more differentially expression genes compared to categorical comparison of HT versus GD (Figure 4C, Supplementary Figure 12, Supplementary Table 17). The presence of multiple DC populations, including those with phenotypes known to cross-present to CD8^+^ (13.MNP_DC1) and present to CD4^+^ T cells (4.MNP_DC2)^18^, suggests that the tissue milieu in high infiltration supports direct antigen-presentation to T cells. The marker genes associated with the 9.MNP_DC3:CXCL9 are consistent with a DC3 phenotype^19^, including MHC class II, *CD1C*, *CLEC10A* as well as intermediate levels of *CD163* and *CD14* (Figure 4A). In the tumor microenvironment, CD8^+^ T cell expression of IFN-**γ** has been shown to induce CXCL9^+^ myeloid cells; these cells may in turn recruit and retain CXCR3^+^ CD8^+^ T-cells in the tumor microenvironment^20–22^. Our data show that 9.MNP_DC3:CXCL9 are the primary source of CXCR3 ligands (i.e., *CXCL9*, *CXCL10*, *CXCL11*) among all tissue infiltrating immune cells (Figure 4D, E, Supplementary Figure 5E, Supplementary Table 11). Indeed, when we performed cell-cell interaction analysis between CD8^+^ and CD4^+^ T cells and MNPs, we found that, in addition to broad signaling between myeloid cells and T cells in general, 9.MNP_DC3:CXCL9 cells were uniquely poised to recruit and retain tissue infiltrating T cells via direct interactions between CXCR3 and its ligands, in addition to other chemokine-receptor interactions (Figure 4F, Supplementary Figure 6, Supplementary Tables 12, 13). In short, *IFNG*-expressing CD8^+^ T cells may induce *CXCL9*, *CXCL10*, and *CXCL11* expression in 9.MNP_DC3:CXCL9 cells, which in turn may recruit and retain those same CD8^+^ T-cells via CXCR3.

**Figure 4:**
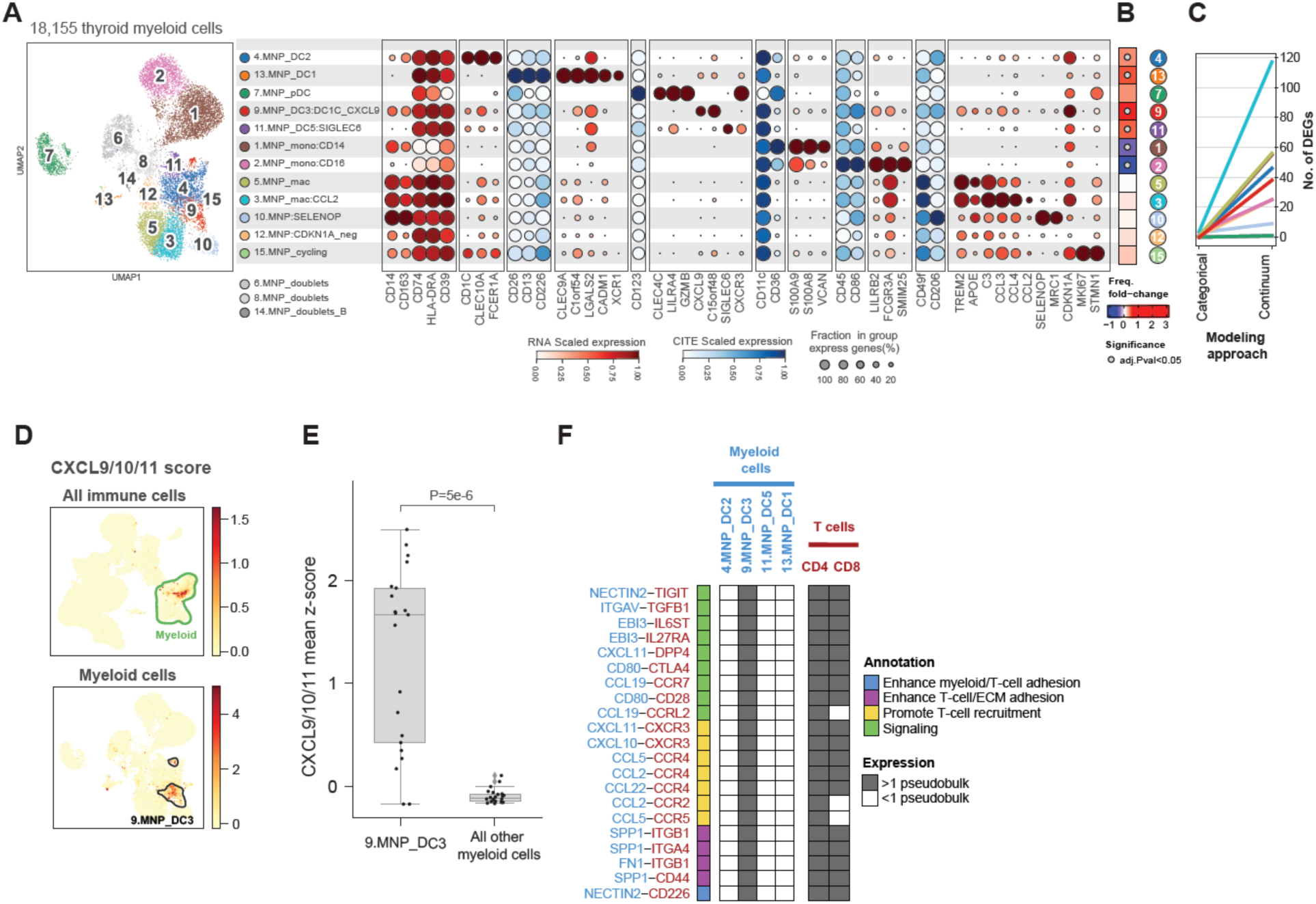
An immune infiltration-enriched *CXCL9*^+^ dendritic cell population expresses genes associated with T cell activation, recruitment and retention. **A.** UMAP embedding of thyroid-associated myeloid cells (left) and dot plot illustrating main gene and protein markers for each population (right). Dot size represents the percent of cells in the population with non-zero expression of a given gene or protein. Color indicates scaled expression. **B.** Differential abundance (DA) of myeloid populations.Color denotes frequency log(fold-change) between samples with high- and low-immune infiltration. Gray dots denote significant differential abundance. **C.** Number of differentially expressed genes (DEGs) in CD8^+^ T cell populations found by categorically comparing HT to GD patients (left) or by regression based on immune infiltration (right). FDR cut-od in both approaches is 0.05. Color denotes CD8^+^ T cell population. **D.** UMAP embedding of all thyroid-associated immune cells (top) and thyroid-associated myeloid cells (bottom) colored by mean Z-score of *CXCL9*, *CXCL10* and *CXCL11* expression. **E.** Boxplots quantify mean z-score of *CXCL9*, *CXCL10* and *CXCL11* expression in 9:MNP_DC3 versus other thyroid-associated myeloid cells. **F.** Putative 9.MNP:DC3-specific myeloid-T-cell paracrine interactions. Each row represents a potential protein-protein interaction between molecules expressed in myeloid cells (blue) and T cells (maroon). Myeloid and T-cell populations that express the interacting molecule at a pseudo-bulk abundance >1 are marked in gray.

Collectively, our data suggest that progressive immune infiltration is associated with IFN-**γ**-responsive signaling networks, in which IFN-**γ**-producing GZMK^+^ and T_rm_-like CD8^+^ T cells induce 5.E_IRTs and 9.MNP_DC3:CXCL9 cells, which in turn promote hub stability by locally recruiting and retaining these IFN-**γ**-producing CD8^+^ T cells.

### T_FH_ enrichment and clonal expansion of T_FH_ and T_reg_ are hallmarks of high immune infiltration

Since 5.E_IRTs specifically express components of MHCII and both 5.E_IRTs and 9.MNP*^CXCL^*^9^ cells are poised to interact with CD4^+^ T cells, we focused on defining the cellular landscape of tissue-associated CD4^+^ T cells. Sub-clustering 70,681 CD4^+^ T cells revealed 16 cell populations, including naïve, memory, and regulatory T cells (Figure 5A-B). As shown for CD8^+^ and myeloid cells, when we examined differential gene expression in CD4^+^ T cell populations, categorical analysis comparing HT and GD missed differentially expressed genes that were identified when the data were modeled along the immune infiltration continuum(Figure 5C, Supplementary Figure 12, Supplementary Table 17).

**Figure 5:**
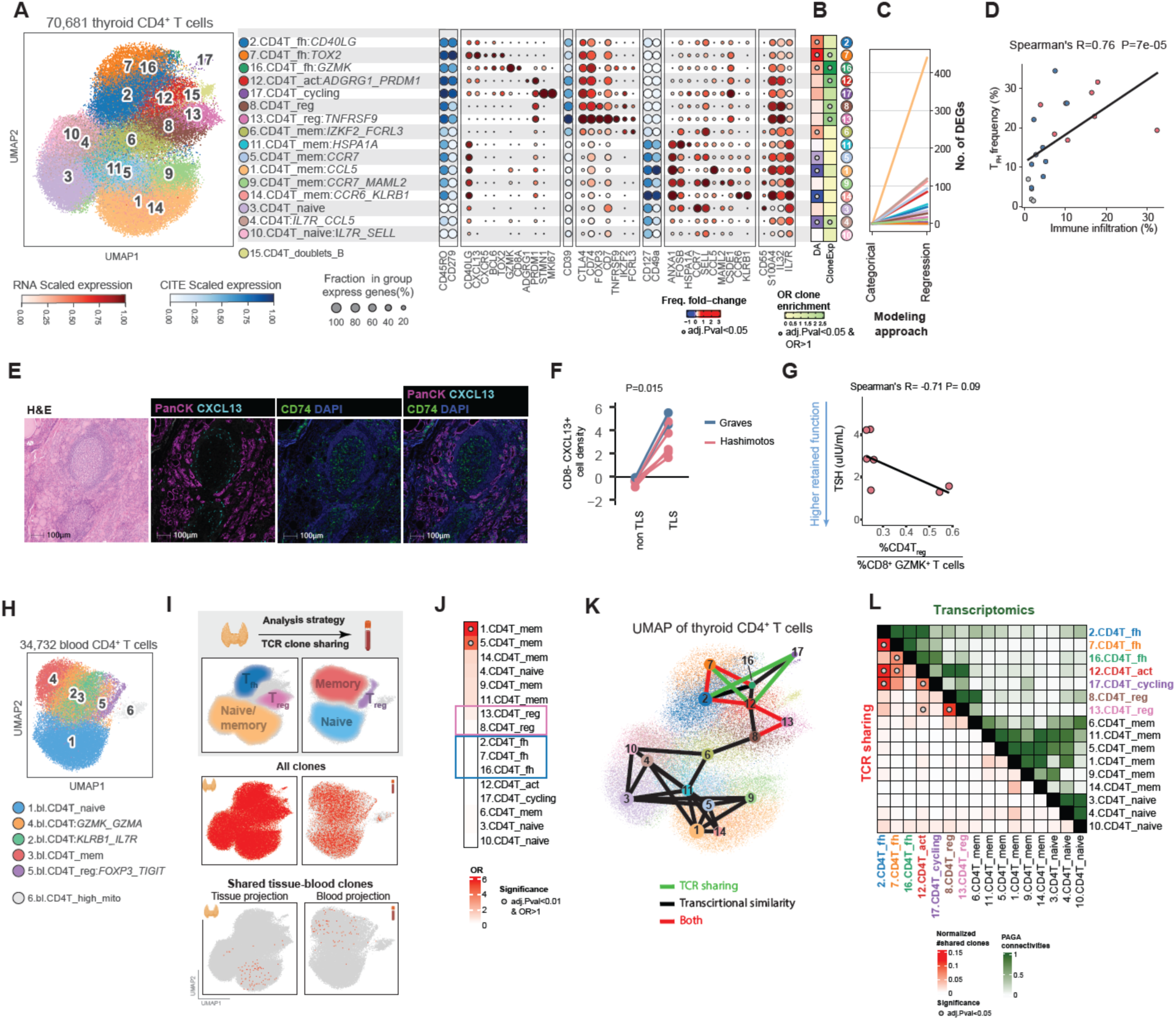
T_reg_ and T_fh_ cells emerge in tissue during immune infiltration. **A.** UMAP embedding of thyroid-associated CD4^+^ T cells (left) and dot plot illustrating main gene and protein marker (right). Dot size represents percent of cells with non-zero expression. Color indicates scaled expression. **B.** Differential abundance (DA, left) and clonal expansion enrichment (CloneExp, right) of CD4^+^ T cell populations. For DA, color denotes frequency log(fold-change) between samples with high- and low-immune infiltration. For CloneExp color denotes the odds ratio (OR) test for enrichment of expanded clones. Gray dots denote significance. **C.** Number of differentially expressed genes (DEGs) in CD4^+^ T cell populations found by categorically comparing HT to GD patients (left) or by regression based on immune infiltration (right). FDR cut-od in both approaches is 0.05. Color denotes CD4^+^ T cell population. **D.** Relationship between immune infiltration (x-axis) and frequency of T_fh_ among all CD4^+^ T cells (y-axis). **E.** Representative H&E image of a tertiary lymphoid structure (TLS) with staining for DAPI (blue), panCK (magenta), *CD74* (green) and CXCL13 (cyan). **F.** Quantification of *CD8*^-^*CXCL13*^+^ cells in TLS versus other regions in the thyroid based on imaging. Each line indicates an individual sample. **G.** Relationship between the ratio of T_reg_ to *GZMK*^+^ *CD8*^+^ T cells (x-axis) and thyroid function (serum TSH level, y-axis) in the 7 HT patients who were not on hormone replacement. The line represents linear regression of the data. **H.** UMAP embedding of blood CD4^+^ T cells. **I.** Top: analysis strategy for TCR sharing. Middle: UMAP embeddings of CD4^+^ T cells with TCR clone data for either tissue (left) or blood (right). Bottom: Cells with a TCR clone that is shared in both tissue and blood shown in the UMAP embeddings. **J.** Heatmap summarizing odds ratio (OR) for association between clone sharing of each thyroid-associated CD4^+^ T cell population with all blood CD4^+^ T cells. Boxes highlight no TCR sharing in T_reg_ and T_fh_. Color denotes OR values, dots denote significance. **K.** Relationship between tissue-associated CD4^+^ T cell populations based on TCR clone sharing (green), transcriptional profile (black) or both (red). Each pair of CD4^+^ T cell populations with a PAGA connectivity score >0.7 or a significant number of shared TCR clones are connected by an edge. The similarity network is overlaid on the thyroid-associated CD4^+^ T cell UMAP embedding. **L.** Heatmap quantifying pairwise association between thyroid-associated CD4^+^ T cell populations. Top(green) shows pairwise PAGA connectivity scores. Bottom (red) denotes clone sharing between each pair of thyroid-associated CD4^+^ T cell populations; color indicates normalized number of shared clones and dot denotes significance.

Three of the CD4^+^ populations (2.CD4T_fh*^CD40LG^*, 7.CD4T_fh*^TOX^*^2^, and 16.CD4T_fh*^GZMK^*) that were significantly more abundant in high infiltration (Figure 5B, Supplementary Table 4) and significantly correlated with immune cell infiltration to the tissue (Figure 5D, Supplementary Table 4) expressed marker genes characteristic of T follicular helper (T_FH_) cells (*CXCL13*, *CXCR5*, *BCL6*, *TOX2*, *CD40LG*) (Figure 5A, B, Supplementary figure 7A). To validate that these were indeed T_FH_ cells, we performed RNAish and found that *CXCL13*^+^*CD8*^-^ T cells enriched in close apposition to tertiary lymphoid structure (TLS) (Figure 5E, F, Supplementary Table 14), which are ectopic lymphoid aggregates with visible germinal centers that form in chronic inflammation^23^. Among all thyroid-associated immune cells, *CXCL13* was primarily expressed in *CD4*^+^ and *CD8*^+^ T cells (Supplementary figure 7B). Therefore, the *CXCL13*^+^*CD8*^-^ T cells seen on imaging likely correspond to the scRNAseq clusters 2.CD4T_fh^CD40LG^, 7.CD4T_fh*^TOX^*^2^, and 16.CD4T_fh*^GZMK^*, which in turn likely represent true T_FH_ cells that may promote the formation and maintenance of TLS germinal centers.

We found significant enrichment of expanded TCR clones in 7.CD4T_fh*^TOX^*^2^, 16.CD4T_fh*^GZMK^* as well as a likely activated *CD4*^+^ population (12.CD4T_act*^ADGRG^*^1^*^,PRDM^*^1^), and two populations of regulatory T cells (8.CD4T_reg and 13.CD4T_reg*^TNFRSF^*^9^) (Figure 5B, Supplementary figure 7C-E, Supplementary Tables 8, 9). Given the clonal expansion observed in T_reg_ populations, we examined whether T_reg_ might contribute toward restricting the activity of effector-like *GZMK*^+^ *CD8*^+^ T cells. Indeed, among patients with HT who were not on hormone replacement, thyroid function (TSH) was anti-correlated with the frequency ratio of *CD4*^+^ T_reg_ to *GZMK*^+^ *CD8*^+^ T cells (Figure 5G, Supplementary Table 15); thus an increased *CD4*^+^ T_reg_ to *GZMK*^+^ *CD8*^+^ ratio is associated with improved thyroid function.

### T_FH_, T_reg_, and activated CD4^+^ T cells may expand and adopt their phenotypic state *in situ*

Strikingly, we found almost no shared TCR clones between any of these 5 expanded tissue populations and *CD4*^+^ T cells in blood (Figure 5 H-J, Supplementary figure 7F-I, Supplementary Table 10). Only 1.CD4_mem*^CCL^*^5^ and 5.CD4T_mem*^CCR^*^7^ had a significant enrichment of shared clones with blood *CD4^+^* T cells. Both these populations are memory T cells that were enriched in samples with low immune infiltration (Figure 5B, J), suggesting that these were in fact circulating populations. Notably, tissue T_reg_ populations (8.CD4T_reg and 13.CD4T_reg*^TNFRSF^*^9^) together shared only one clone from one patient with blood, suggesting that tissue-infiltrating T_reg_ cells in thyroid autoimmunity are induced in the thyroid rather than recruited from the blood.

To understand the origin of thyroid-infiltrating T_FH_ and T_reg_, we examined both TCR clone sharing and transcriptional similarity among thyroid-associated *CD4*^+^ T cell populations. As expected, we noted significant TCR sharing and high transcriptional similarity between T_reg_ populations and transcriptional overlap and shared TCR clones among T_FH_ populations (Figure 5K, L, Supplementary Table 16). Furthermore, we noted significant TCR clone sharing between cycling cells (17.CD4T_cycling) and both 2.CD4T_fh^CD40LG^, suggesting that T_FH_ cells are actively replicating in tissue and consistent with clonal expansion in these populations. Surprisingly, we noted that 12.CD4T_act^ADGRG^^1_PRDM1^ shared both TCR clones and transcriptional overlap with both T_reg_ and T_FH_ populations, suggesting that these populations share a common precursor.

Collectively, these data suggest that both T_FH_ and T_reg_ cells may be induced and expand in the thyroid in HT and GD.

### TLS are a hallmark of immune infiltration in thyroid autoimmunity

To define how thyroid-infiltrating T_FH_ cells, which provide B cell help and support maturation, contribute to progression of thyroid autoimmunity, we sub-clustered thyroid-associated B-cell populations. Sub-clustering of 51,047 single B and plasma cells identified 9 populations, including 7.B_GC and 8.B_GC_cycling. These two populations were higher in abundance in samples with high immune infiltration and expressed markers of germinal center B cells *(TCL1A*, *BCL7*, *KLHL6*); 8.B_GC_cycling additionally expressed *MKI67* and other markers of cycling cells (Figure 6A, B, Supplementary Figure 8A, Supplementary Table 4). Additionally, consistent with all other lineages, differential gene expression identified more differentially expression genes when we examined gene expression changes along the continuum of immune infiltration as opposed to categorical analysis by disease (Figure 6C, Supplementary Figure 7F, Supplementary Table 17). Similarly, germinal center B cells were strongly correlated with immune infiltration (Figure 6D, Supplementary Table 4), consistent with the presence of TLS in thyroid autoimmunity. Paired BCR data analysis did not reveal enrichment of expanded clones in any B cell populations (Supplementary Figure 8B-D). Furthermore, we did not find significant differences in B cells from paired blood between patients with high and low infiltration (Supplementary Figure 8E,F).

**Figure 6:**
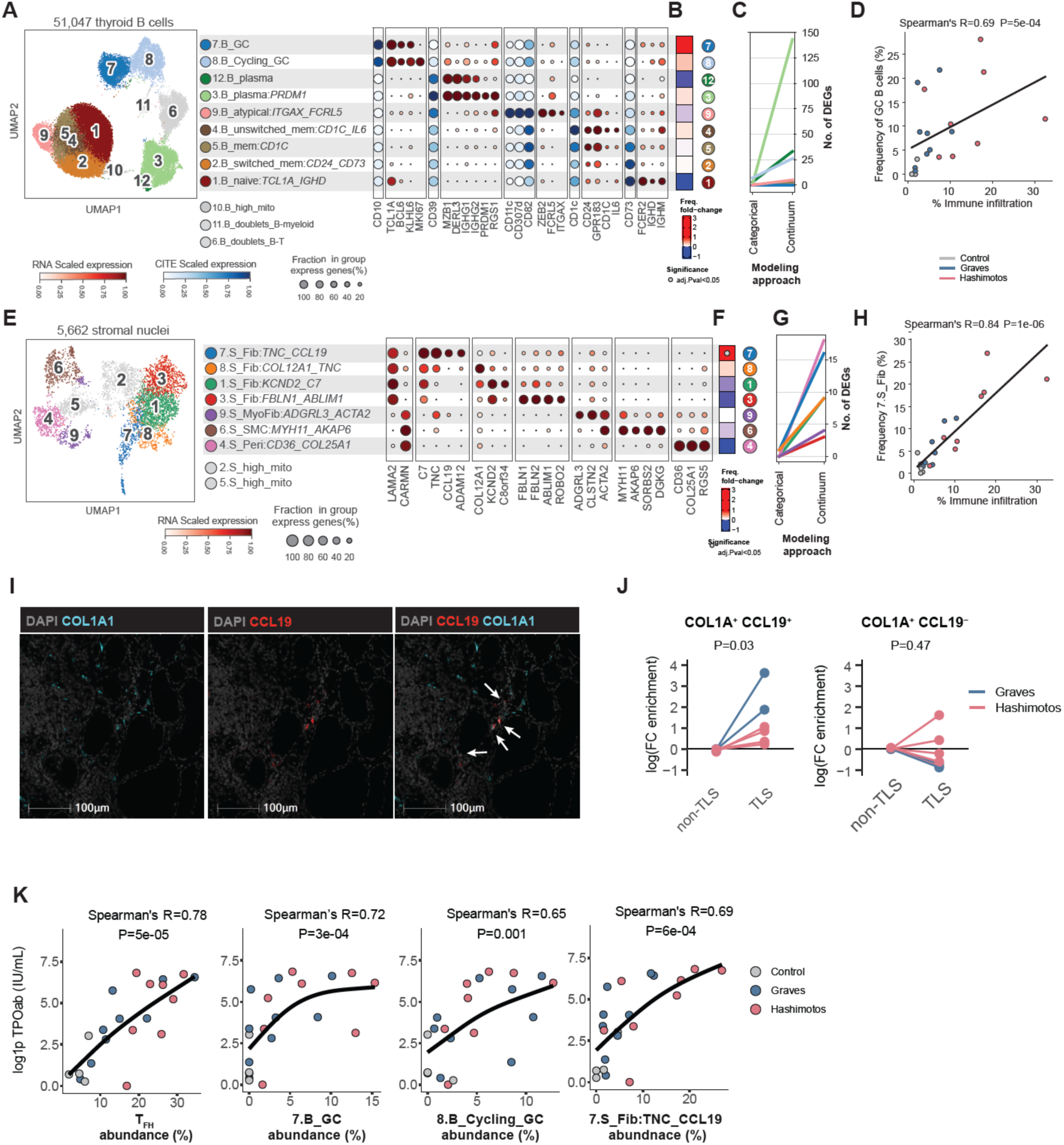
Components of tertiary lymphoid structures, *CCL19*^+^ fibroblasts and germinal center B cells, are a hallmark of thyroid autoimmunity and strongly correlated with anti-TPO antibodies. **A.** UMAP embedding of thyroid-associated B lineage cells (left) and dot plot illustrating main gene and protein markers (right). Dot size represents percent of cells with non-zero expression. Color indicates scaled expression. **B.** Differential abundance (DA) of B lineage populations. Color denotes log(fold-change) of frequency between high- and low-infiltration samples. **C.** Number of differentially expressed genes (DEGs) in B lineage cell populations found by categorically comparing HT patients to GD patients (left) or by regression based on immune infiltration (right). FDR cut-od in both approaches is 0.05. Color denotes B cell population. **D.** Relationship between immune infiltration (x-axis) and frequency of germinal center B cells (7.B_GC and 8.B_cycling_GC, among all B lineage cells, determined by scRNAseq, y-axis). The line represents linear regression of the data. **E.** UMAP embedding of thyroid-associated stromal cells from snRNAseq (left) and dot plot displays marker genes for each population (right). **F.** Differential abundance (DA) of stromal cell populations. Color denotes log(fold-change) of frequency between high- and low-infiltration samples. Gray dots denote significance. **G.** Number of differentially expressed genes (DEGs) in stromal cell subsets found by comparing HT patients to GD patients (categorial) or by regression based on immune infiltration. FDR cut-od in both approaches is 0.05. **H.** Relationship between immune infiltration (x-axis) and frequency of 7.S_Fib^TNC,CCL^^19^ among all stromal cells, determined by snRNAseq (y-axis). The line represents linear regression of the data. **I.** RNAish staining of thyroid tissue from a patient with HT for DAPI (gray), *COL1A1-2* (teal), and *CCL19* (red). White arrows indicate *CCL19^+^* fibroblasts, which stain for both *COL1A1* and *CCL19*. **J.** Quantification of *CCL19^+^* fibroblasts in TLS versus other regions of thyroid tissue. Each line indicates an individual sample. **K.** Relationship between frequency of putative TLS-associated cell types (x-axes) and serum anti-TPO autoantibody level (y-axis). Lines represent general additive models.

Stromal and endothelial cells are associated with TLS^23^; we therefore sub-clustered these lineages to determine how they might contribute to TLS formation (Figure 6E, Supplementary Figure 9A-D). Among 5,662 stromal cells profiled, we identified a population of *CCL19*^+^ fibroblasts, 7.S_Fib:TNC_CCL19, that were significantly more abundant in tissue samples with high infiltration (Figure 6F, Supplementary Figure 9E) and strongly correlated with immune infiltration and, like other stromal cells showed more differentially expressed genes when modeled along the continuum of immune infiltration (Figure 6G-H, Supplementary Table 4). We validated the presence of *CCL19*^+^ fibroblasts in our thyroid samples using RNAish (*CCL19*^+^*COL1A1-2*^+^ cells, Figure 6I) and determined that these fibroblasts, the imaging correlate of 7.S_Fib*^TNC,CCL^*^19^, were enriched at sites of TLS which was not the case for *CCL19*^-^ fibroblasts (Figure 6J, Supplementary Table 14). Consistent with the suspected role of TLS in autoantibody production, the frequencies of 7.S_Fib:*^TNC,CCL^*^19^, germinal center B cells, and T_FH_ cells were all strongly correlated with anti-thyroid peroxidase antibodies, autoantibodies that are diagnostic of Hashimoto’s thyroiditis^6^ (Figure 6K, Supplementary Table 4).

### IRTs express T cell checkpoint ligands and colocalize with *CD8*^+^ T cells in IFN-γ-responsive immunity hubs

To understand how TLS corresponded to clinical phenotype, we examined the fraction of thyroid tissue covered by TLS as a function of immune infiltration for 11 patients – including control, GD, and HT – spanning a range of immune infiltration. Surprisingly, the two samples from patients with HT with impaired thyroid function (AT47, AT42) were clear outliers (Figure 7A, Supplementary Table 18) in the otherwise tight correlation between immune infiltration and TLS in tissue. This suggests that while the frequency of immune cells does not predict thyroid function (Figure 1E), the tissue distribution of specific cell populations may influence thyroid function. We trained and implemented an AI-based tissue classifier (HALO AI) to identify three different tissue regions: (i) dense lymphocytic infiltrates (including TLS),(ii) sparse immune infiltration, and (iii) normal thyroid tissue (Figure 7B). Strikingly, even patient samples with substantial immune infiltration consisted primarily of regions of normal tissue architecture (Figure 7C). We confirmed that thyroid from patients with impaired thyroid function (AT47, AT42) exhibited more densely infiltrated regions (including TLS) with relatively fewer sparse or normal regions (Figure 7C). To understand how thyroid function is retained in some patients with HT, we examined whether some infiltration-enriched cell populations might be enriched in sparsely infiltrated regions of tissue. Indeed, *CD8*^+^ T cells and panCK^+^*CD74*^+^ cells, which are the imaging correlate of IRTs, were enriched in sparsely infiltrated regions, whereas classical thyrocytes (i.e., panCK^+^*CD74*^-^) were not (Figure 7D, for *CD8*^+^ T cells and panCK^+^*CD74*^+^ cells paired Mann-Whitney test P=0.004, Supplementary Figure 10A, Supplementary Table 19). Enrichment of *CD8^+^* T cells and IRTs appeared less pronounced for patients with loss of thyroid function, suggesting that the distribution of these cell populations could influence thyroid function.

**Figure 7:**
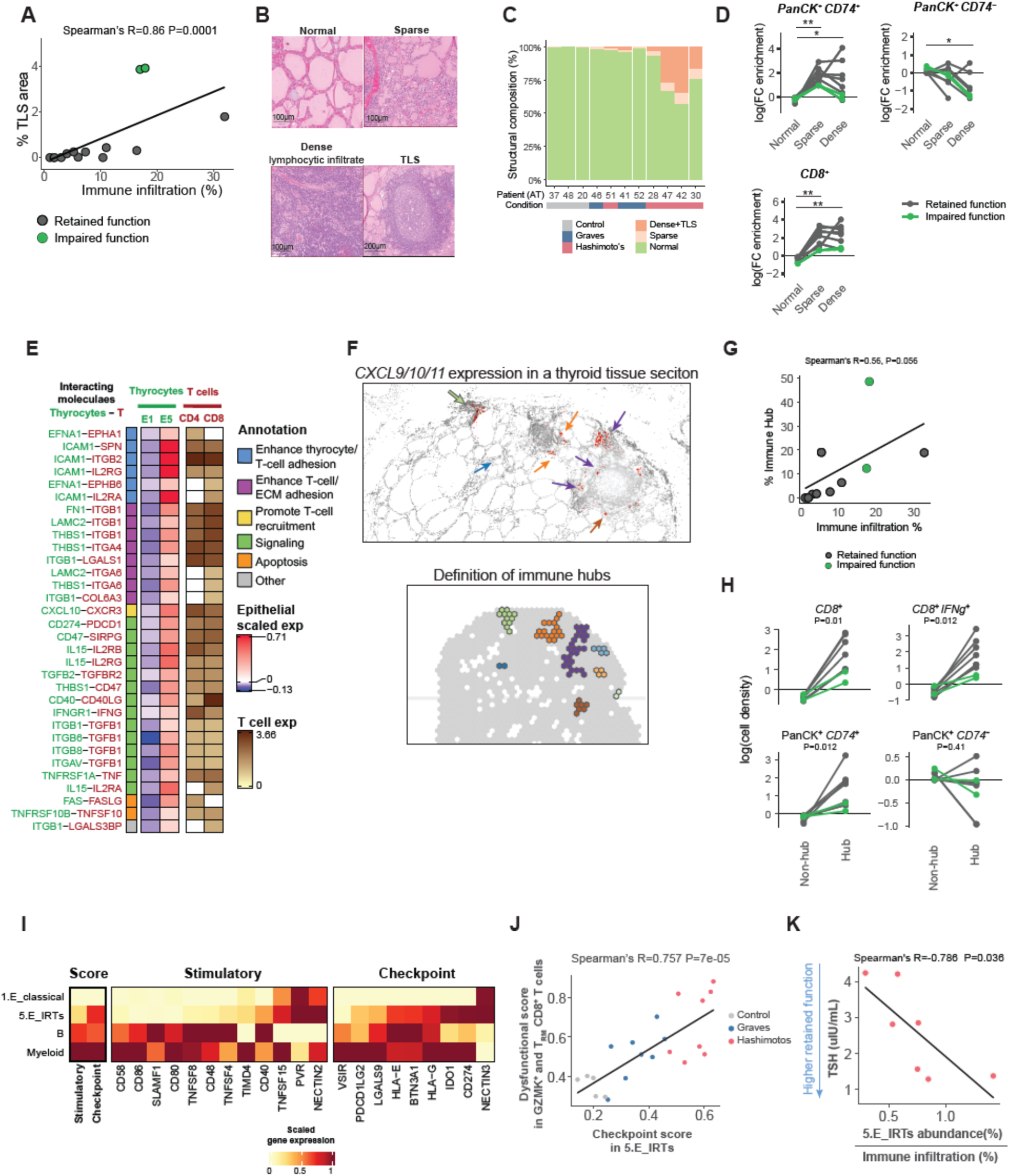
IFN-γ response establishes an infiltration-specific microenvironment. **A.** Relationship between immune infiltration (x-axis) and tertiary lymphoid structure (TLS) area (out of total area of tissue section (y-axis). Color denotes samples from patients with retained or impaired thyroid function. The line represents linear regression of the data. **B.** Representative H&E staining images of four macroscopic tissue structures in a thyroid sample: normal thyroid tissue, sparsely infiltrated tissue, dense lymphocytic infiltrate, and TLS. **C.** Distribution of macroscopic tissue structures within each thyroid section. Each bar represents an individual specimen; color indicates tissue structure. **D.** Density of cellular phenotypes in three major macroscopic regions: Normal, sparsely infiltrated (Sparse), and combined dense lymphocytic infiltrate and TLS (Dense). Cellular phenotypes shown are: panCK^+^*CD74*^+^ (corresponding to IRTs), panCK+*CD74*^-^ (corresponding to classical thyrocytes) and *CD8A*^+^ T cells. Dots denote average frequency of each cell type per region type. Each line represents an individual specimen. *CD8A*^+^: dense vs normal P=0.004 ;*CD8A*^+^ sparse versus normal P=0.004; panCK^+^*CD74*^+^ dense versus normal P=0.02; panCK^+^*CD74*^+^ sparse versus normal P=0.004; panCK^+^*CD74*^-^ dense versus normal P=0.019. **E.** Putative thyrocyte-T cell paracrine interactions. Each row represents a potential protein-protein interaction between molecules expressed in thyrocytes (1.E_classical and 5.E_IRTs) and T cells (*CD4*^+^ and *CD8*^+^). Color for thyrocyte columns represents scaled expression across 1.E_classical and 5.E_IRTs. Color in T cell columns represents expression in log(CPM). White boxes indicate genes expressed < 1 in pseudo-bulk. **F.** Representative image of thyroid tissue section stained DAPI (gray), *CXCL9/CXCL10/CXCL11* (red, top). Colored arrows indicate immunity hubs corresponding to the bottom image. Bottom: Approach to define IFN-**γ** response hubs. A hexagonal tessalon (35um side length) was overlaid on images and *CXCL9/CXCL10/CXCL11^+^*windows were identified based on the presence of a *CXCL9/CXCL10/CXCL11^+^*and adjacent windows were aggregated into immunity hubs (bottom). **G.** Relationship between immune infiltration (x-axis) and area of tissue covered by immunity hubs (y-axis). The line represents linear regression of the data. **H.** Density of cellular phenotypes in immunity hubs versus the rest of the thyroid tissue section. Each line represents an individual specimen **I.** Scaled pseudo-bulk gene expression of stimulatory and checkpoint genes in 1.E_classical thyrocytes, 5.E_IRTs thyrocytes, B lineage cells, and myeloid cells, determined by snRNAseq. Stimulatory and checkpoint scores (left) indicate mean value of stimulatory or checkpoint gene expression, respectively. **J.** Relationship between checkpoint score in 5.E_IRTs (x-axis) and dysfunctional score (defined in Figure S3I) in *GZMK*^+^ and T_RM_ CD8^+^ T cells. Each dot represents an individual patient. The line represents linear regression of the data. **K.** Relationship between the ratio of 5.E_IRT frequency out of all thyrocyte populations to immune infiltration (x-axis) and thyroid function (serum TSH levels, y-axis) in the 7 HT patients who were not on hormone replacement. The line represents linear regression of the data.

Given co-enrichment of both panCK^+^*CD74*^+^ cells (IRTs) and *CD8^+^* T cells in sparsely infiltrated regions, it is possible that signaling interactions between these cell types might facilitate co-localization. To determine whether 5.E_IRTs might also provide feedback to retain T cells, we examined putative interactions between 5.E_IRTs and CD8^+^ or CD4^+^ T cells. 5.E_IRTs, but not 1.E_classical thyrocytes, were poised to recruit CXCR3^+^ T cells, promote direct IRT-T cell interactions, and promote T-cell adhesion to thyroid extracellular matrix (Figure 7E, Supplementary Figure 10B-D, Supplementary Table 20). These putative interactions suggest the IRTs can locally recruit and retain T cells in the thyroid parenchyma.

To understand how IFN-**γ**-dependent signaling could alter local tissue organization, we examined immunity hubs, which have been previously defined as microscopic foci of disease-specific T-cells that are poised to interact directly with parenchymal or other cell populations. These immunity hubs have been found to regulate disease progression in both colorectal^24^ and lung^25^ cancer. As previously described, we defined an IFN-**γ**-responsive immunity hub as the contiguous neighborhood around *CXCL9*/*CXCL10*/*CXCL11*^+^ cells (Figure 7F, Supplementary Figure 11). Our analysis identified 3,337 IFN-**γ**-responsive immunity hubs across 11 samples. The fraction of tissue covered by these immunity hubs was strongly correlated with immune infiltration (Figure 7G, Supplementary Table 21). Additionally, *CD8*^+^ T cells, *CD8*^+^*IFNG*^+^ T cells and panCK^+^*CD74*^+^ cells were enriched in immunity hubs relative to other parts of tissue, whereas classical thyrocytes (i.e.,panCK^+^*CD74*^-^) were not (Figure 7H, Supplementary Table 22). Strikingly, enrichment was less pronounced for two patients with impaired thyroid function.

Given this apparent hub enrichment of panCK^+^*CD74*^+^ cells (i.e., IRTs) in patients with retained thyroid function as well as our finding that the 5.E_IRTs were poised to recruit and interact with T cells (Figure 4F), we assessed how 5.E_IRTs might signal to T cells. We constructed a gene expression score for co-stimulatory versus T cell checkpoint molecules and found that 5.E_IRTs, but not classical thyrocytes, were strongly enriched for expression of checkpoint molecules rather than co-stimulatory molecules (Figure 7I, Supplementary Table 23). Strong expression of checkpoint molecules in 5.E_IRTs and expression of a gene module associated with T cell dysfunction in T_rm_-like and GZMK^+^ CD8+ T cells (Supplementary Figure 3H) raises the possibility that IRTs might be poised to promote CD8+ T cell dysfunction. Indeed, on a per patient basis, we found a highly significant correlation between checkpoint score in 5.E_IRTs and dysfunction score in GZMK^+^ and T_rm_-like CD8^+^ T cells, suggesting that IRTs may be capable of directly promoting T cell checkpoints and consequent dysfunction (Figure 7J, Supplementary Table 24). Strikingly, we found that the ratio of 5.E_IRTs to immune infiltration was strongly anti-correlated with TSH (i.e., that a higher ratio of 5.E_IRTs to immune cells implied greater thyroid function) in patients with HT who were not on hormone replacement (Figure 7K, Supplementary Table 25). Collectively, these data suggest that IFN-**γ**-responsive hubs may localize immune-modulatory 5.E_IRTs with *CD8*^+^*IFNG*^+^ T cells, thus decreasing T-cell activity and preserving tissue function.

## DISCUSSION

Our comprehensive multimodal atlas of the human thyroid in health and disease leverages this singular model to dissect autoimmunity directly in human tissue. Our data suggest that distinct multicellular signaling networks are hallmarks of progressive immune infiltration and that tissue organization, rather than immune infiltration *per se*, may be a key regulator of progressive infiltration and loss of function.

We identify an infiltration-correlated thyrocyte population that upregulates IFN-**γ**-stimulated genes, MHCII machinery, and checkpoint-promoting genes, which we term IRTs (IFN-**γ**-responsive thyrocytes). It has long been established that MHCII is expressed in thyrocytes in patients with HT and GD^26,27^, however our data demonstrate that this expression is confined to a small, specifically localized population of thyrocytes that appear poised for immune-modulatory capability. We show that IRTs are enriched in IFN-γ responsive immunity hubs, distinct microenvironments where *CD8*^+^*IFNG*^+^ T cells, IFN-γ responsive *CXCL9/CXCL10/CXCL11*-expressing cells, and IRTs co-localized, suggesting that the these hubs may facilitate interactions between immune-modulatory IRTs and *CD8*^+^ T cells, potentially influencing local T-cell activity and preserving tissue function in HT. Strikingly, we show that IRTs express T cell checkpoint-promoting molecules in proportion to T cell expression of a dysfunction signature and that the relative abundance of IRTs to immune cells correlates with thyroid function. One limitation of our approach was the relatively small number of HT patients with loss of thyroid function. Further exploration of IFN-γ-responsive immunity hubs in larger datasets could address this and elucidate the processes underlying loss of tissue function in thyroid autoimmunity and in tissue-specific autoimmunity more broadly.

Moreover, we show that even high-infiltration samples have a majority of preserved normal tissue architecture, which suggests that the immune hubs have a key role in regulating local T-cell activity and preserving tissue function in HT. This may help explain why most individuals with inappropriate thyroid immune infiltration have preserved thyroid function. Indeed, such discontinuous pathological immune infiltration is a characteristic feature across autoimmune disease, evidenced by skip lesions in Crohn’s disease^28^, heterogeneous islet involvement in type 1 diabetes^29^ or discontinuous skin involvement in plaque psoriasis^30^. It is intriguing to speculate that parenchymal cells, such as thyrocytes, may facilitate this discontinuity.

A distinct feature of the tissue microenvironment in both HT and GD was the presence of TLS. Across all patient samples, regardless of underlying disease, key TLS cell populations in the thyroid -- T_FH_, germinal center B cells, and *CCL19*^+^ fibroblasts -- were strongly correlated with systemic levels of thyroid-specific autoantibodies (anti-TPO antibodies) that are diagnostic of HT. This correlation provides compelling human evidence to the prevailing concept that tissue-specific autoantibodies form in TLS in tissue autoimmunity. We identified an infiltration-dependent stromal population of CCL19^+^ fibroblasts (7.S_Fib*^TNC^*^,*CCL*^^19^) that are enriched at TLS, consistent with data from a mouse model of Sjogren’s disease in which *CCL19*^+^ fibroblasts were critical drivers of early TLS formation^31^. Furthermore, 7.S_Fib*^TNC^*^,*CCL*^^19^ shared similar features with a previously published *CXCL10*^+^ *CCL19*^+^ fibroblast population that was found to be broadly prevalent in human autoimmunity across tissue samples, including salivary gland, lung, synovium and gut^32^. Integrating these findings with our data suggest that *CCL19*^+^ fibroblasts may also be drivers of TLS formation in humans across disease.

The diverse local structures within the tissue microenvironment, as described above, may additionally support continued *in situ* evolution of cellular phenotype, which in turn contributes to disease progression. For example, by using scTCR-seq as a lineage tracer, our data argue that several T-cell populations expand and adopt their phenotypic state *in situ*. These include CD4^+^ T_FH_ and T_reg_ cells, as well as CD8^+^ Trm-like cells and *GZMK*^+^ cells. The extraordinary diversity and uneven distribution of the TCR repertoire constrains comprehensive sampling^33,34^, thereby limiting our capacity to fully track the developmental trajectories of clones, despite analyzing TCR data from 84,573 cells. Furthermore, exceptionally rare, transient cell states may be key determinants of trajectory that are difficult to capture in human tissue samples, where it is not possible to monitor the same tissue over time.

Strikingly, we determine that although GD and HT are different clinical entities driven by distinct underlying mechanisms in response to different autoantigens^10,35–37^, they converge onto a single continuum of progressive thyroid immune infiltration. Indeed, we consistently observed this continuum through analyses of epithelial, stromal, and immune cell population abundance, infiltration-associated gene expression changes across all cell populations, autoantibody production, and in the frequency of specific tissue sub-structures including TLS and IFN-γ responsive immunity hubs. These data harmonize with existing epidemiologic and genetic evidence that GD and HT share underlying genetic architecture^38,39^, coincide in the same families and even in the same patients^40–42^ and share common risk factors^43^. In short, by modeling thyroid immune infiltration as a continuous spectrum, rather than as discrete categories, we uncover a shared tissue endotype that underlies both conditions. This suggests that – despite distinct antigenic triggers and clinical manifestations – GD and HT might reflect distinct stages or degrees of a common autoimmune ecosystem. Specific tissue tolerance mechanisms may act as critical regulators that maintain homeostasis or slow progression along this shared autoimmune continuum. Indeed, induced autoimmune thyroid disease, such as checkpoint inhibitor-induced thyroiditis^44^ or postpartum thyroiditis, may reflect breaching of these specific tolerance mechanisms and accelerating progression along this continuum. This paradigm of shared autoimmune continuum, modulated by tissue-specific checkpoints, may extend beyond thyroid autoimmunity and offer a new framework for understanding, and potentially intervening in, the progression of other organ-specific autoimmune disease.

## ACKNOWLEDGEMENT

We are deeply grateful to all study participants. M.R. is supported by the National Institutes of Health K08 Award (5K08DK133497-01) and was supported by a National Institutes of Health T32 Award (T32DK007191). R.N. was supported by a Massachusetts General Hospital ECOR Medical Discovery Research Fellowship. This work was made possible by the generous support from the Doris Duke Physician-Scientist Fellowship (to M.R.), an American Thyroid Association Research Grant (to M.R.), the Massachusetts General Hospital Gilbert H. Daniels, MD Award for Innovative Research in Thyroid Cancer (to M.R.), the Massachusetts General Hospital John T. Potts, Jr. Pilot Award Program (to M.R.), the National Institute of Health Director’s New Innovator Award (DP2CA247831; to A.C.V.), the Chan Zuckerberg Initiative (2023-323359 (5022) GB-1587789; to A.C.V.), the Lupus Research Alliance (835103; to A.C.V.), Juvenile Diabetes Foundation (to A.C.V.), the Broad Institute Next Generation Fund(to A.C.V.), the Massachusetts General Hospital Transformative Scholar in Medicine Award (to A.C.V.), and the Massachusetts General Hospital Howard M. Goodman Fellowship (to A.C.V.).

## AUTHOR CONTRIBUTIONS

M.R., R.N., and A.C.V. conceived of and led the study; M.R. and A.C.V. led the experimental design; M.R. generated the single-cell sequencing data with contribution from B.A. and A.T. R.N. led the design of and performed single-cell sequencing computational analysis, with contribution from H.T.; K.S. developed web browser to visualize single-cell data; M.R., L.T.N, K.X., P.S., E.R., and Y.S. designed and performed microscopy experiments and analysis; M.R., M.C., A.E.S, P.S., S.P., G.H.D., A.D.L provided clinical expertise, coordinated and performed sample acquisition, and/or established research protocols; A.D.L and A.C.V. supervised the study; M.R. and A.C.V. provided funding for this work; M.R., R.N., and A.C.V. wrote the manuscript, with input from all authors.

## METHODS

### Patient selection, surgical tissue and blood acquisition and processing

Thyroid surgical samples and paired blood was obtained from patients undergoing clinically indicated thyroid surgery at Massachusetts General Hospital or Newton-Wellesley Hospital. Informed consent was obtained from all patients in accordance with a protocol approved by the Mass General Brigham institutional review board (2008P001466). We excluded patients who had received current or prior immune-modulatory therapies and excluded patients on drugs known to affect thyroid function except levothyroxine, methimazole, and potassium iodide. Control and HT samples were obtained from patients undergoing total thyroidectomy or completion thyroidectomy for a unilateral thyroid nodule and the indication was removal of the contralateral lobe was required to be either planned radioactive iodine treatment, monitoring, or patient preference. We excluded patients for whom there were nodules or other concerns in the contralateral lobe, and samples were obtained from the contralateral lobe. For control samples, we excluded data from samples for which clinical pathology revealed chronic lymphocytic thyroiditis or other diffuse abnormalities. For HT samples, we excluded data from samples for which clinical pathology did not confirm chronic lymphocytic thyroiditis. HT patients included both euthyroid patients (not on levothyroxine) and hypothyroid patients (on levothyroxine). GD samples were obtained from patients undergoing total thyroidectomy as treatment for GD. Clinical data, including demographics, prior drug exposure, and past medical history including prior thyroid function, were obtained retrospectively from electronic medical records.

Following resection, thyroid surgical tissue specimens were immediately placed in ice-cold HypoThermosol solution (BioLife Solutions, Bothell, WA) and kept on ice during transfer to the research facility. Tissue was then washed twice with cold phosphate buffered saline and dissected into separate sections for dissociation for scRNA-seq and flash freezing for single nuclei isolation. Individual sections for downstream nuclei isolation and profiling were placed into cryo-vials, flashed frozen on dry-ice, and stored at -80°C. Most samples for dissociation were processed fresh (below). Samples AT18, AT19, AT27, AT28, AT30 were transferred to cryotubes containing Cryostor CS10 (StemCell Technologies, 07930) and cut into 1 mm pieces with standard laboratory tissue dissection scissors before tubes were transferred to a Mr Frosty slow freezing container (ThermoFisher Scientific, 5100-0036) at -80°C. Samples were then transferred to liquid nitrogen for long-term storage.

Blood was collected from patients within 60 minutes of the start of thyroid surgery. Blood and serum were collected into EDTA and heparin-coated vacutainer tubes (BD Biosciences, 366643 and 668660), respectively, and processed within 2 hours of collection. Serum was centrifuged at 2100g and supernatant was stored at -80°C. PBMCs were isolated from EDTA collection tubes using Ficoll-Paque density gradient centrifugation. PBMCs were resuspended in Cryostor and frozen slowly at - 80°C in a Mr Frosty before being transferred to liquid nitrogen for long-term storage.

Thyroid function testing on serum samples at the Brigham Research Assay Core lab. Chemiluminescence assays were performed to measure TSH (Access HYPERsensitivy hTSH assay, Beckman Coulter), free thyroxine (Access Free T4, Beckman Coulter), anti-TPO antibodies (Access TPO Antibody assay, Beckman Coulter), and anti-thyroglobulin antibodies (Access Thyroglobulin Antibody II, Beckman Coulter).

### Isolation and of thyroid-associated immune cells for 10x scRNA-sequencing

Specimens of thyroid tissue were collected as indicated above. Fresh tissue was washed 5 times in PBS and transferred to HBSS without calcium or magnesium (Thermo Fisher Scientific 14170112) and cut into 1 mm pieces with standard laboratory tissue dissection scissors; samples that were frozen in Cryostor were washed 3 times in HBSS to remove any residual cryostor and resuspended in HBSS. Enzymatic cocktail was then added to the following final concentrations: 225U/mL collagenase type I (Worthington LS004196), 225U/mL collagenase type IV (Worthington LS004188), 5U/mL dispase (Corning 354235), 1mg/mL Pronase (Sigma-Aldrich 10165921001), 0.1 mg/mL DNAse (Roche 10104159001). The kinase inhibitor Y27632 was added to a final concentration of 10µM. Tubes containing tissue fragments in the enzymatic cocktail were placed in a heating block at 37° C with shaking at 500 RPM for 30 minutes with the machine placed on its side to prevent tissue fragments from settling. Following enzymatic digestion, the sample was mechanically disrupted by passing through a 100 µm mesh filter. Following centrifugation at 350 x g for 7 minutes, the supernatant was removed, and RBC lysis was performed for 1 minute (ACK lysing buffer, Lonza, Basel, Switzerland). Cells were washed and resuspended in DMD with 10% (v/v) human AB serum and 10uM Y27632.

For samples without CITE-seq, cells were resuspended in DMEM with 10% (v/v) human AB serum with 10uM Y27632 and were incubated on ice for 30 minutes with the following antibodies: CD66b-FITC (1:70, Biolegend 984102), CD45-PE (1:50, Biolegend 393412), and CD235a PE-Cy5 (1:50, Biolegend 306606). Cells were washed once and resuspended in phenol-free DMEM with 10% (v/v) human AB serum and 10uM Y27632 containing DAPI (ThermoFisher 62248). Single-color controls were performed with these same antibodies using BD CompBeads (BD Biosciences 552843). Gating strategy is shown in Supplementary Fig. 1H. CD45^+^ cells were sorted on a Sony MA900 or BD FACSAria Fusion cell sorter. Data were analyzed using FlowJo 10.7.2 (BD Life Sciences). Isolated CD45^+^ cells were centrifuged and resuspended at a concentration of ∼1000 cells/µl in DMEM. Single cell suspensions containing up to 10,000 cells were loaded per channel in 10x Chromium chips. Cell/bead emulsion and subsequent cDNA libraries were generated using Chromium Single Cell 5’ V1.1 kits (10x Genomics PN-1000165) according to the manufacturer’s instructions. TCR-enriched and BCR-enriched cDNA libraries were generated with the Chromium Single Cell V(D)J Enrichment kit (10x Genomics PN-1000005, PN-1000016, respectively). Library quality was assessed with an Agilent 2100 Bioanalyzer.

Samples with CITE-seq were processed as described above except that following RBC lysis, samples were resuspended in DMEM with 10% (v/v) human AB serum, 10uM Y27632, 2mM EDTA and TruStain FcX blocker (Biolegend 422302). Sample was split into 5 equal aliquots and each aliquot was stained as above with the addition to Total-SeqC cell hashing antibodies (Biolegend). Samples were then washed 3 times with DMEM with 10% (v/v) human AB serum, 10uM Y27632 and 2mM EDTA and sorted as above, ensuring that equal number of cells from each hashing aliquot were sorted. TruStain FcX blocker was added, and sorted cells were stained with TotalSeq-C CITE-seq antibody cocktail (Biolegend). Samples were then washed twice with DMEM with 10% (v/v) human AB serum, 10uM Y27632 and 2mM EDTA, then twice with DMEM with 10% (v/v) human AB serum and 10uM Y27632. Subsequent steps were performed as above, except that up to 50,000 cells were loaded per channel and single cell libraries were generated with the Chromium Single Cell 5’ kit (V1.1, 10x Genomics PN-1000165) together with the 5’ Feature Barcode library kit (10x Genomics PN-1000080) (Supplementary Tables X).

### Isolation of single nuclei from thyroid surgical samples for 10x snRNA-sequencing

Nuclei were extracted from flash frozen thyroid surgical specimens using dounce homogenization and lysis in a tween-based buffer as previously described^45^. In brief, specimens were thawed, rinsed in PBS, chopped into 1-2mm fragments with surgical scissors and homogenized with a 2 ml Dounce Tissue Grinder (Sigma D8938) in lysis buffer containing 10% (w/v) Tween, ST buffer (146 mM NaCl, 1 mM CaCl_2_, 21mM MgCl_2_, 10 mM Tris-HCl pH 9), and RNAse inhibitor (Takara 2313, diluted 1:1000). Homogenate was then filtered through a 35µm filter, centrifuged at 500 x *g*, resuspended in ST buffer with RNAse inhibitor, and filtered through a 10µm filter. Nuclei were counted with bright field microscopy and resuspended at ∼1000 nuclei/µL in ST buffer. Single nuclei suspensions containing up to 8,000 nuclei were loaded per channel in 10x Chromium chips using Chromium Single Cell 5’ V1.1 kits (10x Genomics PN-1000165) and subsequent steps were performed according to the manufacturer’s instructions.

### PBMC CD45^+^ enrichment, cell hashing, and CITE-seq staining

Cryopreserved PBMC samples were thawed at 37°C, diluted with a 10x volume of RPMI with 10% heat-inactivated human AB serum (Sigma), and centrifuged at 300 x g for 7 minutes. Cells were resuspended in CITE-seq buffer (RPMI with 2.5% (v/v) human AB serum and 2mM EDTA) and added to 96-well plates. Dead cells were removed with an Annexin-V-conjugated bead kit (Stemcell 17899) and red blood cells were removed with a glycophorin A-based antibody kit (Stemcell 01738); modifications were made to manufacturer’s protocols for each to accommodate a sample volume of 150 uL. Cells were quantified with an automated cell counter (Bio-Rad TC20), after which 2.5 x 10^5^ cells were resuspended in CITE-seq buffer containing TruStain FcX blocker (Biolegend 422302) and MojoSort CD45 Nanobeads (Biolegend 480030). Live cells were counted with trypan blue, and 6-8 samples (each sample with 60,000 cells) were pooled together at equal concentrations. Pooled samples were filtered with 40 μM strainers, centrifuged, and resuspended in CITE-seq buffer with TotalSeq-C antibody cocktail (Biolegend; Supplementary Table 27). Cells were incubated on ice for 30 minutes, followed by three washes with CITE-seq buffer and a final wash in the same buffer without EDTA. Cells were resuspended in this buffer without EDTA, filtered a second time, and counted.

Hashed PBMC single cell libraries were generated with the Chromium Single Cell 5’ kit (V1.1, 10x Genomics PN-1000165) together with the 5’ Feature Barcode library kit (10x Genomics PN-1000080) (Supplementary Tables X). TCR-enriched and BCR-enriched cDNA libraries were generated with the Chromium Single Cell V(D)J Enrichment kit (10x Genomics PN-1000005, PN-1000016, respectively). Library quality was assessed with an Agilent 2100 Bioanalyzer.

### scRNA-seq data generation

Samples were sequenced on either an Illumina Nextseq 500/550 instrument or an Illumina NovaSeq instrument (SupplementaryTable 2). For Nextseq, gene expression libraries were sequenced using the high output v2.5 75 cycles kit with the following sequencing parameters: read 1 = 26; read 2 = 56; index 1 = 8; index 2 =0. TCR-enriched libraries were sequenced using the high output v2.5 150 cycle kit with the following sequencing parameters: read 1= 30, read 2 = 130, index 1 = 8, index 2 = 0. For Novaseq, all libraries were sequenced using the S4 300 cycles flow with the following sequencing parameters: read 1 = 26 ; read 2 = 91 ; index 1 = 8; index 2 =0.

### scRNA-seq and sn-RNAseq read alignment and quantification

#### RNA data

Raw sequencing data was pre-processed with CellRanger (v6.0.1, 10x Genomics) to demultiplex FASTQ reads, align reads to the human reference genome (GRCh38), and count unique molecular identifiers (UMI) to produce a cell x gene count matrix^46^. We used the Terra platform (https://app.terra.bio/) to run Cell Ranger via the workflow script that is part of a collection called Cumulus^47^. Blood samples were genetically demultiplexed using Souporcell^48^ (v2.0). Hashtag oligos (HTOs) were demultiplexed by Pegasus (v1.7.1, python) functions estimate_background_probs and demultiplex. To ensure robustness when demultiplexing cells, we ran Souporcell with two different set-up. Besides running Souporcell on the raw BAM output from Cellranger, we implemented a semi-supervised run of Souporcell, where pooled PBMC cells are merged with thyroid-associated CD45+ cells of known donors that were run on separate channels. Souporcell assignments between two separate runs were compared, and the identity of the assignments were mapped using a majority vote of cells with known labels.

All count matrices were then aggregated with Pegasus using the aggregate_matrices function. Low-quality droplets were filtered out of the matrix prior to proceeding with downstream analyses using the percent of mitochondrial transcripts and number of unique genes detected as filters (>300 unique genes, <20% mitochondrial content for scRNAseq and <5% mitochondrial content for snRNAseq). The percent of mitochondrial transcripts was computed using 13 mitochondrial genes (*MT-ND6, MT-CO2, MT-CYB, MT-ND2, MT-ND5, MT-CO1, MT-ND3, MT-ND4, MT-ND1, MT-ATP6, MT-CO3, MT-ND4L, MT-ATP8*) using the qc_metrics function in Pegasus. The counts for each remaining cell in the matrix were then log-normalized by computing the log1p (counts per 100,000), which we refer to in the text and figures as log(CPM).

#### Protein data

Raw sequencing data was pre-processed with CellRanger (v6.0.1, 10x Genomics). Antibody-derived tags (ADT) data was demultiplexed by matching cell barcodes of the protein data to that of the RNA data cell barcodes. Only cells that had RNA data were paired with the protein data - we did not include cells that had only protein data without a matching RNA data. Protein data demultiplexing was therefore derived from the RNA-based demultiplexing. All protein values were CLR (centered log ratio) transformed^49^ by the following formula, where x denotes an ADT vector of a single cell c and p is the value of a single protein expression in x:

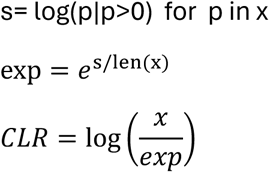

### Lineage-level single-cell and single-nuclei clustering

To identify different cell populations in the thyroid and blood samples we clustered the 3 datasets we profiled separately: snRNAseq on thyroid samples (including epithelial, stromal, endothelial and immune cells from the thyroid), thyroid scRNAseq cells (CD45^+^ immune cells from the thyroid) and PBMC scRNAseq.

Each clustering analysis included the following steps: we identified robust genes as those that are expressed in at least 0.05% of the cells. Among the robust genes, 2,000 highly variable genes (HVGs) were selected using the highly variable features function in Pegasus and used as input for principal component analysis. TCR and BCR genes were pre-excluded from the candidate HVGs. To account for batch effects, the resulting principal component scores were aligned using the Harmony algorithm^50^. The Harmony-adjusted principal components were used as input for Leiden clustering and Uniform Manifold Approximation and Projection (UMAP) algorithm. We identified main lineages using marker genes: T/NK cells (*CD3E, CD3D, CD8A, CD8B, CD4, NCAM1, FCGR3A*), B and plasma cells (*MS4A1, CD19, JCHAIN*), myeloid cells (*VCAN, LYZ, CD136, MARCO, CD14, HLA-DRB1, HLA-DRA*), thyrocytes (*TSHR, TPO, PAX8*), stromal cells (*COL3A1, COL3A2, LAMA2*) and endothelial cells (*PECAM1*). Based on the expression of these markers we picked the following leiden resolutions: main snRNAseq resolution=0.8, main scRNAseq thyroid resolution=1.5, blood scRNAseq resolution=1.5. For every lineage we identified doublets as clusters that express more than one canonical lineage marker. We annotated all the clusters with the same lineage marker expression as being part of that lineage and continued sub-clustering every lineage separately.

#### Sub clustering lineages of thyroid samples

For every lineage separately we re-ran the clustering procedure: identified 2,000 HVGs, ran PCA, computed a UMAP embedding based on 50 PCs, and clustered using the Leiden algorithm. For lineages that were part of the snRNAseq dataset we removed doublets and reran the clustering pipeline in order to be left with a “clean” clustering. The final resolutions for snRNAseq lineage clustering were: epithelial nuclei resolution=1.2, stromal nuclei resolution=1.2, lymphatic nuclei resolution=0.8, vascular nuclei resolution=1.5. For scRNAseq: B, CD4 T and CD8 T cell lineages were clustered and sub-clustered in a 2 step clustering as detailed in Supplementary Note. In these sub-clustering 1,000 HVG and 40 PCs were used to create the embeddings. All final clusters from all sub-clustering were projected back to the original embeddings.

Protein data from the 9 samples that had CITEseq was aggregated. Since these samples were hashed, doublets were identified using the hashtags by Pegasus’ functions estimate_background_probs() and demultiplex(). We used only singlets for downstream analysis and used the protein data by projecting it onto the RNA-based UMAP.

#### Sub clustering lineages of blood samples

When integrating blood samples from all the batches together, samples from Plate 2 5P introduced a batch effect and contained more doublets compared to the rest of the batches. We noted that the batch effect is resolved when doublets were removed from Plate 2 and integration of cells was done at the lineage level (see Supplementary Note). We therefore clustered Plate 2 5P samples separately, identified the main lineages, and merged every lineage with the lineages defined from the remaining batches. For each lineage separately, we re-ran the clustering procedure by identifying 1,000 HVGS, running PCA, removing batch effect with Harmony, computing a neighborhood graph and an UMAP embedding based on 30 PCs, and clustering using the Leiden algorithm. We used the following resolutions for each lineage: myeloid res=1.5, B-Plasma res=1.5, CD8 T cells res=1.8, CD4 T cells res=1.1, and NK-ILC res=1.1.

### Single-cell subset annotation

To annotate cell subsets at the lineage-level clustering, we used two complementary methods to identify marker genes and proteins: “one versus all” (OVA) in which we test which genes are differentially expressed between each cluster and the rest of the cells that do not belong to it; and “all versus all” (AVA) in which we test which genes are differentially expressed between every pair of clusters. For OVA, we calculated the area under the receiver operating characteristic (AUROC) curve for the log(CPM) values of each gene as a predictor of cluster membership using the de_analysis function in Pegasus. Genes with an AUROC ≥0.75 were considered as marker genes for a particular cell subset. In addition, we computed a pseudo-bulk count matrix by counting UMIs for every patient in every cluster and running differential expression analysis on pseudo-bulk normalized counts. Pseudobulk counts were computed only for samples with a minimum of 20 cells per cluster per sample. For AVA, we used the pseudo-bulk matrix to model the gene expression in each cluster separately by the formula gene ∼ clust using the lmFit() function from the Limma R package^51^, where clust denotes the cluster, we then contrasted every cluster with every other cluster in this model using Limma’s contrasts.fit() function. Marker genes for each cell subset were interrogated and investigated for several different cluster resolutions using Cell Guide^52^. For each lineage, we identified the final resolution in which every cluster has a unique set of marker genes differentiating it from the rest of the clusters. If two cell clusters were indistinguishable, they were merged to create one cell subset to obtain the best possible resolution.

### Single-cell diaerential abundance analysis

To test for differential abundances in different cell subsets across conditions, we computed the abundance of each cell subset in every patient and tested differential abundance by Mann-Whitney U test across desired conditions. We then corrected for multiple hypotheses using Benjamini-Hochberg FDR for every lineage / sub-lineage separately.

### Single-cell Diaerential Expression analysis

We test for differentially expressed genes for each cell subset in two different approaches: categorical approach - between high immune infiltration and low immune infiltration sample groups; continuous approach - using a regression model *gene ∼ imm_inf* where *imm_inf* is the immune infiltration frequency per patient. Differential testing was done using the R package DESeq2^54^ (version 4.3), and the input raw count matrices were constructed for each sub-lineage using a pseudobulk aggregation strategy using ADPBulk (https://github.com/noamteyssier/adpbulk). DESeq2 p-values were adjusted for multiple hypothesis testing using the Bonferori-Hochberg procedure.

### Repertoire analysis

#### Clone definitions, frequency of clones

A *clone* was defined by concatenating the following sequences that are the output of CellRanger for T cells: TRA_cdr3, TRB_cdr3, TRA_v_gene, TRB_v_gene, TRA_j_gene, TRB_j_gene; and for B cells: IGL_cdr3, IGK_cdr3, IGH_cdr3, IGL_v_gene, IGK_v_gene, IGH_v_gene, IGL_j_gene, IGK_j_gene, IGH_j_gene. Any cell that lacked any one of the above components was defined as having no clone information.

*Clone frequency* is defined as the ratio between the number of single cells with the exact same clone sequence for a patient and the total number of cells with TCR/BCR information for that patient.

An *expanded clone* is defined as a clone with a frequency of at least 1% out of a sample and that appears in at least 5 single cells in that sample.

### Enrichment of expanded clones in cell subsets

To identify enrichment of expanded clones in cell subsets in different conditions, we defined the frequency of expanded clones per cell subset per patient by

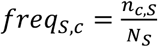

Where

*n_c,s_* is the total number of cells that are part of an expanded clone in sample S in cell subset c

*N_S_* is the total number of cells that are part of an expanded clone in sample S

To test whether cell subsets are enriched with expanded clones we modeled every cluster by the fit *clust ∼ 1+expanded* using the glm() function in R, where ‘clust’ is a binary variable denoting whether a cell is part of the cluster and ‘expanded’ is a binary variable denoting whether a cell is part of an expanded clone. To test whether a cluster is enriched with expanded clones we computed odds ratio (OR) anova between the full model and the null model *clust ∼ 1* using the anova() function in R.

### Comparing clones between tissue and blood and between tissue cell subsets

To test whether tissue cell subsets have similar clones in the blood we modeled every the tissue cell subsets by *clust ∼ 1 +shared* using the glm() function in R, where ‘clust’ is a binary variable denoting whether a cell is part of the cluster and ‘shared’ is a binary variable denoting whether the clone of the cell is shared with another specific cell subset / with blood cells. To test whether a cluster is enriched with shared clones we computed odds ratio (OR) anova between the full model and the null model *clust ∼ 1* using the anova() function in R.

### Cell-cell interaction analysis

To identify which CD8 T cells interact with immuno-thyrocytes in the IFN-ISG axis we computed the the Spearman correlation between the expression of IFN in CD8 T cell subsets and the expression score of ISGs in immuno-thyrocytes. The p-values of all the correlations were adjusted using Benjamini-Hochberg multiple hypothesis testing method.

To identify cell-cell interactions in an unsupervised manner we downloaded all the ligand-receptor interactions available in OmniPath^53^ (downloaded in January 2023). The initial list included 8,278 ligand-receptor pairs and was filtered by the following criteria to include only highly credible pairs:

1. Remove any interactions where the ‘target’ is not transmembrane.
2. Remove any pair with no references (blank in the n_references column).
3. Remove anything with n_sources < 5.
4. Removed interactions that included HLA-I genes.

We then added 36 immune-related ligand-receptor pairs that were missing from OmniPath. Eventually we were left with 1,609 ligand-receptor pairs on which we started the analysis (Supplementary table 12).

Our criteria for identifying relevant cell-cell interactions (CCIs) between epithelial and T cell subsets was requiring:

1. High expression of the ligand in the T cell population (>1 in pseudo-bulk expression).
2. High expression of the ligand in 5.E_IRTs (>1 in pseudo-bulk expression).
3. Ligand is differentially expressed between 5.E_IRTs and 1.E_classical.

* BioRender.com was used in some illustrations in the figures.

### Imaging

#### Tissue Staining

Formalin-fixed paraffin-embedded (FFPE) tissue sections from 5 HT samples, 3 GD samples, and 3 control samples were stained using a Leica Bond RX automated stainer. Heat mediated antigen retrieval was performed with Bond ER2 solution (Tris-EDTA buffer pH 9.0) for 15 mins. Two staining panels were developed using the RNAscope LS Multiplex Fluorescent Assay Combined with sequential Immunofluorescence protocol from ACD, which uses RNAscope LS 4-Plex Multiplex Fluorescent Kits (ACD catalog #322800, #323830), RNAscope ISH probes, and Opal fluorophores (Akoya Biosciences). For each case, the two panels were stained on serial sections. Both panels included cytokeratin AE1/AE3 antibody (Agilent M351529-2) at 1:100 labeled with Opal 780 (1:100). Both panels also shared one RNA probe channel: Hs-IFNG labeled with Opal 690 (1:500). The “T cell” panel was closely based on a previously published panel^24^ and consisted of 5 additional RNA probes: Hs-CXCL13 with Opal 480 (1:1250), Hs-CD8a with Opal 480 (1:620), and a pool of Hs-CXCL9, Hs-CXCL10, and Hs-CXCL11 with Opal 520 (1:1500). The “Fibroblast” panel consisted of 3 additional RNA probes: Hs-COL1A-pool with Opal 480 (1:750), Hs-CCL19 with Opal 620 (1:750), Hs-CD74 with Opal 520 (1:1500). Tyramide Signal Amplification (TSA) was used to boost Opal fluorophore signal. Fluorophore concentrations were optimized to balance signal intensity across all channels. DAPI was used as a nuclear counterstain.

For each case, a third serial section was stained with hematoxylin and eosin (H&E).

#### Image Acquisition

Whole slide images of H&E stained slides were acquired at 20x magnification (0.52 um/pixel resolution) using a MoticEasyScan Infinity digital pathology scanner.

Whole slide images of fluorescent stained slides were acquired on an Akoya Biosciences PhenoImager HT multi-spectral slide scanner at 20x magnification (0.50 um/pixel resolution). Imaging exposures in each channel were optimized for each slide to avoid pixel saturation or underexposure. Akoya inForm software was used to spectrally separate signals from each fluorophore and to mitigate the effect of native tissue autofluorescence. The spectral library used to unmix the raw images was created using the synthetic Opal spectra and autofluoresce spectra from unstained human tonsil tissue. Unmixed tiles produced by inForm were stitched to reproduce a whole-slide pyramidal TIF file using Indica Labs HALO software.

#### Image Analysis

Quantitative image analysis was performed using the HALO image analysis platform (Indica Labs). Tissue sections were manually annotated to exclude large folds and debris from quantification. The three serial section images (H&E and two ISH-IF) were co-registered and overlaid using HALO. Quality of registration was manually verified to be 50 um or less discrepancy between the same tissue structures.

Two approaches were used for regional characterization of tissue. In the first approach (Figures 5-6), TLS and dense lymphocytic infiltrate regions were manually annotated in the H&E images. These two categories of annotations were transcribed to the co-registered ISH-IF images and dilated by 22um to account for small discrepancies during co-registration and to include cells bordering the regions of interest in analysis. In the second approach (Figure 7), two additional annotation categories were added: sparse infiltrate and normal regions. For each ISH-IF panel, Halo AI DenseNet convolutional neural network classifiers were trained to distinguish between these four categories of annotations. TLS and dense infiltrate regions were combined in this analysis as the classifiers were unable to distinguish reliably between these regions. The classifiers were applied across the tissue sections and the resulting classification annotations were manually validated and modified as needed.

For each ISH-IF panel, staining was quantified on a cellular level in the annotated regions using the HALO FISH-IF module v.2.1.5 and HALO AI module v.3.6.1434. Nuclear detection and cell segmentation were performed using the default AI nuclear segmentation algorithm based on the DAPI channel. Cells were phenotyped as panCK-positive based on signal intensity of the antibody channel within the nuclear and cytoplasmic compartments. Positivity of ISH channels were determined based on signal intensity and dot size within a cell, with brighter and larger dots having more weight. The autofluorescence channel was employed as an exclusion marker to reduce false positives from red blood cells, folds, and debris. Additional cell phenotypes were defined using multiple marker criteria (e.g. panCK^+^*CD74*^+^). Analysis settings were manually validated for each image for quality of cell segmentation and phenotyping. Annotation region areas and total tissue area analyzed were also quantified for each image.

#### Definition and analysis of IFN-γ responsive immunity hubs

Based on a published approach^25^, a tessellation of hexagons (side length = 35 um) was overlaid on tissue images. Hexagon tiles that had no cells were excluded. Touching hexagonal tiles with at least one CXCL9/10/11 positive cells were aggregated into a single “hub.” Cell phenotype counts within each hub were calculated and normalized by the hub area.

#### Analysis of regional cell density

For each sample and for each cell population, we quantify the cell density of distinct tissue regions of interest (ROI) as (n_ROI_/n_sample_)(area_ROI_/area_sample_), where n_ROI_ refers to the number of the cell type in ROI, n_sample_ refers to the total number of the cell type in the sample, area_ROI_ refers to the area of the ROI, and area_sample_ refers to the total tissue area examined for a given sample. Enrichment of a given cell population within a specific region was determined by t-test between cell densities. To compare cell type enrichment in regions of interest (TLS vs. non-TLS, hubs vs. non-hubs, normal vs. sparse vs. dense) p-values computed by two-sided Mann-Whitney paired test.

## DATA AND CODE AVAILABILITY

scRNAseq, snRNAseq, CITE-seq, and repertoire data were deposited to GEO and are available in the accession number <u>GSE271434</u>. The raw human sequencing data will be available in the controlled access repository dbGaP (https://www.ncbi.nlm.nih.gov/gap) upon publication. The source code used for single-cell analysis and figures generation will be available in GitHub at https://github.com/villanilab/Thyroid and in the Zenodo repository upon publication of this study. A full list of software packages and versions included in the analysis can be found in Supplementary Table 26. A user-friendly portal^52^ is available to browse the single-cell data generated in this manuscript at https://villani.mgh.harvard.edu/thyroid (user: “Thyroid”; password “hashimotos”).

## SUPPLEMENTARY FIGURES

**Supplementary Figure 1:**
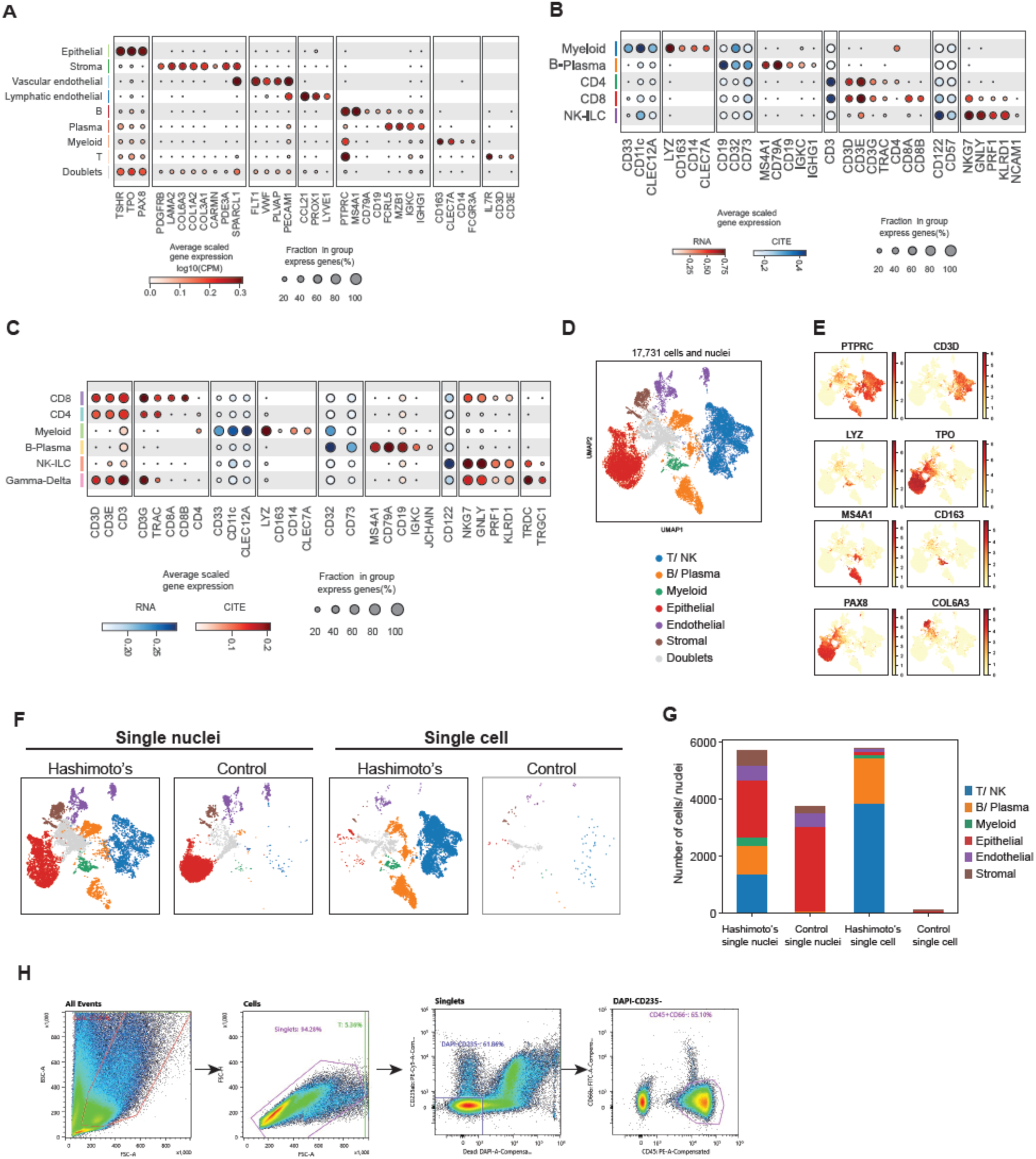
Major cell lineages in thyroid tissue and paired blood lineages in thyroid and blood. **A-C.** Dot plots display marker genes and/or proteins for the cell types indicated. Dot size represents the percent of cells in the population with non-zero expression of a given gene. Color indicates scaled expression of each gene. **A.** Major thyroid cell lineages, profiled by snRNAseq. **B.** CD45^+^ (immune) cells in thyroid, profiled by scRNAseq. **C.** Major peripheral blood mononuclear cell (PBMC) lineages, profiled by scRNAseq. **D-G.** Thyroid tissue specimens from 2 patients (Hashimoto’s and control) were profiled by both snRNAseq and by scRNAseq. **D.** UMAP embedding of cells and nuclei, from both approaches. Color denotes main lineage annotations. **E.** Feature plots of markers used to annotate lineages in **D**. **F.** UMAP embedding for each library. **G.** Bar plot of the number of cells or nuclei from each library (x-axis), colored by lineage. **H.** Fluorescence assisted cell sorting (FACS) strategy for CD45^+^ (immune) cell enrichment from thyroid tissue.

**Supplementary Figure 2:**
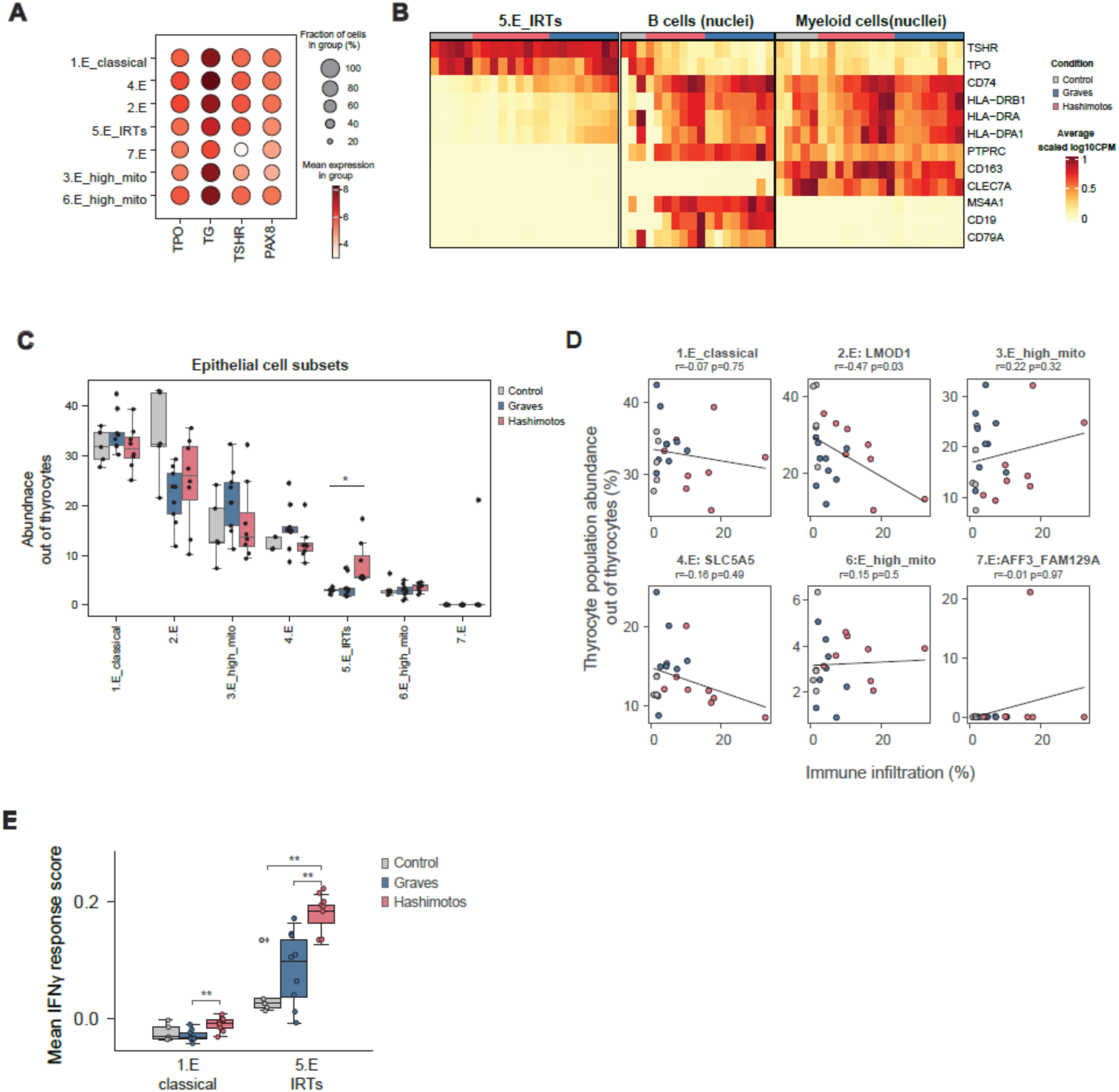
Features of thyrocyte populations. **A.** Dot plot showing expression of genes associated with thyroid hormone production in all thyrocyte populations. **B.** Heatmap displaying expression of markers of thyrocyte identity (*TSHR*, *TPO*), genes encoding components of MHCII (*CD74*, *HLA-DRB1*, *HLA-DRA*, *HLA-DPA1*), immune cell marker (*PTPRC*), and markers of myeloid (*CD163*, *CLEC7A*) and B (*MS4A1*, *CD19*, *CD79A*) cell identity, in E5_IRTs, B cells, and myeloid, based on snRNAseq. Color denotes expression averaged by patient and scaled across epithelial, B and myeloid cells. **C.** Boxplot with frequency of thyrocyte population frequencies. **D.** Abundance of each thyrocyte population (y-axis) versus immune infiltration (x-axis). Spearman correlation and correlation p-value are noted for each thyrocyte population. Lines represent linear regression of the data. **E.** Boxplots quantifying mean IFN-**γ** response Z-score for 1.E_classical and 5.E_IRTs. P-value: ** (<0.01).

**Supplementary Figure 3:**
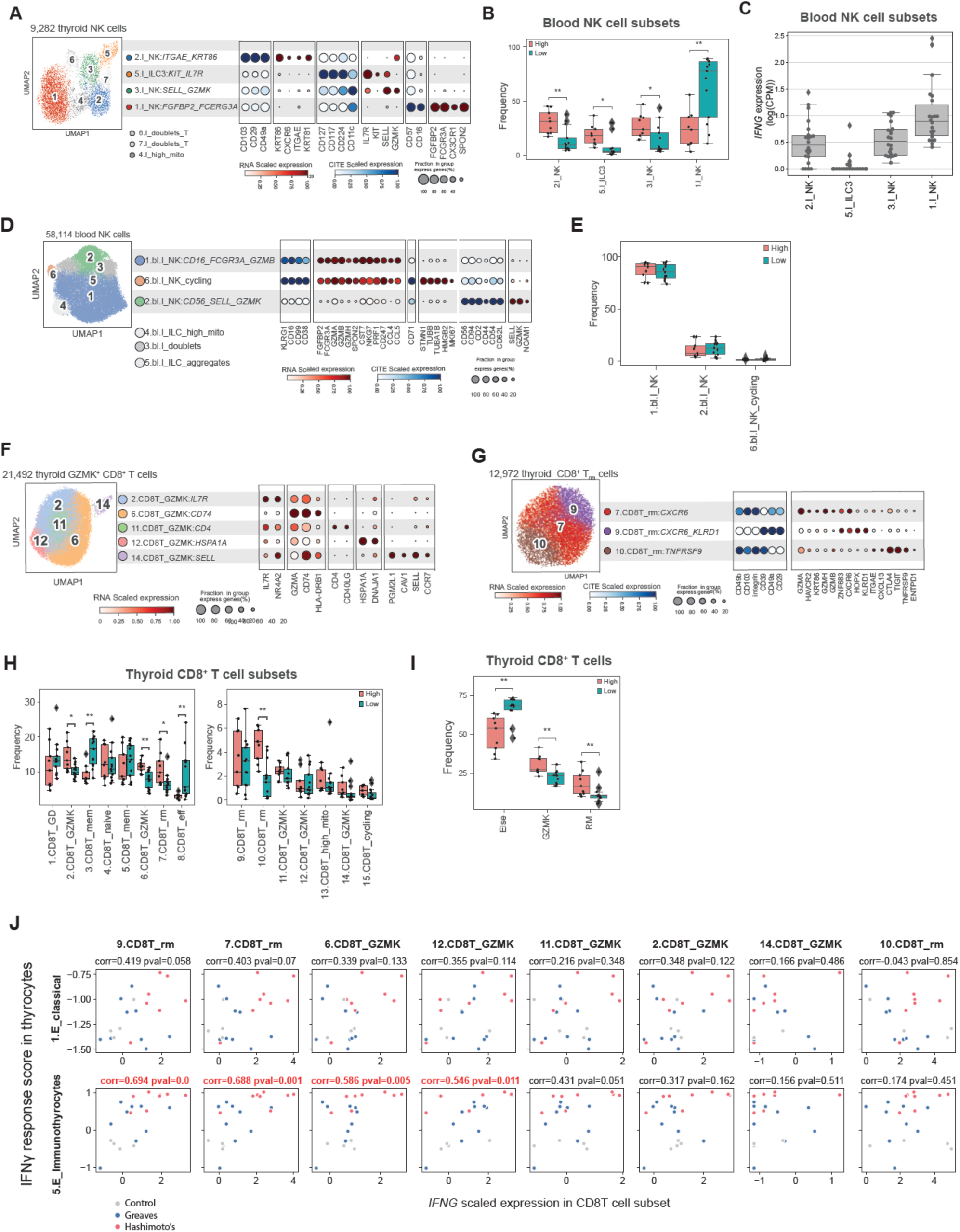
Features of CD8^+^ T cells and NK cells in blood and tissue. **A, D, E:** For dot plots, dot size represents percent of cells in the population with non-zero expression of a given gene or protein. Color indicates scaled expression. **A.** UMAP embedding of thyroid-associated NK and ILC populations (left) and dot plot illustrating main gene and protein expression markers of each population (right). **B.** Boxplot illustrating differential abundance for NK and ILC populations in samples with high versus low immune infiltration. **C.** Boxplot of *IFNG* expression in thyroid-associated NK and ILC populations. **D.** UMAP embedding of thyroid-associated CD8^+^ GZMK^+^ cells (left) and dot plot illustrating main gene expression markers of each population (right). **E.** UMAP embedding of thyroid-associated CD8^+^ T_RM_-like cells (left) and dot plot illustrating main gene expression markers of each population (right). **F.** Boxplot illustrating differential abundance analysis comparing frequencies of thyroid-associated CD8^+^ T cell populations in samples with high versus low immune infiltration. **G.** Boxplot illustrating differential abundance for thyroid-associated CD8^+^ cell populations: GZMK^+^ cells, T_RM_-like cells and other CD8^+^ T cells across high- and low-infiltration samples. **H.** Relationship between *IFNG* expression in CD8^+^ T cell populations (x-axis, each column is a distinct population) and IFN-**γ** response signature in thyrocyte subsets (1.E_classical in the top row, 5.E_IRTs in the bottom row). Spearman correlation and p-value were determined for each T cell/thyrocyte pair. Significant correlations are marked in red.

**Supplementary Figure 4:**
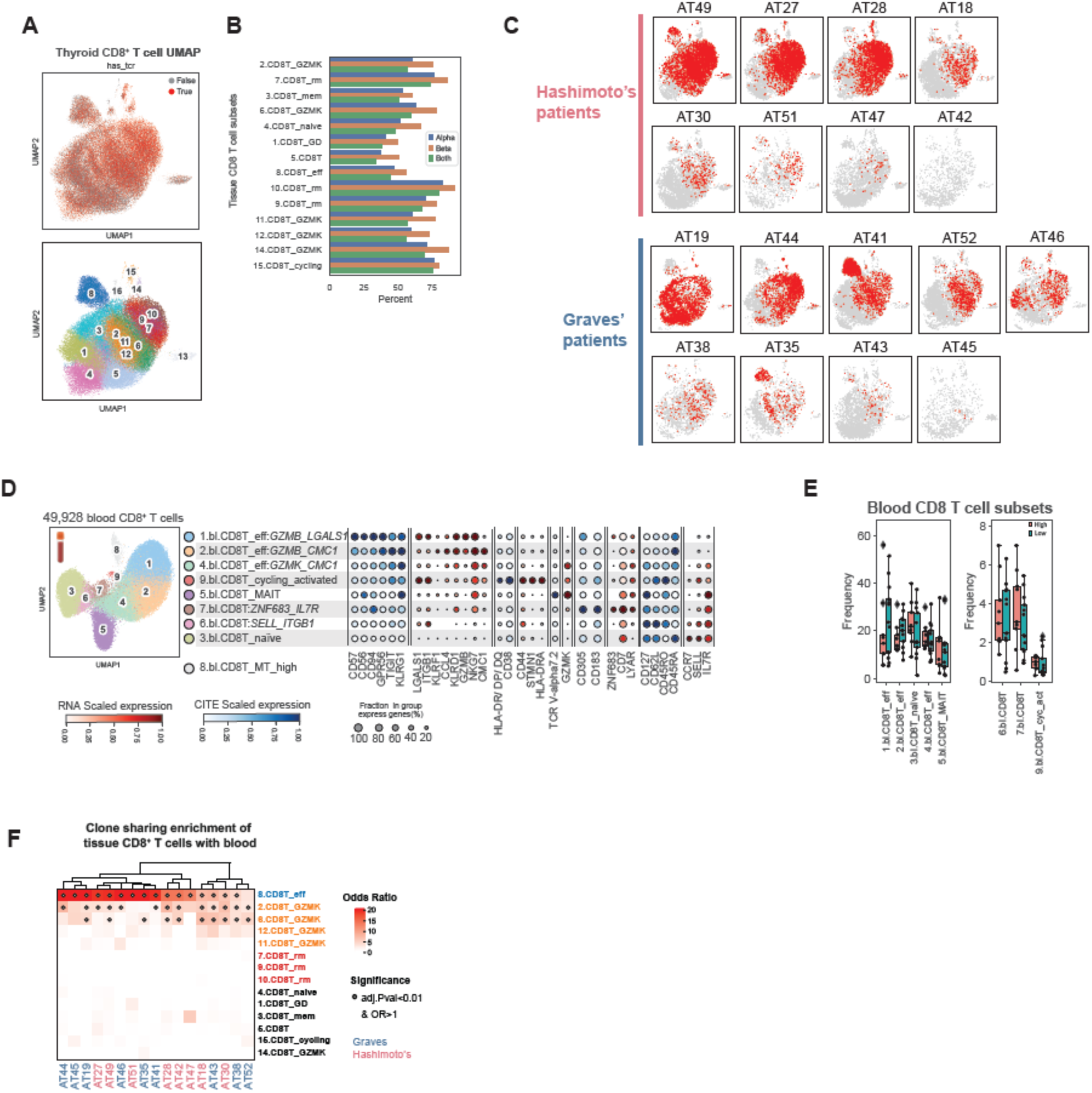
TCR analysis of CD8^+^ T cells. **A, B.** scTCRseq data availability for thyroid-associated CD8^+^ T cells. **A.** UMAP embeddings of CD8^+^ T cells colored by presence of scTCR-seq data (top) and cell populations (bottom). **B.** Bar plot quantifying percent of cells per CD8^+^ T cell population with scTCRseq data. Color denotes the chain type (‘Alpha’ or ‘Beta’ or ‘Both’). Asterisks denote comparisons in which OR>1 and adj. p-value<0.05. **C.** UMAP embedding of CD8^+^ T cells colored based on whether it is part of an expanded CD8^+^ T cell clone. **D.** UMAP embedding of CD8^+^ T cells from blood (left) and dot plot illustrating main gene and protein expression markers of each population (right). Dot size represents percent of cells in the population with non-zero expression of a given gene or protein. Color indicates scaled expression. **E.** Boxplot illustrating differential abundance analysis comparing frequencies of blood CD8^+^ cell populations in samples with high versus low immune infiltration. **F.** Heatmap summarizing the odds ratio (OR) of a logistic regression model that tests the association between clone sharing of each thyroid-associated CD8^+^ T cell population (y-axis) with all blood CD8^+^ T cells per patient (x-axis). Color denotes OR values, dots denote significance. **G.** UMAP embedding of blood NK and ILC populations (left) and dot plot illustrating main gene and protein expression markers of each population (right). Dot size represents percent of cells in the population with non-zero expression of a given gene or protein. Color indicates scaled expression. **H.** Boxplot illustrating differential abundance analysis comparing frequencies of blood NK and ILC populations across in samples with high versus low immune infiltration.

**Supplementary Figure 5:**
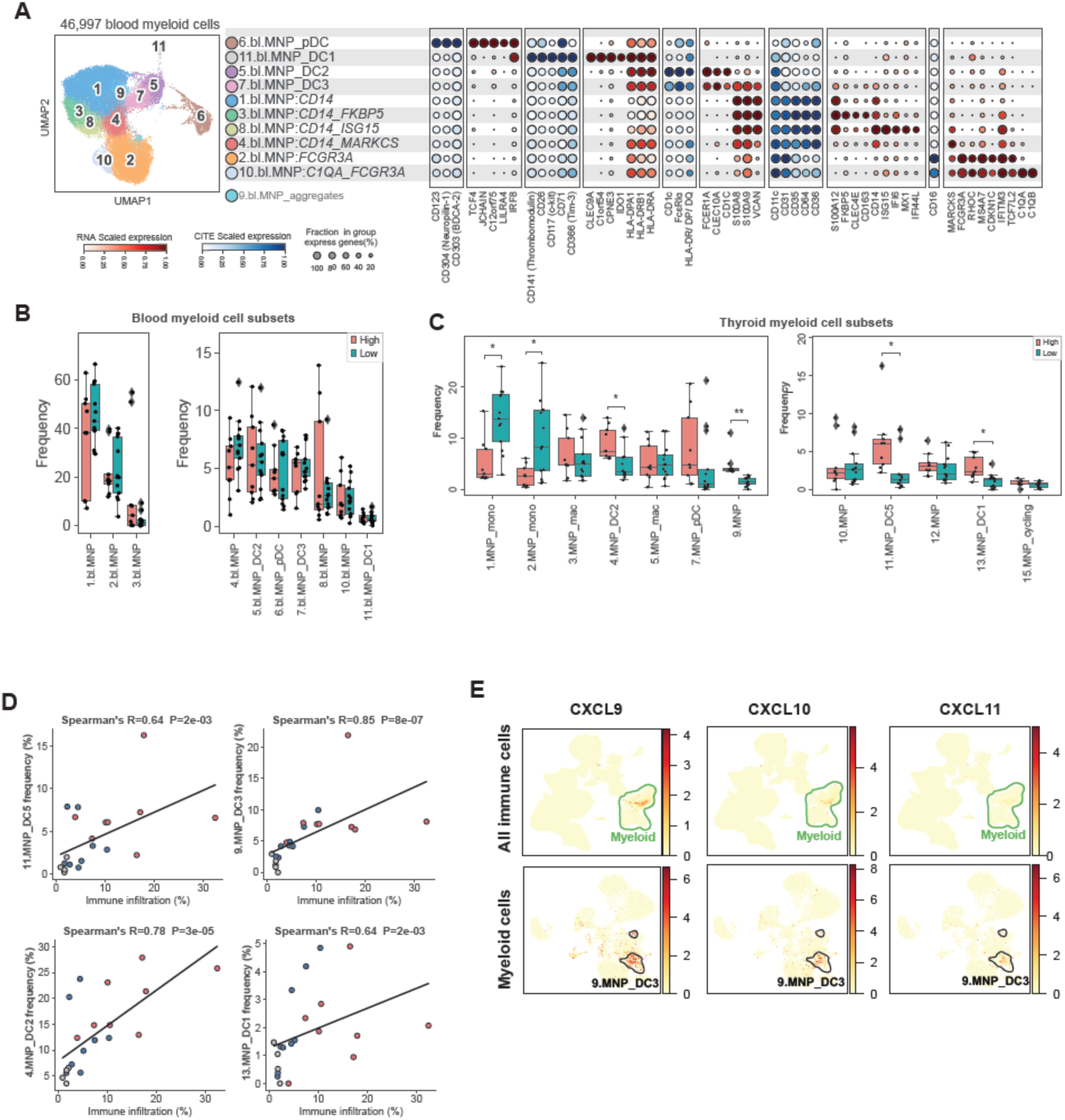
Features of myeloid cells in blood and tissue. **A.** UMAP embedding of myeloid cells in blood (left) and dot plot illustrating main gene and protein expression markers of each population (right). Dot size represents percent of cells in the population with non-zero expression of a given gene or protein. Color indicates scaled expression. **B, C.** Boxplots illustrating differential abundance analysis comparing frequencies of myeloid cell populations in samples from patients with high versus low thyroid immune infiltration. **B.** Myeloid populations in blood. **C.** Myeloid populations in tissue. **D.** Relationship between immune infiltration (x-axis) and frequency of DC cell populations (among all myeloid cells, determined by scRNAseq, y-axis). Lines represent linear regression of the data. **E.** UMAP embeddings of thyroid-associated immune cells (top) and thyroid-associate myeloid cells (bottom) colored by normalized expression of *CXCL9*, *CXCL10* and *CXCL11*.

**Supplementary Figure 6:**
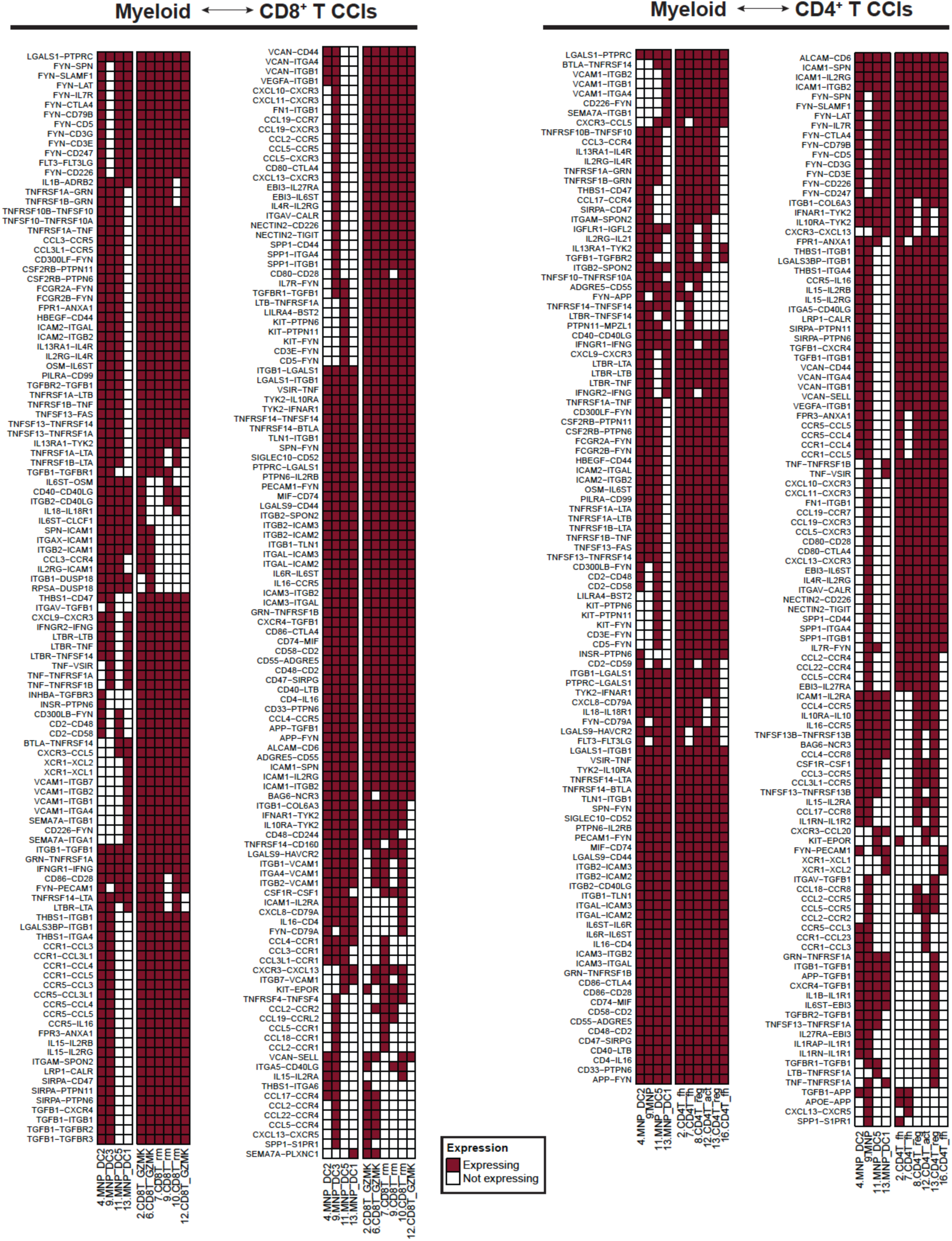
Putative DC-T cell interactions. Each row represents a potential protein-protein interaction between molecules expressed in DC populations (4.MNP_DC2, 9.MNP_DC3, 11.MNP_DC5, 13.MNP_DC1) and T cells (CD4^+^ and CD8^+^). Maroon boxes indicate genes that are expressed; white boxes indicate genes with <1 in pseudo-bulk expression.

**Supplementary Figure 7:**
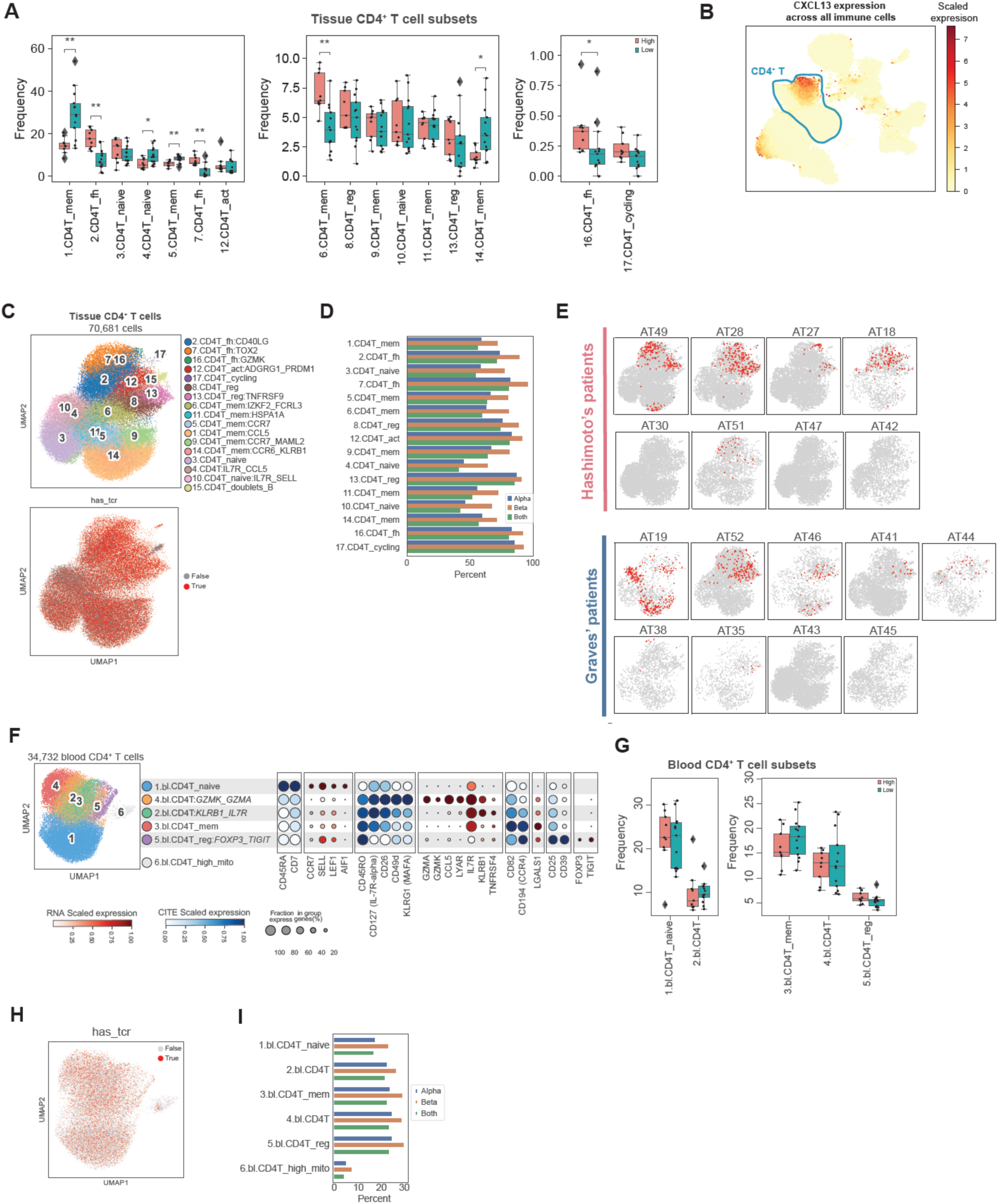
Features of CD4^+^ T cells in blood and tissue. **A.** Boxplot illustrating differential abundance analysis comparing the frequencies of thyroid-associated CD4^+^ T cell populations in samples with high versus low immune infiltration samples. **B.** UMAP embedding of thyroid-associated immune cells, colored by the expression of *CXCL13.* **C.** UMAP embedding of thyroid-associated CD4^+^ T cells, colored by cell populations (top), or by the presence of scTCRseq data (bottom). **D.** Bar plot quantifying percent of cells per CD4^+^ T cell population with scTCRseq data. Color denotes chain type (‘Alpha’,‘Beta’, or ‘Both’). **E.** UMAP embedding per patient of thyroid-associated CD4^+^ T cells for each patient. Red dots indicate cells that are part of an expanded CD4^+^ T cell clone. **F.** UMAP embedding of CD4^+^ T cells in blood (left) and dot plot with main genes and protein markers of each population (right). Dot size represents percent of cells in the population with non-zero expression of a given gene or protein. Color indicates scaled expression. **G.** Boxplot illustrating differential abundance analysis comparing frequencies of blood CD4^+^ T cell populations in samples with high versus low immune infiltration. **H.** UMAP embedding of blood CD4^+^ T cells, colored by presence of scTCRseq data. **I.** Bar plot quantifying percent of cells per blood CD4^+^ T cell population with scTCR-seq data. Color denotes the chain type (‘Alpha’, ‘Beta’, or ‘Both’).

**Supplementary figure 8:**
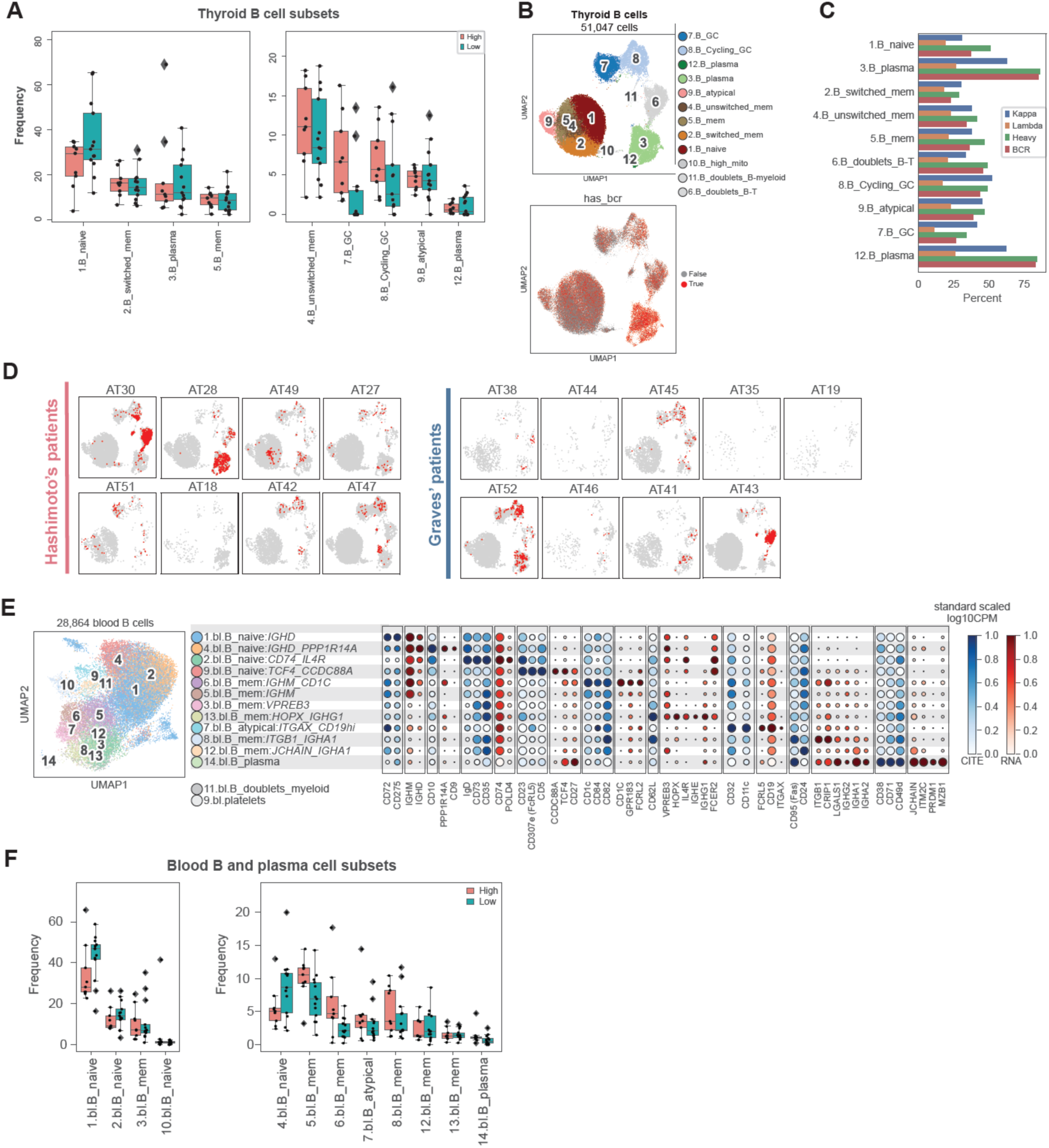
Features of tissue and blood B cells. **A.** Boxplot illustrating differential abundance analysis comparing frequencies of thyroid-associated B cell populations in samples with high versus low immune infiltration. **B.** UMAP embedding of thyroid-associated B cells, colored by cell population (top), or presence of scBCR-seq data (bottom). **C.** Bar plot quantifying percent of cells per thyroid-associated B cell population that have scBCR-seq data. Color denotes the chain type (Kappa’/ ‘Lambda’/ ‘Heavy’) or the availability of data from a heavy chain <u>and</u> either Kappa or Lambda chains (‘Both’). **D.** UMAP embedding of thyroid-associated B cells for each patient. Red dots indicate cells that are part of an expanded B cell clone. **E.** UMAP embedding of B cells in blood (left) and dot plot illustrating main gene and protein expression markers of each population (right). Dot size represents percent of cells in the population with non-zero expression of a given gene or protein. Color indicates scaled expression. **F.** Boxplots illustrate differential abundance analysis comparing the frequencies of blood B cell populations across high- and low-infiltration samples.

**Supplemental Figure 9:**
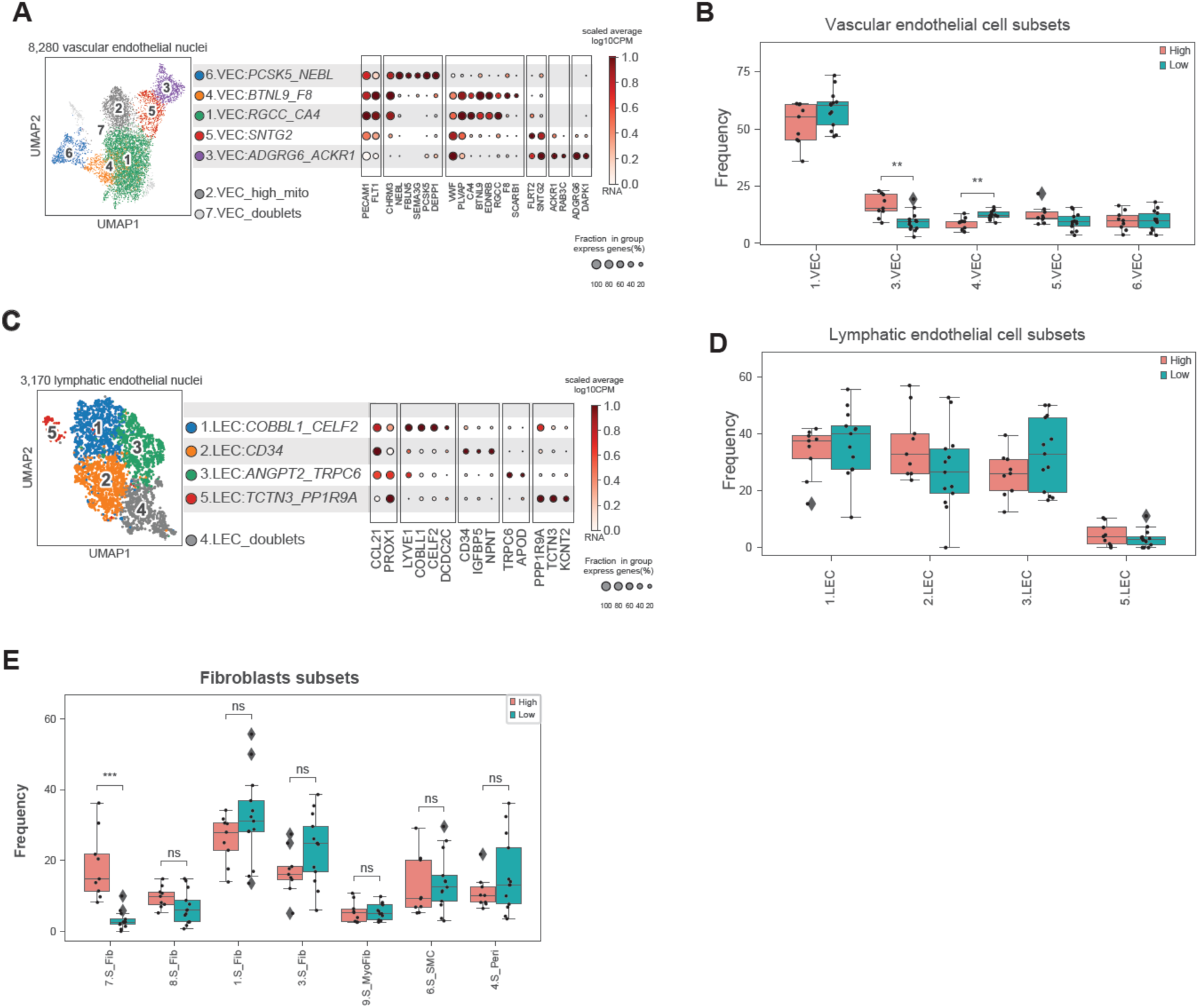
Features of tissue stromal and endothelial cells. **A, C.** For dot plots, dot size represents percent of cells in the population with non-zero expression of a given gene or protein. Color indicates scaled expression. **A.** UMAP embedding of thyroid vascular endothelial cells (VEC) d(left) and dot plot illustrating gene and protein expression markers of each population (right). **B, D, E.** Boxplots illustrate differential abundane analysis, comparing samples with high versus low immune infiltration. **B.** Differential abundance analysis of VEC populations. **C.** UMAP embedding of thyroid lymphatic endothelial cells (LEC) (left) and dot plot illustrating gene and protein expression markers of each population (right). **D.** Differential abundance analysis of LEC populations. **E.** Differential abundance analysis of stromal populations.

**Supplementary Figure 10:**
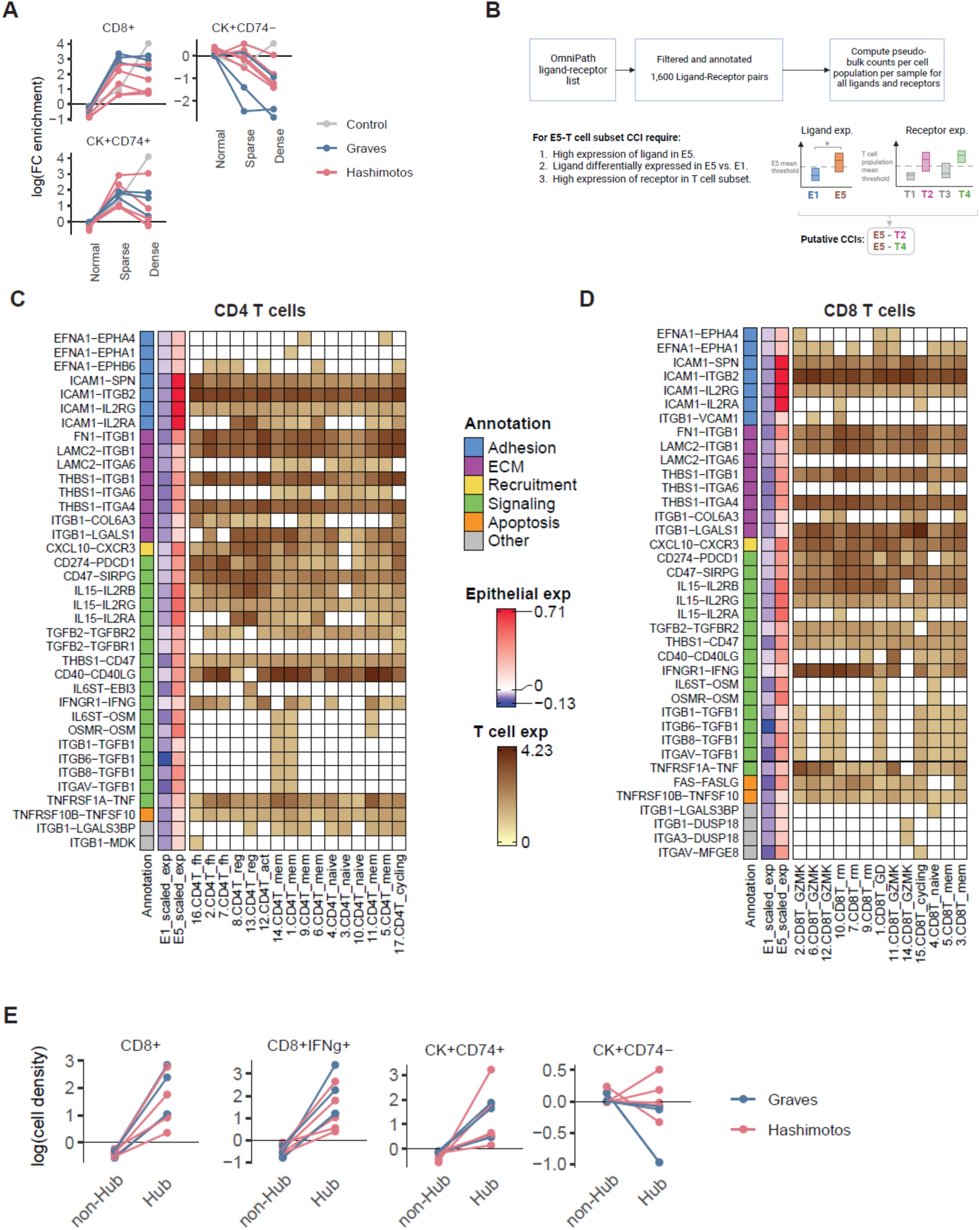
Immune-epithelial cell interactions and immune hub features. **A.** Density of cellular phenotypes in three major macroscopic regions, as shown in Figure 7C, but colored by disease condition. Each pair of dots connected by a line represents a patient, colored by condition. **B.** Schematic of analysis pipeline to identify putative T cell-thyrocyte cell-cell interactions (CCIs). Ligand-receptor pair list was downloaded from OmniPath and filtered to obtain reliable pairs. Pseudo-bulk expression was computed for each ligand and receptor in each CD8^+^ and CD4^+^ T cell subsets as well as 1.E_classical and 5.E_IRTs. CCIs were then filtered to include only those in which one interacting molecule is expressed above a threshold (pseudo-bulk expression>1) in at least one T cell population and its partner is expressed in 5.E_IRTs but not in 1.E_classical. **C, D.** Putative thyrocyte-T cell paracrine interactions. Each row represents a potential protein-protein interaction between molecules expressed in thyrocytes (1.E_classical and 5.E_IRTs) and T cell populations (*CD4*^+^ and *CD8*^+^). Color for thyrocyte columns represents scaled expression across 1.E_classical and 5.E_IRTs. Color in T cell columns represents expression in log(CPM); genes expressing <1 in pseudo-bulk expression are colored white. **E.** Density of cellular phenotypes in three major macroscopic regions, as shown in Figure 7H, but colored by disease condition.

**Supplementary Figure 11:**
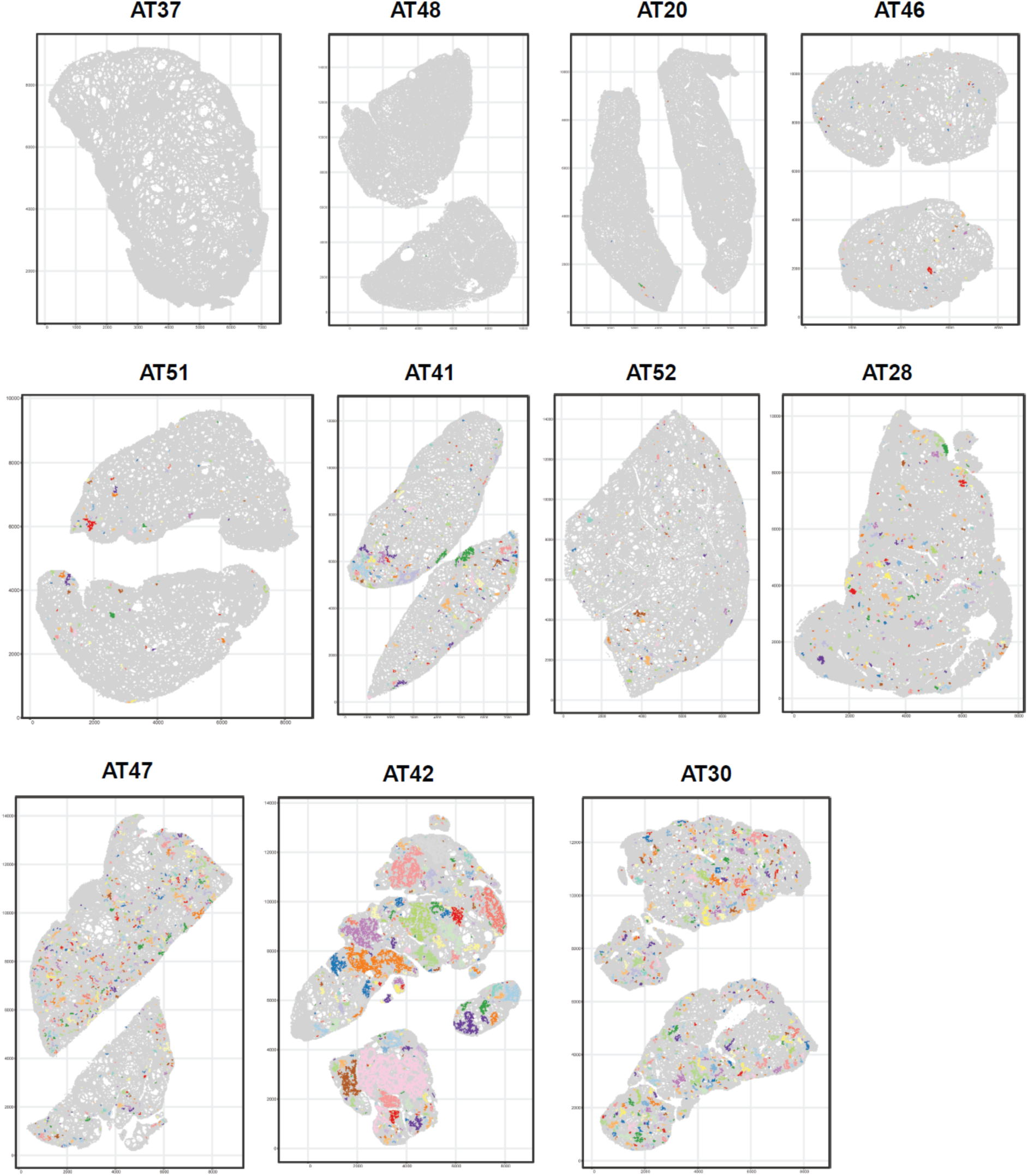
Categorial versus regression analysis of diaerentially expressed genes (DEGs) for all cell populations. Bar plots summarizing the number of significant DEGs found using two different modeling approaches. Data are based on snRNAseq of non-immune cells in tissue (top), scRNAseq of thyroid-associated immune cell populations (middle) and scRNAseq of PBMC populations (bottom). Bars above the x-axis denote the number of DEGs found by regression on immune infiltration. Bars below the x-axis denote the number of DEGs found by comparing samples from patients with HT versus GD. For each approach we summarized the number of DEGs found by a cutoff of FDR=0.05 (red) and FDR=1 (blue).

**Supplementary Figure 12:**
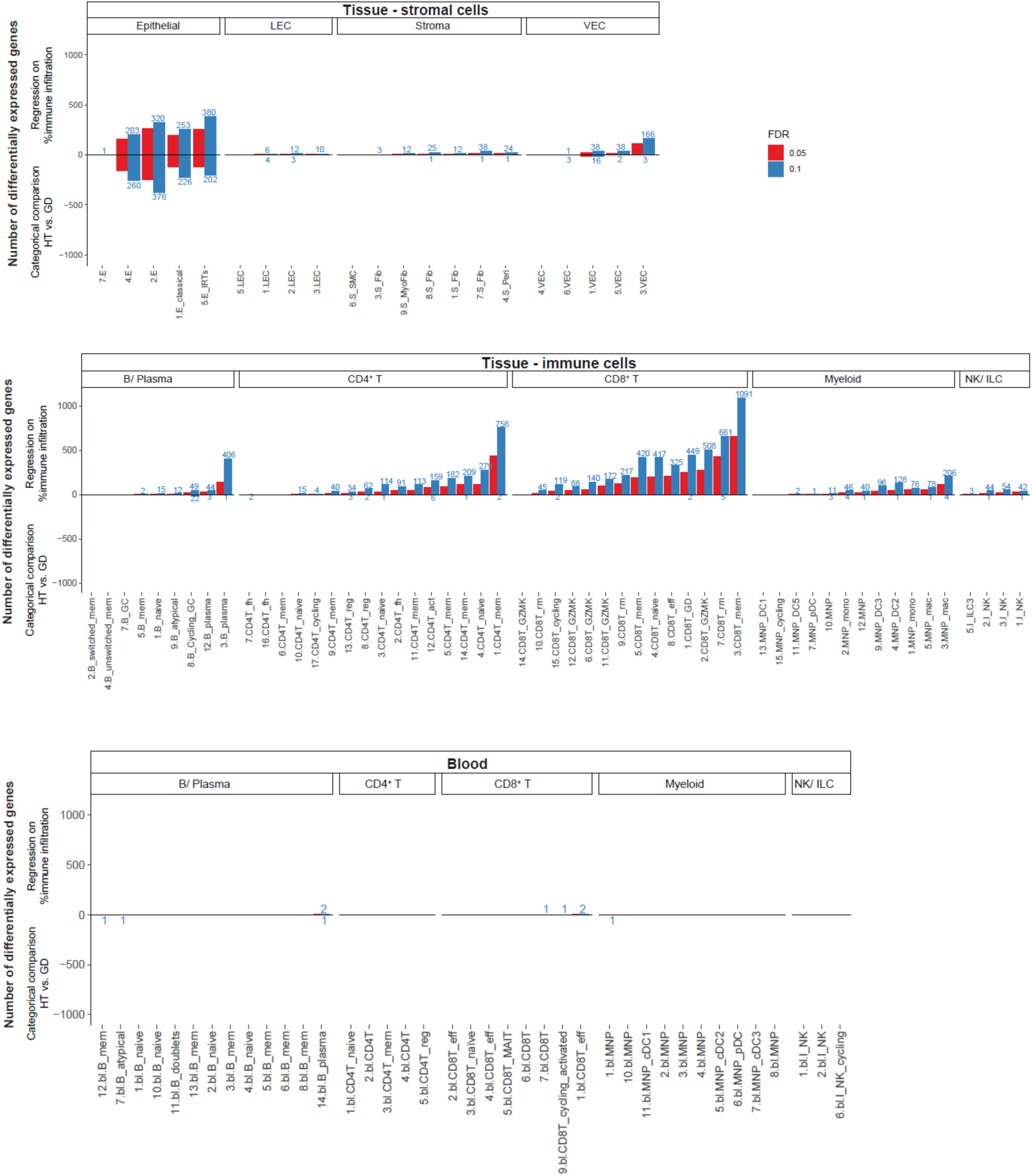
IFN-γ response hubs in tissue specimens. Each image is a thyroid tissue section from a distinct patient converted to a hexagonal tesselation as described in **Methods**. Colored regions indicate an IFN-**γ** response hub, with each color within each patient indicating a distinct hub.

## SUPPLEMENTARY NOTES

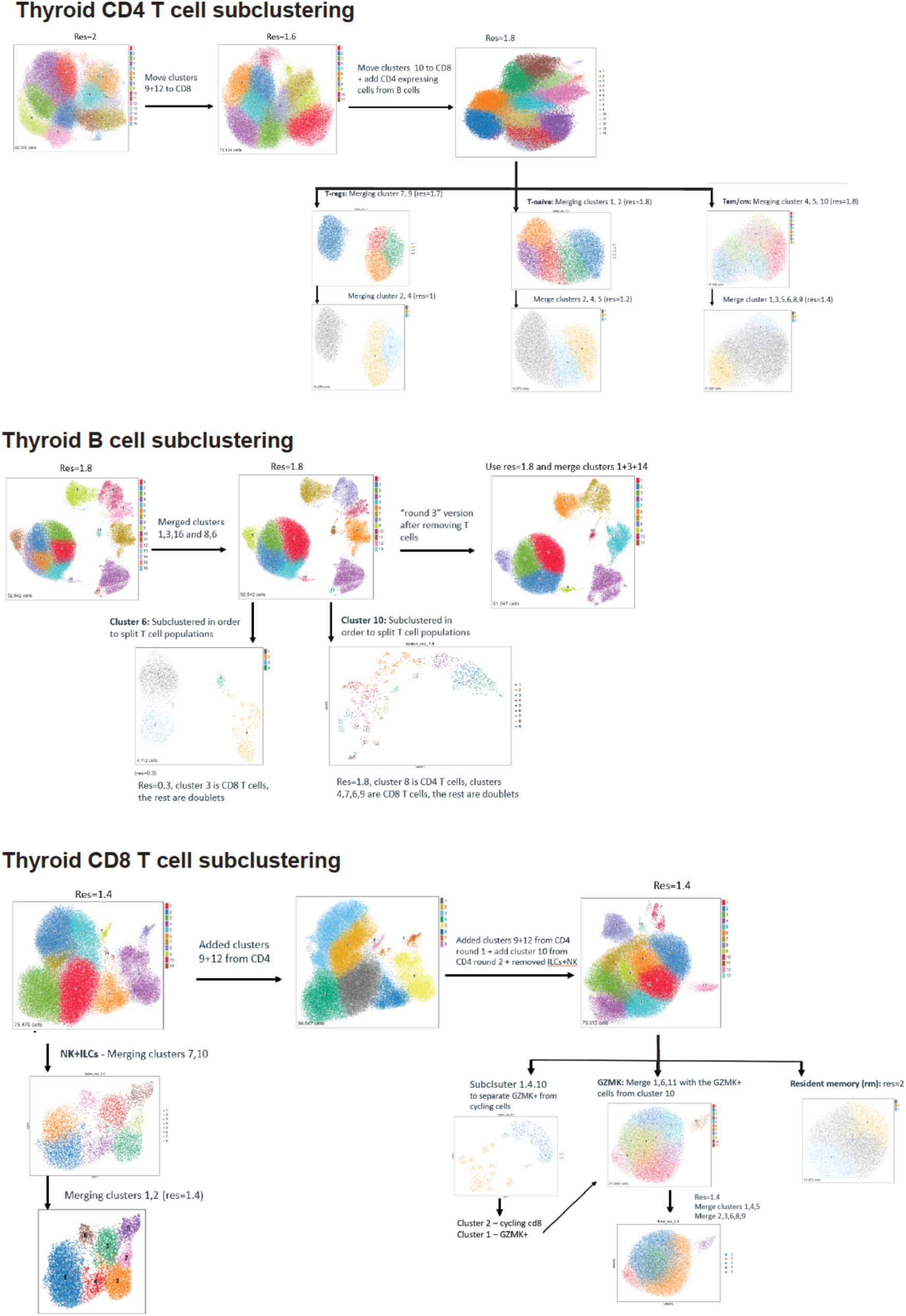

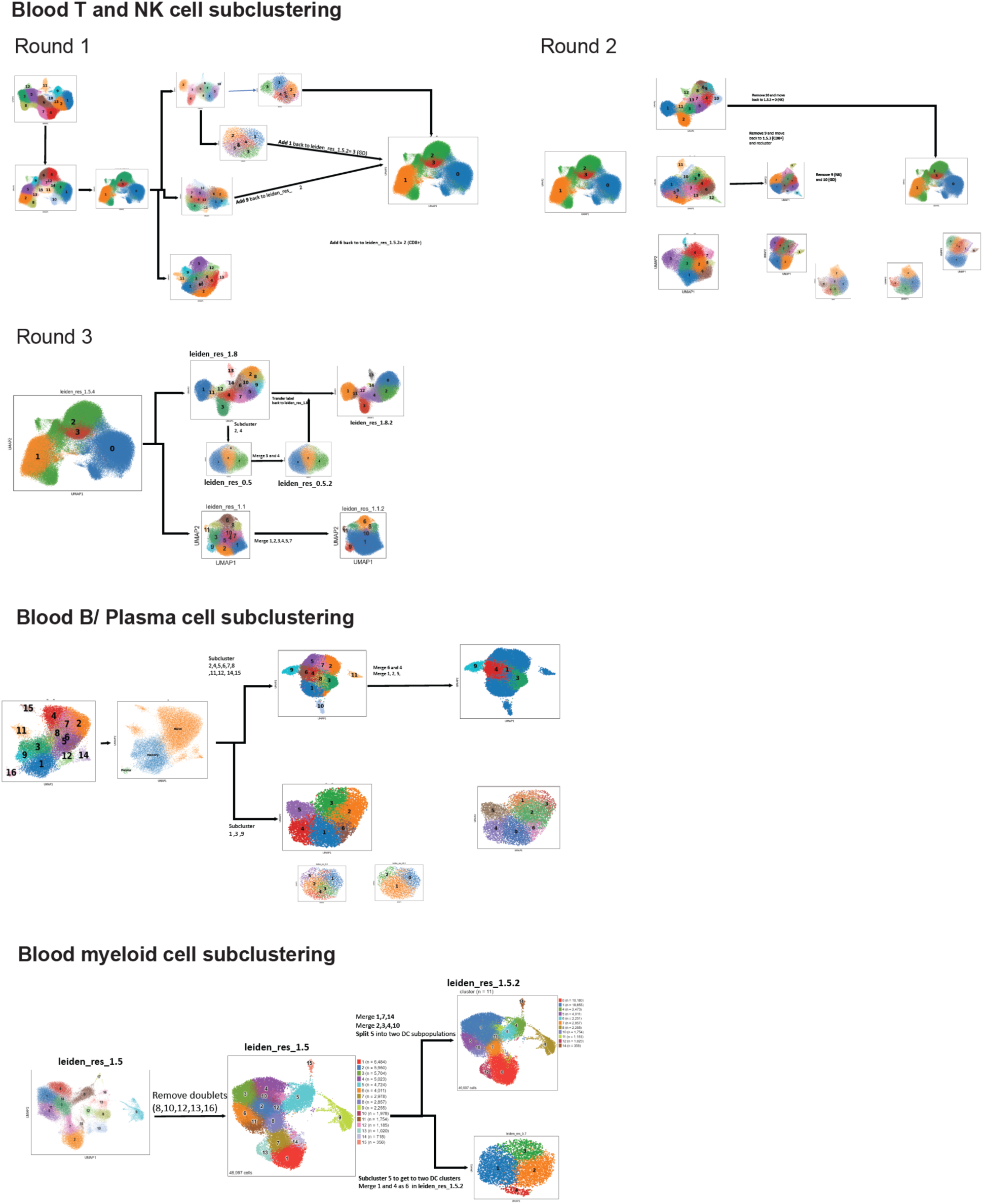

## SUPPLEMENTARY NOTES LEGEND

Illustration of the clustering and subclustering process of different cell lineages as described in the Methods section. For every lineage we detail the clustering resolution we used, which clusters were identified as belonging to another lineage, which clusters were merged, and which were taken for further sub-clustering.

## LIST OF SUPPLEMENTARY TABLES

**Supplementary Table 1. Clinical metadata for patient cohort.** Clinical metadata for donors and patients who contributed tissue or blood for analysis. “Sample ID” denotes patient or donor study identifiers. The table includes clinical metadata, what profilings were done per sample for this study and serum measurements at the day of sampling. In addition, for every patient there are columns denoting the amount of single cell/ single nuclei that were capture for blood and tissue as well as the mean number of genes and the mean percent of mitochondrial content for every sample. **Related to all Figures.**

**Supplementary Table 2. Library information.** Each row represents a unique library sequenced for our dataset. “Library name” denotes a unique library identifier. “Library type” can be one of 4 different values: ‘GEX’ for gene-expression library, ‘FBC’ for CITEseq library, ‘TCR’ for T cell receptor repertoire library and ‘BCR’ for B cell receptor repertoire library. “Chemistry denotes the 10X Genomics chemistry used to generate the library. “Batch” denotes the batch the sample was processed in; a lot of the libraries were prepared individually and run as a ‘single channel’, but when batches were used (blood libraries) we detail a unique batch identifier. “Index” is the unique primer used for multiplexing different libraries in a sequencing run. Finally, “Enrichment” denotes whether the library underwent selection for immune cell enrichment. **Related to all Figures.**

**Supplementary Table 3. Main lineage frequencies and diaerential abundance.** This table includes the frequency of every main cell lineage for every sample, and differential abundance testing per lineage between conditions. The differential abundance section contains 3 different comparisons - Control vs. Hashimoto’s, Graves vs. Hashimoto’s and Control vs. Graves. Each comparison and cell lineage include the mean abundance per condition (“mean1”, “mean2”) the log(fold-change) between the conditions “log(FC)”, the p-value and adjusted p-value (“q-value”) for the Mann-Whitney test. **Related to** Figure 1.

**Supplementary Table 4. Frequencies and diaerential abundance of all cell subsets.** Every row in this table represents a cell subset either in tissue or blood. Columns D-AA include the frequency of the relevant cell subset in a specific donor. Blank cells represent data not available for that patient. Columns AB-AM include differential abundance testing. Columns AB-AD shows that statistics of comparing the abundance in samples with low immune cell infiltration vs. samples with high immune cell infiltration (annotations of the samples to which is ‘low’ and which is ‘high’ can be found under the samples names in columns D-AA. Columns AE-AM include differential abundance between samples from different conditions and shows 3 different comparisons: Control vs. Hashimoto’s, Graves vs. Hashimoto’s and Control vs. Graves. Row 5 includes the of TPO auto-antibody levels per patients ln(TPO_ab +1). **Related to** Figures 2-6.

**Supplementary Table 5. *IFNg* stimulated genes information.** This table includes the IFNg stimulated genes (ISGs) used for ISG expression scores in epithelial cells (‘Gene list’). The table on the right includes the expression levels of this ISG score for every patient in either 1.E_classical or 5.E_IRTs. **Related to** Figures 1**, 2.**

**Supplementary Table 6. List of dysfunction and cytotoxic genes** used to measure functionality of T cell populations. **Related to** Figures 3**, 7.**

**Supplementary Table 7. Cell-cell interactions between epithelial and CD8+ T cells.** This table includes the correlations between *IFNg* expression in CD8T+ cell subsets (‘Sender_population’) and ISGs expression in epithelial cell subsets (‘Reciever_population’). For every cell subset pair there is the spearman correlation (‘correlation’), the p-value and adjusted p-value for that correlation (‘P-value’, ‘adj_P-value’) and the -log(adj_pvalue) that was used for plotting. **Related to** Figure 2.

**Supplementary Table 8. CD8T and CD4T clones.** This table includes all the TCR clones of CD8T+ (left table) and CD4+ (right table) T cells in the thyroid tissue. Every row is a different single cell that has clone information from TCR/ BCR VDJ sequencing. Every single cell includes information about which donor it came from (‘sample’); the clone itself (‘clone’), which is concatenation of: CDR3, V gene and J gene nucleic acid sequences from TCR-alpha, TCR-ꞵ; The cluster it is part of (‘Final_cluster’); Is the clone expanded (‘Is clone expanded?’) and is it shared with the blood in even 1 copy (‘Is clone shared with blood?’). **Related to** Figure 3**, 5.**

**Supplementary Table 9. Thyroid CD8 T cell and CD4 T cell expanded clones enrichment.** In this table we summarize the enrichment of expanded clones in the different CD4+ T cell subsets (right table) and CD8+ T cell subsets (left table). The enrichment was computed by computing an odds ratio (OR) test between 2 regression models: *clust ∼ 1 +expanded* and *clust ∼ 1* (see Methods). Every table includes the cell subset name (‘CD4T cell subset’ / ‘CD8T cell subset’), the model’s p-value (‘model.pvalue’) and the adjusted p-value (‘model.padj’) and the OR (‘expanded.OR’).

**Supplementary Table 10. Shared clones between blood and tissue for CD4+ and CD8+ T cell subsets**. This table contains the logistic regression model results for testing enrichment of tissue cell subsets with clones that are shared with the blood. Two models were compared and an odds ratio (OR) was computed to detect which better describes the data: *clust ∼ 1 +shared* and *clust ∼ 1.* See Methods for more details. The table includes the cell subset in teh model (‘clust’ in the model, ‘CD8/CD4 T cell subset’ in the table), the sample, the model p-value (‘model.pvalue’) and adjusted p-value (‘model.padj’) and the OR (‘Shared.OR’). **Related to** Figures 3**, 5**.

**Supplementary Table 11. CXCL9/10/11 expression score in myeloid cells.** This table includes the mean expression of the genes *CXCL9, CXCL10, CXCL11* in every sample in 9.MNP_DC3 and in the rest of the myeloid cells. **Related to** Figure 4.

**Supplementary Table 12. Ligand-Receptor list.** This table includes the initial list of ligands and receptors used to find cell-cell interactions (CCIs) between different cell subsets in our data. It was queried from OmniPath and was filtered to a robust set of ligand-receptor pairs (see Methods), with the addition of thyroid specific cell-cell interactions. **Related to** Figure 4.

**Supplementary Table 13. Myeloid-T cell CCIs.** This table includes all the CCIs between T cells and DCs we found in our tissue data. Every row denotes a logan-receptor pair we tested for possible connection between different cell subsets. ‘interactor_DC’ is the gene that is expressed in DC subsets and ’interactor_T’ is the T cell subsets. We tested three different possible connections: in ***Test 1*** (left) first we tested whether there is a connection between specific DC subsets and all CD4+ T cells or all CD8+ T cells; in ***Test 2*** (middle) we tested whether there is a connection between specific DC subsets and specific CD4+ T cell subsets, and in ***Test 3*** (right) we tested whether there is a connection between specific DC subsets and specific CD8+ T cells. For every row we included the expression of the specific interactor (interactor_DC / interactor_T) in the cell subset written in the header, and whether there is a connection for that test, which will be TRUE when both interactors have an expression of >1 in normalized pseudo-bulk. **Related to** Figure 4.

**Supplementary Table 14. Cells in TLS vs nonTLS.** This table includes quantification of different kinds of cell types in RNAish images in regions we defined as TLS and as non-TLS. The tables include the patient for which the slide is done (‘SampleID’ ), the patient’s condition (‘Condition’’), the region type (‘Region’: ’‘TLS’/ ‘non_TLS’), the total number of cells in that regions: CCL19+ cells, CCL19-cells and Tfh cells (‘Observed *’), the total tissue/ region area, the expected number of cells in a given region assuming their distribution is uniform across the tissue (‘expected *’), and the log fold-change between the expected number of cells and the actual number of cells in a region (‘LogFC *’). **Related to** Figure 5**, 6.**

**Supplementary Table 15. TSH vs cell types.** This table includes for every patient (‘Sample’): the ratio between frequency of Tregs and GZMK+ CD8+ T cells (‘CD4T_reg/CD8T_gzmk’), TSH levels (‘TSH (uIU/mL)’), immune infiltration percent (‘Immune infiltration (%)’), frequency of IRTs (‘Frequency 5.E_IRTs (%)’) and the ratio between frequency of immune cells in the tissue and frequency of IRTs (IRTs/Immune_infiltration‘’). **Related to** Figure 5**, 7.**

**Supplementary Table 16. CD4+ T cell TCR sharing and transcriptional similarity.** This table includes the normalized counts of shared TCR clones between pairs of tissue CD4+ T cell subsets (top), the empirical p-values of shared clones across CD4T cell subsets based on 10,000 data shuffles (middle), and transcriptional similarity values between CD4+ T cell subsets based on PAGA algorithm (bottom). **Related to** Figure 5.

**Supplementary Table 17. Top 500 DEGs in all cell subsets 2 diaerent comparisons.** In this table we include the differential expression (DE) analysis done between conditions in all cell subsets in our dataset. We computed DE in two different ways: first, by comparing samples of patients with Hashimoto’s to samples from patients with Grave’s disease (‘contrast’ column value is ‘Hashimotos_vs_Graves’); second, by modeling all the samples along a continuum of immune cell infiltration (‘contrast’ column value is ‘Regression by Immune infiltration’). We included the top 500 differentially expressed genes (DEGs) for every contrast. Every row represents one DEG in one contrast. The columns include the gene’s name (‘Gene’), the lineage and tissue the cell subset is from (‘Lineage’), the cell subset name (‘cluster’), the mean expression of that gene across all samples (‘baseMean’), the log fold-change between conditions or the slope of the regression model (‘log2FoldChange’), the standard error of the fold-change/ slop (‘lfcSE’), the wald test statistic (‘stat’), the p-value (‘pvalue’), adjusted p-value (‘padj’), and absolute log fold-change or absolute slope (‘abslog2FoldChange’).

**Supplementary Table 18. TLS area vs immune infiltration**. This table includes the TLS area as well as the immune infiltration percentage per patient. Every row represents a sample from one patient, every patient has information about their condition (‘condition’) and whether the gland is functioning or not (‘Function’ - may be ‘retained’ or loss of function (‘LoF’)). For every sample we included the total number of TLSs observed in the slide (‘observed_nTLS’), the percent of immune cells infiltrating the gland as depicted by single nuclei sequencing (‘Pct_immune_cells’), the area covered by TLS in the slide (‘Area_covered_by_TLS(um2)’), the total tissue are on the slide (‘Overall_Tissue_Area(um2)’) and the percent of area on the slide that is covered by TLSs (‘Pct_area_covered_by_TLS’). **Related to Figure 7**.

**Supplementary Table 19. Cell enrichment in 3 main region types**. In this table we quantify different cell types in different region types in RNAish slides. For every patient (‘Sample’) we quantify the existence of different cell types (‘Cell_type’: may be CD8+ T cells, epithelial cells that express MHC-II (‘CK+CD74+’), or epithelial cells that don’t express MHC-II (‘CK+CD74-’)) in each of the three main region types we defined (‘Region’: may be ‘Normal’, ‘Dense’ or ‘Sparse’). For every sample and cell types we include the number of cells in that specific region (‘Observed_ncells’), the number of cells we would expected to have in that region if these cells would be uniformly distributed in the tissue (‘Expected_ncells’), the total region size (‘Region size (um^2)’) and the log fold-change between the observed and expected amount of cells in the region (‘log(Observed/Expected_ncells)’). **Related to** Figure 7.

**Supplementary Table 20. Epithelial-T cell CCI.** This table includes all the CCIs between T cells and epithelial cells we found in our tissue data. ‘source’, ‘target and ‘lig-rec’ denote the source gene expressed in the epithelial cell subset and the target gene expressed in the T cell subset. ‘Annotation’ denotes the annotation we gave that CCI. ‘E5_exp’, ‘E1_exp’, ‘E5_scaled_exp’, ‘E1_scaled_exp’ denote the source gene expression levels and scaled expression levels of the epithelial cell subsets 1.E_classical (E1) and 5.E_IRTs (E5). ‘epithelial_pval’ and ‘epithelial_adj_pval’ are the p-value and adjusted p-value of the differential expression between E1 and E5. ‘T_CD8’, ‘T_CD4’, ‘CD8_result’, ‘CD4_result’ are all associated with the main figure 7 plot. ‘T_CD8’, ‘T_CD4’denote the expression of the target gene in either all CD4 T cells or all CD8 T cells. ‘CD8_result’, ‘CD4_result’ is a boolean identifier representing whether this CCI came up in CD8 T cells or CD4 T cells. The rest of the columns in this table associate with supp figure 7 and contain the expression values of the target gene in the CD4 and CD8 T cell subsets in which we looked for CCI with epithelial cells. Any empty cell means that the CCI was not a hit in that cell subset. **Related to** Figure 7.

**Supplementary Table 21. Cell subset frequencies in hubs versus non-hubs.** In this table we quantify different cell types in hubs vs. non-hub regions in RNAish slides. For every patient (‘Sample’) we quantify the existence of different cell types (‘Cell_type’: may be ‘CD8+’ for CD8+ T cells, ‘CD8T+IFNg+’ for CD8+ T cells that express *IFNG*, ‘CK+CD74+’ epithelial cells that express MHC-II, and ‘CK+CD74-’ epithelial cells that don’t express MHC-II. Every row represents a patient and a cell type. For every row we included information about the immune cell infiltration based on single nuclei sequencing (‘ImmunePer’), the condition of the patient (‘Condition’), whether the gland is functioning or not (‘Function’ - may be ‘retained’ or loss of function (‘LoF’)), the total number of cells in hubs for this patient’s slide (‘total_inhub’), the total number of cells in regions that are not hubs for this patient’s slide (‘total_nonhub’), the total number of tiles that are part of a hub for the patient’s slide (‘total_hubtile’), the total number of tiles in the slide (‘nTiles’), the total number of cells of the in the whole slide (‘total_cells’), the density of the cells in the hub tiles (‘Hub_density’), the density of these cells if they were uniformly distributed across the tissue (‘null_density’), the density of the cells in the regions that are non hub tiles (‘nonHub_density’), and the density values normalized and log scaled (‘log_nonHub_normalized’, ‘log_Hub_normalized’). **Related to** Figure 7.

**Supplementary Table 22. Percent of hubs versus immune infiltration**. In this table includes for every sample (‘Sample’) the total number of hub tiles in the RNAish images (‘total_hubtile’) out of the total number of tiles in the slide (‘nTiles’), the percentage of hubs out of the total number of hubs (‘pct_hub’) and the immune cell infiltration based on single nuclei sequencing (‘ImmunePer’). In addition we include information about the condition of every patient (‘condition’) and whether the gland is functioning or not (‘Function’ - may be ‘retained’ or loss of function (‘LoF’)). **Related to** Figure 7.

**Supplementary Table 23. Stimulatory vs checkpoint gene expression.** This table includes the expression values of several stimulatory and checkpoint genes in four different cell subsets. Every row represents a cell subset, the first two columns represent summarizing scores for the checkpoint genes and the stimulatory genes. The rest of the columns represent a different gene each, with an annotation for each whether it is checkpoint or stimulatory. **Related to** Figure 7.

**Supplementary Table 24. Dysfunction gene expression in T cells versus checkpoint gene expression in IRTs.** This table includes for every sample (‘Sample’), the patient condition (‘Condition’) and thyroid function status (‘Thyroid function’) as well as the gene expression of a list of checkpoint genes in 5.E_IRTs (‘checkpoint_score’) and the gene expression of dysfunctional genes in GZMK^+^ and T_rm_-like CD8^+^ T cells (‘Dysfunctional_score’). **Related to** Figure 7.

**Supplementary Table 25. TSH versus IRTs and Immune Infiltration ratio.** Every row in this table represents one Hashimoto’s patient. This table includes the values of TSH (‘TSH (uIU/mL)’), the immune cell infiltration frequency as measured by single nuclei data (‘Immune infiltration (%)’), the frequency of the cell subset 5.E_IRTs (‘IRTs frequency (%)’), and the ratio between these two frequencies (‘IRTs/perc_imm’). **Related to** Figure 7.

**Supplementary Table 26. Software package versions.** This table details all the packages used for the data analysis of this study. The left table includes the Python packages including the repository’s name (‘Repo name’), the repository’s version (‘Version’), the repository’s build (‘Build’), and the channel from which it was obtained into the conda environment (‘Channel’). The right matrix includes all the R packages used and includes the name of the package and the version used. **Related to all figures.**

**Supplementary Table 27. Surface antibodies (CITE-seq) and hashtags.** This table includes information about the antibodies and hashtags used for CITEseq. Every row represents a different antibody and includes the DNA ID (‘Cite-seq DNA ID’), the antibody description (‘Antibody description’) the clone (‘clone’), the antibody oligonucleotide conjugate (‘barcode’), the protein the antibody attaches to (‘Protein name’), the gene that codes this protein (‘Gene Symbol’), and the category (‘Category’ - can be either ‘CITE’ for antibodies specific proteins, or ‘hashtags’ for *B2M* antibodies used for cell hashing). **Related to** Figures 3-6.

## REFERENCES

1. Miller, F.W. (2023). The increasing prevalence of autoimmunity and autoimmune diseases: an urgent call to action for improved understanding, diagnosis, treatment, and prevention. Curr. Opin. Immunol. 80, 102266. 10.1016/j.coi.2022.102266.

2. Conrad, N., Misra, S., Verbakel, J.Y., Verbeke, G., Molenberghs, G., Taylor, P.N., Mason, J., Sattar, N., McMurray, J.J.V., McInnes, I.B., et al. (2023). Incidence, prevalence, and co-occurrence of autoimmune disorders over time and by age, sex, and socioeconomic status: a population-based cohort study of 22 million individuals in the UK. Lancet Lond. Engl. 401, 1878–1890. 10.1016/S0140-6736(23)00457-9.

3. Hollowell, J.G., Staehling, N.W., Flanders, W.D., Hannon, W.H., Gunter, E.W., Spencer, C.A., and Braverman, L.E. (2002). Serum TSH, T(4), and thyroid antibodies in the United States population (1988 to 1994): National Health and Nutrition Examination Survey (NHANES III). J. Clin. Endocrinol. Metab. 87, 489–499. 10.1210/jcem.87.2.8182.

4. Yoshida, H., Amino, N., Yagawa, K., Uemura, K., Satoh, M., Miyai, K., and Kumahara, Y. (1978). Association of serum antithyroid antibodies with lymphocytic infiltration of the thyroid gland: studies of seventy autopsied cases. J. Clin. Endocrinol. Metab. 46, 859–862. 10.1210/jcem-46-6-859.

5. Vanderpump, M.P., Tunbridge, W.M., French, J.M., Appleton, D., Bates, D., Clark, F., Grimley Evans, J., Hasan, D.M., Rodgers, H., and Tunbridge, F. (1995). The incidence of thyroid disorders in the community: a twenty-year follow-up of the Whickham Survey. Clin. Endocrinol. (Oxf.) 43, 55–68. 10.1111/j.1365-2265.1995.tb01894.x.

6. Dayan, C.M., and Daniels, G.H. (1996). Chronic autoimmune thyroiditis. N. Engl. J. Med. 335, 99–107. 10.1056/NEJM199607113350206.

7. Smith, T.J., and Hegedüs, L. (2016). Graves’ Disease. N. Engl. J. Med. 375, 1552–1565. 10.1056/NEJMra1510030.

8. Stoeckius, M., Hafemeister, C., Stephenson, W., Houck-Loomis, B., Chattopadhyay, P.K., Swerdlow, H., Satija, R., and Smibert, P. (2017). Simultaneous epitope and transcriptome measurement in single cells. Nat. Methods 14, 865–868. 10.1038/nmeth.4380.

9. Hadlow, N.C., Rothacker, K.M., Wardrop, R., Brown, S.J., Lim, E.M., and Walsh, J.P. (2013). The relationship between TSH and free T₄ in a large population is complex and nonlinear and differs by age and sex. J. Clin. Endocrinol. Metab. 98, 2936–2943. 10.1210/jc.2012-4223.

10. Taylor, P.N., Medici, M.M., Hubalewska-Dydejczyk, A., and Boelaert, K. (2024). Hypothyroidism. Lancet Lond. Engl. 404, 1347–1364. 10.1016/S0140-6736(24)01614-3.

11. Roche, P.A., and Furuta, K. (2015). The ins and outs of MHC class II-mediated antigen processing and presentation. Nat. Rev. Immunol. 15, 203–216. 10.1038/nri3818.

12. Kambayashi, T., and Laufer, T.M. (2014). Atypical MHC class II-expressing antigen-presenting cells: can anything replace a dendritic cell? Nat. Rev. Immunol. 14, 719–730. 10.1038/nri3754.

13. Wang, S.H., Van Antwerp, M., Kuick, R., Gauger, P.G., Doherty, G.M., Fan, Y.Y., and Baker, J.R. (2007). Microarray analysis of cytokine activation of apoptosis pathways in the thyroid. Endocrinology 148, 4844–4852. 10.1210/en.2007-0126.

14. Cooper, M.A., Fehniger, T.A., and Caligiuri, M.A. (2001). The biology of human natural killer-cell subsets. Trends Immunol. 22, 633–640. 10.1016/S1471-4906(01)02060-9.

15. Li, H., Leun, A.M. van der, Yofe, I., Lubling, Y., Gelbard-Solodkin, D., Akkooi, A.C.J. van, Braber, M. van den, Rozeman, E.A., Haanen, J.B.A.G., Blank, C.U., et al. (2019). Dysfunctional CD8 T Cells Form a Proliferative, Dynamically Regulated Compartment within Human Melanoma. Cell 176, 775–789.e18. 10.1016/j.cell.2018.11.043.

16. Chiffelle, J., Genolet, R., Perez, M.A., Coukos, G., Zoete, V., and Harari, A. (2020). T-cell repertoire analysis and metrics of diversity and clonality. Curr. Opin. Biotechnol. 65, 284–295. 10.1016/j.copbio.2020.07.010.

17. Wu, T.D., Madireddi, S., de Almeida, P.E., Banchereau, R., Chen, Y.-J.J., Chitre, A.S., Chiang, E.Y., Iftikhar, H., O’Gorman, W.E., Au-Yeung, A., et al. (2020). Peripheral T cell expansion predicts tumour infiltration and clinical response. Nature 579, 274–278. 10.1038/s41586-020-2056-8.

18. Cabeza-Cabrerizo, M., Cardoso, A., Minutti, C.M., Pereira da Costa, M., and Reis e Sousa, C. (2021). Dendritic Cells Revisited. Annu. Rev. Immunol. 39, 131–166. 10.1146/annurev-immunol-061020-053707.

19. Villani, A.-C., Satija, R., Reynolds, G., Sarkizova, S., Shekhar, K., Fletcher, J., Griesbeck, M., Butler, A., Zheng, S., Lazo, S., et al. (2017). Single-cell RNA-seq reveals new types of human blood dendritic cells, monocytes, and progenitors. Science 356, eaah4573. 10.1126/science.aah4573.

20. Bill, R., Wirapati, P., Messemaker, M., Roh, W., Zitti, B., Duval, F., Kiss, M., Park, J.C., Saal, T.M., Hoelzl, J., et al. (2023). CXCL9:SPP1 macrophage polarity identifies a network of cellular programs that control human cancers. Science 381, 515–524. 10.1126/science.ade2292.

21. Pascual-García, M., Bonfill-Teixidor, E., Planas-Rigol, E., Rubio-Perez, C., Iurlaro, R., Arias, A., Cuartas, I., Sala-Hojman, A., Escudero, L., Martínez-Ricarte, F., et al. (2019). LIF regulates CXCL9 in tumor-associated macrophages and prevents CD8+ T cell tumor-infiltration impairing anti-PD1 therapy. Nat. Commun. 10, 2416. 10.1038/s41467-019-10369-9.

22. Chow, M.T., Ozga, A.J., Servis, R.L., Frederick, D.T., Lo, J.A., Fisher, D.E., Freeman, G.J., Boland, G.M., and Luster, A.D. (2019). Intratumoral Activity of the CXCR3 Chemokine System Is Required for the Efficacy of Anti-PD-1 Therapy. Immunity 50, 1498–1512.e5. 10.1016/j.immuni.2019.04.010.

23. Pipi, E., Nayar, S., Gardner, D.H., Colafrancesco, S., Smith, C., and Barone, F. (2018). Tertiary Lymphoid Structures: Autoimmunity Goes Local. Front. Immunol. 9. 10.3389/FIMMU.2018.01952.

24. Pelka, K., Hofree, M., Chen, J.H., Sarkizova, S., Pirl, J.D., Jorgji, V., Bejnood, A., Dionne, D., Ge, W.H., Xu, K.H., et al. (2021). Spatially organized multicellular immune hubs in human colorectal cancer. Cell 184, 4734–4752.e20. 10.1016/J.CELL.2021.08.003.

25. Chen, J.H., Nieman, L.T., Spurrell, M., Jorgji, V., Elmelech, L., Richieri, P., Xu, K.H., Madhu, R., Parikh, M., Zamora, I., et al. (2024). Human lung cancer harbors spatially organized stem-immunity hubs associated with response to immunotherapy. Nat. Immunol. 25, 644–658. 10.1038/s41590-024-01792-2.

26. Hamilton, F., Black, M., Farquharson, M.A., Stewart, C., and Foulis, A.K. (1991). Spatial correlation between thyroid epithelial cells expressing class II MHC molecules and interferon-gamma-containing lymphocytes in human thyroid autoimmune disease. Clin. Exp. Immunol. 83, 64. 10.1111/J.1365-2249.1991.TB05589.X.

27. Bottazzo, G., Hanafusa, T., Pujol-Borrell, R., and Feldmann, M. (1983). ROLE OF ABERRANT HLA-DR EXPRESSION AND ANTIGEN PRESENTATION IN INDUCTION OF ENDOCRINE AUTOIMMUNITY. The Lancet 322, 1115–1119. 10.1016/S0140-6736(83)90629-3.

28. Herren, R., and Geva-Zatorsky, N. (2024). Spatial features of skip lesions in Crohn’s disease. Trends Immunol. 45, 470–481. 10.1016/j.it.2024.04.011.

29. Willcox, A., Richardson, S.J., Bone, A.J., Foulis, A.K., and Morgan, N.G. (2009). Analysis of islet inflammation in human type 1 diabetes. Clin. Exp. Immunol. 155, 173–181. 10.1111/j.1365-2249.2008.03860.x.

30. Nestle, F.O., Kaplan, D.H., and Barker, J. (2009). Psoriasis. N. Engl. J. Med. 361, 496–509. 10.1056/NEJMra0804595.

31. Nayar, S., Campos, J., Smith, C.G., Iannizzotto, V., Gardner, D.H., Mourcin, F., Roulois, D., Turner, J., Sylvestre, M., Asam, S., et al. (2019). Immunofibroblasts are pivotal drivers of tertiary lymphoid structure formation and local pathology. Proc. Natl. Acad. Sci. 116, 13490–13497. 10.1073/pnas.1905301116.

32. Korsunsky, I., Wei, K., Pohin, M., Kim, E.Y., Barone, F., Major, T., Taylor, E., Ravindran, R., Kemble, S., Watts, G.F.M., et al. (2022). Cross-tissue, single-cell stromal atlas identifies shared pathological fibroblast phenotypes in four chronic inflammatory diseases. Med N. Y. N 3, 481–518.e14. 10.1016/j.medj.2022.05.002.

33. Sureshchandra, S., Henderson, J., Levendosky, E., Bhattacharyya, S., Kastenschmidt, J.M., Sorn, A.M., Mitul, M.T., Benchorin, A., Batucal, K., Daugherty, A., et al. (2024). Tissue determinants of the human T cell receptor repertoire. bioRxiv, 2024.08.17.608295. 10.1101/2024.08.17.608295.

34. Laydon, D.J., Bangham, C.R.M., and Asquith, B. (2015). Estimating T-cell repertoire diversity: limitations of classical estimators and a new approach. Philos. Trans. R. Soc. B Biol. Sci. 370, 20140291. 10.1098/rstb.2014.0291.

35. Chaker, L., Cooper, D.S., Walsh, J.P., and Peeters, R.P. (2024). Hyperthyroidism. Lancet Lond. Engl. 403, 768–780. 10.1016/S0140-6736(23)02016-0.

36. Chaker, L., Razvi, S., Bensenor, I.M., Azizi, F., Pearce, E.N., and Peeters, R.P. (2022). Hypothyroidism. Nat. Rev. Dis. Primer 8, 30. 10.1038/s41572-022-00357-7.

37. Lanzolla, G., Marinò, M., and Menconi, F. (2024). Graves disease: latest understanding of pathogenesis and treatment options. Nat. Rev. Endocrinol. 20, 647–660. 10.1038/s41574-024-01016-5.

38. Tomer, Y., and Davies, T.F. (2003). Searching for the autoimmune thyroid disease susceptibility genes: from gene mapping to gene function. Endocr. Rev. 24, 694–717. 10.1210/er.2002-0030.

39. Lee, H.J., Li, C.W., Hammerstad, S.S., Stefan, M., and Tomer, Y. (2015). Immunogenetics of autoimmune thyroid diseases: A comprehensive review. J. Autoimmun. 64, 82–90. 10.1016/j.jaut.2015.07.009.

40. Kraiem, Z., Baron, E., Kahana, L., Sadeh, O., and Sheinfeld, M. (1992). Changes in stimulating and blocking TSH receptor antibodies in a patient undergoing three cycles of transition from hypo to hyper-thyroidism and back to hypothyroidism. Clin. Endocrinol. (Oxf.) 36, 211–214. 10.1111/j.1365-2265.1992.tb00960.x.

41. Tamai, H., Kasagi, K., Takaichi, Y., Takamatsu, J., Komaki, G., Matsubayashi, S., Konishi, J., Kuma, K., Kumagai, L.F., and Nagataki, S. (1989). Development of spontaneous hypothyroidism in patients with Graves’ disease treated with antithyroidal drugs: clinical, immunological, and histological findings in 26 patients. J. Clin. Endocrinol. Metab. 69, 49–53. 10.1210/jcem-69-1-49.

42. Tamai, H., Ohsako, N., Takeno, K., Fukino, O., Takahashi, H., Kuma, K., Kumagai, L.F., and Nagataki, S. (1980). Changes in thyroid function in euthyroid subjects with a family history of Graves’ disease: a follow-up study of 69 patients. J. Clin. Endocrinol. Metab. 51, 1123–1127. 10.1210/jcem-51-5-1123.

43. Brix, T.H., and Hegedüs, L. (2012). Twin studies as a model for exploring the aetiology of autoimmune thyroid disease. Clin. Endocrinol. (Oxf.) 76, 457–464. 10.1111/j.1365-2265.2011.04318.x.

44. Lechner, M.G., Zhou, Z., Hoang, A.T., Huang, N., Ortega, J., Scott, L.N., Chen, H.-C., Patel, A.Y., Yakhshi-Tafti, R., Kim, K., et al. (2023). Clonally expanded, thyrotoxic effector CD8+ T cells driven by IL-21 contribute to checkpoint inhibitor thyroiditis. Sci. Transl. Med. 15. 10.1126/SCITRANSLMED.ADG0675.

45. Drokhlyansky, E., Smillie, C.S., Van Wittenberghe, N., Ericsson, M., Griffin, G.K., Eraslan, G., Dionne, D., Cuoco, M.S., Goder-Reiser, M.N., Sharova, T., et al. (2020). The Human and Mouse Enteric Nervous System at Single-Cell Resolution. Cell 182, 1606–1622.e23. 10.1016/j.cell.2020.08.003.

46. Zheng, G.X.Y., Terry, J.M., Belgrader, P., Ryvkin, P., Bent, Z.W., Wilson, R., Ziraldo, S.B., Wheeler, T.D., McDermott, G.P., Zhu, J., et al. (2017). Massively parallel digital transcriptional profiling of single cells. Nat. Commun. 8, 1–12. 10.1038/ncomms14049.

47. Li, B., Gould, J., Yang, Y., Sarkizova, S., Tabaka, M., Ashenberg, O., Rosen, Y., Slyper, M., Kowalczyk, M.S., Villani, A.C., et al. (2020). Cumulus provides cloud-based data analysis for large-scale single-cell and single-nucleus RNA-seq. Nat. Methods 17, 793–798. 10.1038/s41592-020-0905-x.

48. Heaton, H., Talman, A.M., Knights, A., Imaz, M., Gaffney, D.J., Durbin, R., Hemberg, M., and Lawniczak, M.K.N. (2020). Souporcell: robust clustering of single-cell RNA-seq data by genotype without reference genotypes. Nat. Methods 2020 176 17, 615–620. 10.1038/s41592-020-0820-1.

49. Butler, A., Hoffman, P., Smibert, P., Papalexi, E., and Satija, R. (2018). Integrating single-cell transcriptomic data across different conditions, technologies, and species. Nat. Biotechnol. 36, 411–420. 10.1038/nbt.4096.

50. Korsunsky, I., Millard, N., Fan, J., Slowikowski, K., Zhang, F., Wei, K., Baglaenko, Y., Brenner, M., Loh, P., and Raychaudhuri, S. (2019). Fast, sensitive and accurate integration of single-cell data with Harmony. Nat. Methods 16, 1289–1296. 10.1038/s41592-019-0619-0.

51. Ritchie, M.E., Phipson, B., Wu, D., Hu, Y., Law, C.W., Shi, W., and Smyth, G.K. (2015). limma powers differential expression analyses for RNA-sequencing and microarray studies. Nucleic Acids Res. 43, e47–e47. 10.1093/nar/gkv007.

52. Slowikowski, Kamil (2023). cellguide: Navigate single-cell RNA-seq datasets in your web browser. Version 10.5281/ZENODO.8144195.

53. Türei, D., Valdeolivas, A., Gul, L., Palacio-Escat, N., Klein, M., Ivanova, O., Ölbei, M., Gabor, A., Theis, F., Modos, D., et al. (2021). Integrated intra- and intercellular signaling knowledge for multicellular omics analysis. Mol. Syst. Biol. 17, e9923. 10.15252/MSB.20209923.

54. Love, M.I., Huber, W., and Anders, S. (2014). Moderated estimation of fold change and dispersion for RNA-seq data with DESeq2. Genome Biol. 15, 550. 10.1186/s13059-014-0550-8.

